# CD44-CD206 axis drives dendritic cell loss and immunosuppression in immaturely glycosylated bladder tumours

**DOI:** 10.64898/2026.10.25.761623

**Authors:** Andreia Miranda, Dylan Ferreira, Rui Freitas, Andreia Peixoto, Andreia Brandão, Michael de Kok, Eduardo Ferreira, Marta Relvas-Santos, Camila Lourenço, Magali Grange, Flávia Castro, Luís Pedro Afonso, Carlos Palmeira, Lúcio Lara Santos, Maria José Oliveira, Hervé Luche, Yvette van Kooyk, José Alexandre Ferreira

## Abstract

Antigen-presenting-cell dysfunction drives immune exclusion, but how tumour glycans disable these cells remains incompletely defined. Immature *O*-glycosylation, marked by Tn and sialyl-Tn antigens, drives myeloid dysfunction in bladder cancer. Transcriptomics and human tumour profiling linked this state to reduced representation of antigen-presenting myeloid cells and attenuated lymphoid-effector programmes. In glycoengineered models, immature *O*-glycans promoted immunoregulatory macrophage phenotypes and contact-dependent death of monocyte-derived dendritic cells (moDCs), while surviving cells showed impaired maturation and T-cell activation. Tumour-cell CD44 standard isoform was required for DC death, which was accompanied by inflammasome-associated proteins and gasdermin D cleavage, consistent with pyroptosis. Receptor-discovery approaches converged on CD206, which bound immaturely glycosylated CD44. CD206 blockade restored DC viability. In immunocompetent mice, immature glycosylation accelerated tumour growth and depleted conventional DCs, whereas CD206 blockade restrained tumour progression and restored local and nodal myeloid and lymphoid immunity. These findings identify CD44-CD206 as a targetable glycoimmune checkpoint that promotes immune exclusion.

## Main

Immune exclusion is a hallmark of aggressive solid tumours and a major determinant of resistance to immunotherapy^1,2^. Bladder urothelial carcinomas (BLCA) exhibit diverse immune landscapes, ranging from T-cell-inflamed, immune-excluded and immune-desert states that are associated with distinct clinical outcome and response to immune-checkpoint blockade^3,4^. Defective antigen presentation, impaired T-cell activation and T-cell exhaustion are well established, but the tumour-cell-intrinsic programmes that actively generate these immune states remain incompletely understood^5^. In particular, it is unclear how the tumour-cell surface instructs myeloid dysfunction in ways that suppress downstream lymphoid immunity^6^.

The tumour glycocalyx is a critical interface for immune recognition and signalling^7^. Aberrant *O*-glycosylation, including accumulation of the truncated Tn and sialyl-Tn (sTn) antigens, is a recurrent feature of malignancy associated with aggressive disease and immune dysregulation^8,9^. In BLCA, both antigens are associated with high-risk clinical behaviour^10,11^, and experimental models implicate immature *O*-glycosylation in invasion and stress resistance^12,13^. We previously identified the hyaluronic acid receptor CD44 standard isoform (CD44s) as a major scaffold for truncated *O*-glycans in BLCA and showed that this modification promotes epithelial-mesenchymal plasticity, which may foster progression^14,15^. Whether truncated *O-*glycans tumour glycocode also directly contributes to immune exclusion remains unresolved.

Tumour-associated glycans can modulate myeloid-cell function through lectin receptors that sense GalNAc- and sialic-acid-containing motifs^7,16^. Tn can engage the C-type lectin MGL^17^, whereas sialylated glycans can signal through inhibitory Siglec receptors, including Siglec-7, Siglec-9 and Siglec-15, to restrain inflammatory and antigen-presenting programmes^18,19^. These interactions support a model in which aberrant tumour glycosylation promotes myeloid tolerance and indirectly drives suppression of T-cell immunity. Together, these observations raise the possibility that immature tumour *O*-glycans can directly act on antigen-presenting cells (APCs), impairing their survival and functional competence.

Here we identify immature *O*-glycosylation as a tumour-cell-intrinsic programme that disrupts myeloid-lymphoid crosstalk in BLCA. Using glycoengineered tumour-cell models, human specimens and an immunocompetent syngeneic mouse model, and advanced molecular and spatial imaging techniques, we show that truncated *O*-glycans promote immunoregulatory macrophage states and contact-dependent moDC loss. Surviving DCs exhibit impaired maturation and a reduced capacity to present antigens and to support T-cell activation. Tumour-cell CD44s, carrying truncated *O*-glycans, is required for these effects and bind to the mannose receptor CD206 on moDCs surface. CD206 blockade restrains the growth of tumours bearing immature *O*-glycans and partially reverses the associated suppression of antitumour immunity in mouse tumours and tumour-draining lymph nodes. Together, these findings unveil an unrecognized CD44s-CD206 glycoimmune axis that links tumour-cell glycosylation to DC dysfunction and immune exclusion.

## Results

### Immature glycosylation defines an immunosuppressive tumour state in BLCA

To determine whether *O*-glycosylation gene expression is associated with the immune landscape in BLCA, we analysed 407 muscle-invasive bladder cancer (MIBC) TCGA transcriptomes (**Supplementary Table S1**) using a curated panel of *O*-glycan biosynthesis genes and immune-cell markers. Expression of the *O*-glycan extension enzymes *C1GALT1* and *GCNT1* correlated positively with aggregate immune-marker expression, whereas the Tn-sialyltransferase *ST6GALNAC1* showed the opposite relationship (**Fig. 1a,b**). Interestingly, expression of *C1GALT1C1*, which encodes an essential chaperone for C1GalT1 activity, was not significantly associated with immune-marker expression (**Fig. 1b**). Also, *ST3GAL1* and *ST6GALNAC2*, which participate in related core 1 sialylation pathways, and the core 3 synthase *B3GNT6* showed no comparable association (**Supplementary Fig. S1**). Beyond these gene-level correlations, bulk transcriptomic analysis showed reduced expression of multiple immune-cell-associated genes both in *C1GALT1*-low tumours and, particularly, in tumours combining low *C1GALT1* with high *ST6GALNAC1* expression (**Supplementary Fig. S2a-c**). Together, these findings link impaired *O*-glycan elongation, particularly when coupled to increased Tn sialylation, to broad attenuation of immune programmes in BLCA.

**Fig. 1.**
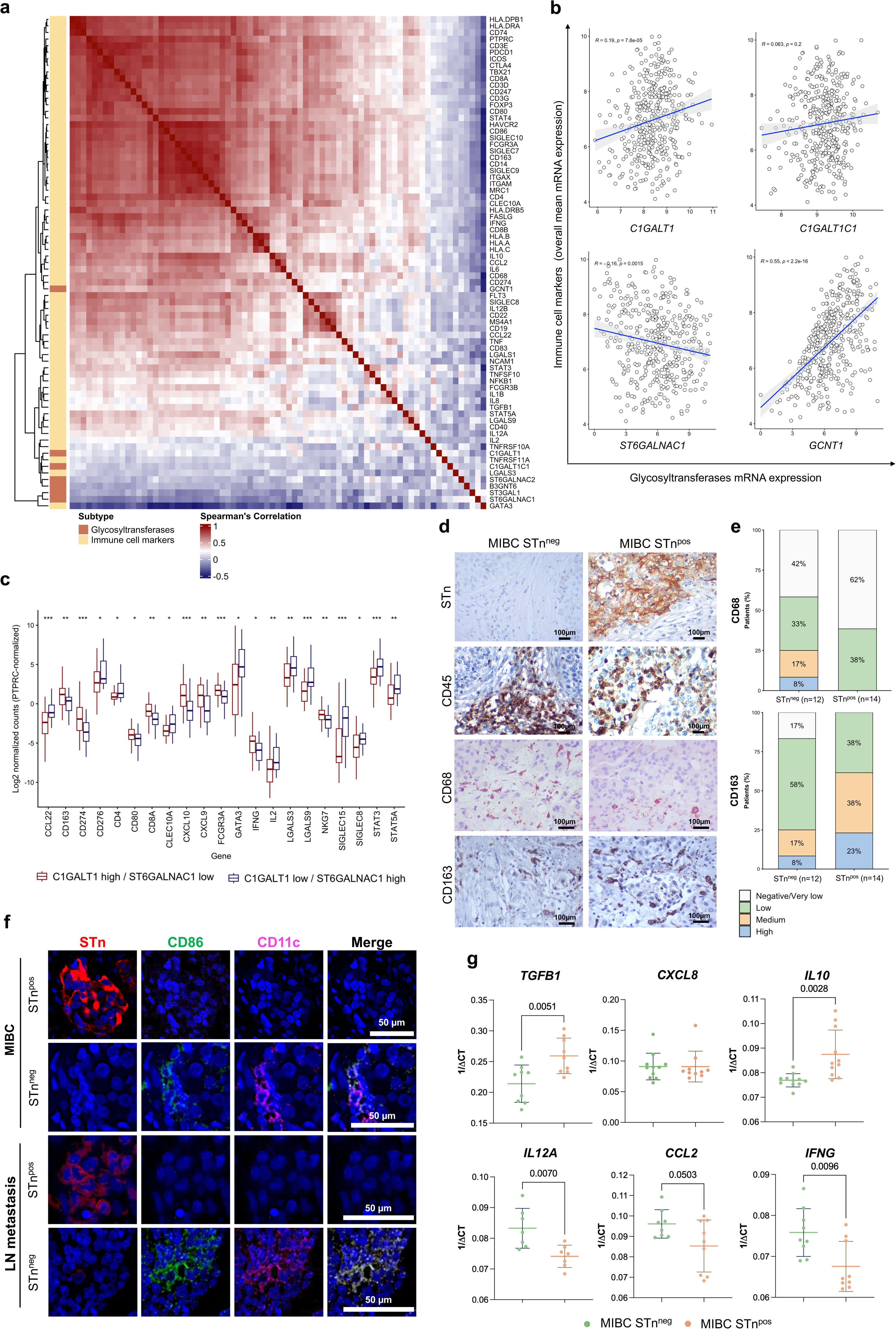
Immature *O*-glycosylation associates with immune suppression in bladder cancer. **a** Hierarchically clustered Spearman correlation matrix of selected *O*-glycosylation and immune-cell-associated genes in samples from muscle-invasive bladder cancers (MIBC; n=407) from TCGA-BLCA. **b** Glycogenes controlling *O*-glycan maturation show distinct associations with immune-marker expression. *C1GALT1* and *GCNT1*, which promote *O*-glycan elongation, correlated positively with mean immune-marker expression, whereas the Tn-sialyltransferase *ST6GALNAC1* correlated inversely; *C1GALT1C1* showed no significant association. Dots denote tumours; lines and shading indicate linear fits and 95% confidence intervals. Spearman ρ and *P* values are shown. **c** The *C1GALT1*-low/*ST6GALNAC1*-high profile, consistent with enhanced sTn formation, is associated with reduced cytotoxic/type 1 and increased immunoregulatory expression. *PTPRC*-normalized immune-gene expression in reciprocal *C1GALT1*/*ST6GALNAC1* groups. Box plots show medians, interquartile ranges and 1.5×IQR whiskers. Two-sided Wilcoxon rank-sum tests; \**P* < 0.05, \*\**P* < 0.01 and \*\*\**P* < 0.001. **d** sTn-expressing MIBC shows predominantly intratumoral CD45+ infiltration and a CD68-low/CD163-high macrophage-associated profile. Representative immunohistochemistry (**d**) and CD68/CD163 staining-score distributions (**e**) in sTn-negative (n=12) and sTn-expressing (n=14) MIBC. Scale bars, 100 μm. **f** sTn expression is associated with marked loss of DC-associated staining in primary MIBC and lymph-node metastases. Representative multiplex immunofluorescence showing near-absent CD11c/CD86 staining in sTn^pos^ cases. sTn, red; CD86, green; CD11c, magenta; DAPI, blue. Scale bars, 50 μm. **g** sTn-expressing MIBC displays a cytokine gene expression profile consistent with reduced type 1 immunity and enhanced immunoregulation. Relative to sTn-negative, sTn-expressing tumours expressed higher levels of *TGFB1* and *IL10* and lower levels of *IL12A*, *IFNG* and *CCL2*, whereas *CXCL8* remained unchanged. *P* values were determined using two-sided Mann-Whitney *U*-tests.

To account for differences in leukocyte abundance, we normalized immune-gene expression to *PTPRC* (CD45), which was reduced in *C1GALT1*-low/*ST6GALNAC1*-high tumours (**Fig. 1c**). After normalization, these tumours showed selective remodeling of myeloid-associated genes, with lower relative expression of *CD163*, *FCGR3A* and *CD80*, but higher relative expression of *CLEC10A*, *SIGLEC8* and *SIGLEC15* (**Fig. 1c**). They also showed lower relative expression of cytotoxic and type 1 immune genes, including *CD8A*, *CXCL9*, *CXCL10*, *IFNG* and *NKG7*, and higher relative expression of immunoregulatory genes, including *CCL22*, *CD276*, *GATA3*, *LGALS3*, *LGALS9*, *STAT3* and *STAT5A* (**Fig. 1c**). Overall, the immune phenotype associated with a Tn/sTn-prone glycosylation programme was characterized by myeloid remodelling and a shift from cytotoxic and type 1 immunity towards an immunoregulatory state.

We next analysed another cohort comprising 26 therapy-naive MIBC patient specimens, matching the disease context represented by TCGA-BLCA. Approximately 54% (14 of the 26) of tumours expressed STn as the predominant immature *O*-glycan epitope, with diffuse staining across the tumour area in all positive cases. Tn co-expression was also observed in a subset of STn-positive tumours. Immunohistochemistry revealed distinct spatial patterns of immune infiltration: immune cells were predominantly intratumoral in sTn-positive MIBC but remained largely confined to the tumour periphery in sTn-negative tumours (**Fig. 1d**). This redistribution may reflect reduced cell–cell adhesion and increased tissue plasticity previously associated with immature *O*-glycosylation^13,16^, although altered chemotactic signalling, as suggested by the TCGA analysis (**Fig. 1c**), may also contribute.STn-positive tumours also showed lower CD68 and higher CD163 staining, consistent with enrichment of immunoregulatory macrophages (**Fig. 1d, e**). Despite the presence of intratumoral CD45-positive cells and CD163-positive macrophages, multiplex immunofluorescence revealed a near-complete absence of CD11c and CD86 staining in STn-positive primary tumours (**Fig. 1f**). This pattern was recapitulated in STn-positive proximal lymph-node metastases, suggesting marked depletion of CD11c+ myeloid cells (**Fig. 1f**). STn-positive tumours further showed higher levels of the immune suppressor cytokines *TGFB1* and *IL10* and lower levels of the pro-inflammatory and chemotactic mediators *IL12A*, *IFNG* and *CCL2* mRNA expression (**Fig. 1g**). Together, these tissue-level findings recapitulate and extend the TCGA-derived immunesupressive programmes. STn expression in human MIBC was associated with spatial reorganisation of immune infiltrates, CD163-skewed macrophage remodelling, marked CD11c+ DC depletion, and broader TME reprogramming from inflammatory and type 1 immunity toward a tolerogenic, immunoregulatory state.

### Immature glycans impair myeloid-cell function and T cell priming

To determine how immature tumour *O*-glycosylation affects myeloid-cell function, we used matched glycoengineered human T24 bladder cancer models. WT cells displayed only vestigial Tn and sTn expression (**Supplementary Fig. S3a**). *C1GALT1* knockout induced homogeneous Tn expression with minor sTn (approximately 7:1 Tn:sTn), whereas additional *ST6GALNAC1* expression shifted this ratio to approximately 1:1 (**Supplementary Fig. S3b**). Both glycoengineered models showed increased invasive capacity compared with WT cells, with the strongest phenotype observed after *ST6GALNAC1* expression (**Supplementary Fig. S3c**).

Co-culture with Tn/sTn-expressing tumour cells polarized human monocyte-derived macrophages towards an immunoregulatory phenotype, characterized by increased expression of CD206 and IL-10 and reduced expression of the co-stimulatory receptor CD86, without comparable changes in the major histocompatibility class II (MHC-II) molecule HLA-DR or the scavenger receptor CD163 (**Fig. 2a-d and Supplementary Fig. S4**). This response was accompanied by a modest increase in macrophage death in co-cultures with unpolarized macrophages (**Supplementary Fig. S5a**). In M1-polarized macrophages, Tn/sTn-expressing tumour cells reduced CD86 and HLA-DR while increasing CD206, CD163, and IL-10 expression, without significantly affecting M1-like macrophage viability (**Fig. 2e-g and Supplementary Fig. S5b**), consistent with reprogramming from a pro-inflammatory to an anti-inflammatory macrophage state. Tn/STn-expressing T24 cells were also less susceptible to killing by either unpolarized or M1-polarized macrophages (**Supplementary Fig. S5c**). Thus, immature *O*-glycans promote anti-inflammatory macrophage polarization while protecting tumour cells from macrophage-mediated killing in a CD47-independent mechanism, as the expression of this anti-phagocytic signal remained unchanged (**Supplementary Fig. S6**).

**Fig. 2.**
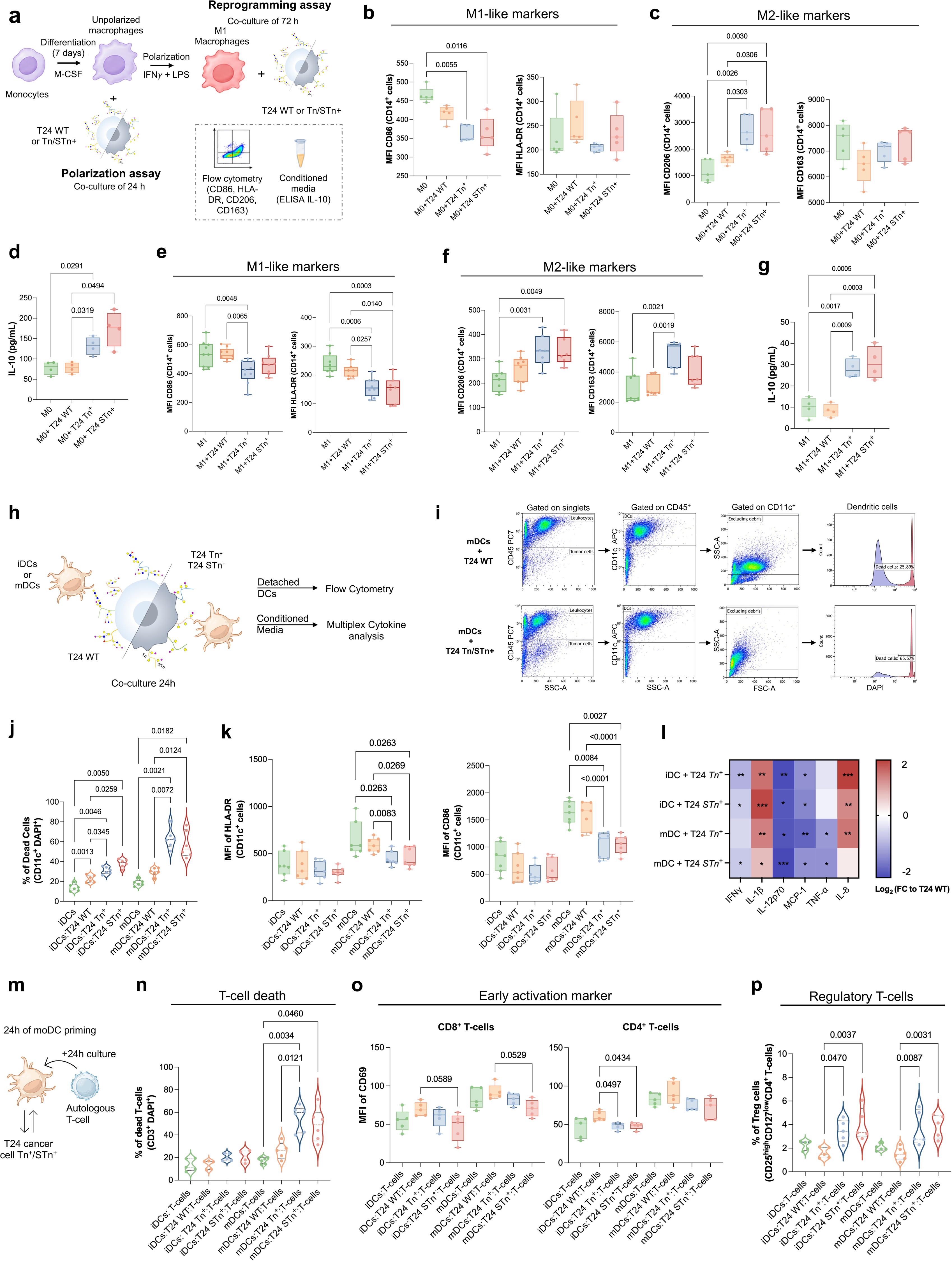
Immature tumour *O*-glycans promote immunoregulatory macrophage states and disrupt DC-T-cell immunity. **a** Design of 24 h co-cultures of unpolarized macrophages and 72 h co-cultures of M1-polarized macrophages with WT, Tn- or sTn-expressing T24 cells. **b-d** Tn/sTn-expressing cells polarize unpolarized macrophages towards an immunoregulatory phenotype. Flow cytometry showed reduced CD86 and increased CD206, without significant changes in HLA-DR or CD163 expression (**b,c**); ELISA showed increased IL-10 secretion (**d**). **e-g** Tn/sTn-expressing cells reprogramme M1-polarized macrophages towards an immunoregulatory phenotype, with reduced M1-associated and increased M2-associated marker expression (**e,f**) and increased IL-10 secretion (**g**). **h** Design of 24 h co-cultures of iDCs or mDCs with WT, Tn- or sTn-expressing T24 cells for analysis of DC viability, activation and cytokine secretion. **i-l** Immature tumour *O*-glycans induce DC death and impair activation. Representative CD45^+^CD11c^+^ gating and DAPI-based viability analysis (**i**), with quantification of increased DC death (**j**) and reduced HLA-DR and CD86 expression (**k**), particularly in mDCs. **l** Immature *O*-glycans alter cytokine production by iDCs and mDCs. Heatmap shows log_2_ fold changes versus T24 WT conditions. **m** Design of autologous T-cell assays. iDCs or mDCs were exposed to WT, Tn- or sTn-expressing T24 cells for 24 h, followed by 24 h autologous T-cell co-culture. **n-p** Tumour cells-exposed DCs compromise T-cell responses, increasing T-cell death (**n**), attenuating CD69 induction on CD8^+^ and CD4^+^ T cells (**o**) and increasing CD4^+^CD25^high^CD127^low^ regulatory T cells (**p**). One-way ANOVA followed by Dunnett’s multiple-comparisons test. Exact *P* values are shown.

We next examined the impact of cancer cell Tn and sTn expression in moDCs viability and activity (**Fig. 2h**). Notably, Tn/sTn-expressing T24 cells induced marked death of both immature moDCs (iDCs) and mature moDCs (mDCs), with an approximately threefold increase in mDC death relatively to WT co-cultures (**Fig. 2i,j**). FAS (CD95) expression remained unchanged (**Supplementary Fig. S7**), suggesting moDC loss was not associated with increased FAS abundance. In contrast to iDCs, surviving mDCs showed reduced HLA-DR and CD86 expression (**Fig. 2k**). Their cytokine profiling further revealed increased IL-1β and IL-8 and reduced IL-12p70, MCP-1 (CCL2), IFN-γ and TNF-α expression (**Fig. 2l**). This combination indicated reduction of pro-inflammatory activity and of moDC activation and prompted investigation of the underlying cell-death programme.

Conditioned medium from glycoengineered T24 cells expressing Tn- and sTn glycoproteins did not induce moDC death (**Supplementary Fig. S8a-c**) and produced modest alterations in CD86 and MHC class II expression (**Supplementary Fig. S8d**). These results suggest that soluble tumour-derived components can influence moDC phenotype but are insufficient to induce moDC death. We extended direct co-culture analyses to *C1GALT1*-knockout gastric cancer AGS and colorectal cancer SW480 cells, both of which expressed Tn and sTn (**Supplementary Fig. S9a**). These experiments revealed model-specific reductions in CD86 and HLA-DR, respectively, without comparable moDC death (**Supplementary Fig. S9b-d**). Immature *O*-glycosylation therefore perturbs moDC activation across tumour models, whereas moDC death is context-dependent. Assessment of tumour-cell viability further showed that both glycoengineered T24 models and *C1GALT1*-knockout SW480 cells underwent less death during moDC co-culture than their WT counterparts, whereas no comparable protection was observed in *C1GALT1*-knockout AGS cells (**Supplementary Fig. S10a-c**). These findings indicate that immature *O*-glycosylation can both perturb moDC function and favour tumour-cell survival during tumour-moDC interactions, although the latter effect varies across tumour models.

To assess potential impact for adaptive immunity, we exposed iDCs or mDCs to glycoengineered or WT T24 cells for 24 h before co-culture with autologous T cells (**Fig. 2m and Supplementary Fig. S11**). T cell viability was reduced following culture with tumour-exposed moDCs, most prominently after mDC exposure to Tn/sTn-expressing cells (**Fig. 2n**). Direct tumour-T cell co-culture did not reduce T cell viability (**Supplementary Fig. S12a**), indicating that the increased death observed in the priming assay was predominantly mediated through tumour pre-conditioned moDCs. Nevertheless, direct exposure to glycoengineered tumour cells increased regulatory T cell frequency (**Supplementary Fig. S12b**), revealing an additional moDC-independent effect on T cell state. In moDC-primed cultures, CD69 induction was reduced on CD8^+^ and CD4^+^ T cells, particularly following mDC priming (**Fig. 2o and Supplementary Fig. S13**). This was accompanied by increased CD4^+^CD25^high^CD127^low^ regulatory T cell frequencies (**Fig. 2p**). T cell-mediated killing of glycoengineered tumour cells was also reduced (**Supplementary Fig. S10d**). Collectively, these findings show that immature tumour *O*-glycans compromise macrophage and moDC function, thereby limiting productive T cell priming and cytotoxicity.

### Immature glycosylation on CD44s induces a pyroptosis-associated dendritic-cell death programme

We next asked whether moDC death required exposed immature *O*-glycans on the tumour-cell surface. Pre-incubation of glycoengineered T24 cells with unconjugated VVA partially rescued moDC viability (**Supplementary Fig. S14**), indicating that accessible Tn-bearing structures contribute to the response. Because conditioned medium did not reproduce moDC death, we reasoned that a membrane glycoprotein carrying immature *O*-glycans might provide the relevant ligand. CD44s was a strong candidate because it is a major carrier of truncated *O*-glycans in bladder cancer^14,15^. Notably, T24 cells expressed abundant CD44s, whereas AGS and SW480 cells, which did not consistently induce moDC death, showed no CD44s expression (**Supplementary Fig. S15a,b**). A synthetic CD44s-Tn glycopeptide was sufficient to induce moDC death, whereas the corresponding unglycosylated peptide and a Tn-glycosylated MUC16 control peptide were inactive (**Supplementary Fig. S15c**). CD44 knockdown reduced moDC death selectively in glycoengineered T24 co-cultures (**Fig. 3a**), whereas ectopic CD44s expression conferred death-inducing activity on glycoengineered AGS cells (**Fig. 3b and Supplementary Fig. S15d**). Inhibition of *N*-glycosylation synthesis with tunicamycin did not reduce moDC death (**Fig. 3c and Supplementary Fig. S15e**) and produced no consistent restoration of CD86 or HLA-DR (**Fig. 3d,e and Supplementary Fig. S16a,b**). By contrast, CD44 knockdown both improved moDC viability and partially restored maturation-associated markers (**Fig. 3c-e**). These results identify CD44s as a protein scaffold required for the dendritic-cell response to immature tumour *O*-glycans.

**Fig. 3.**
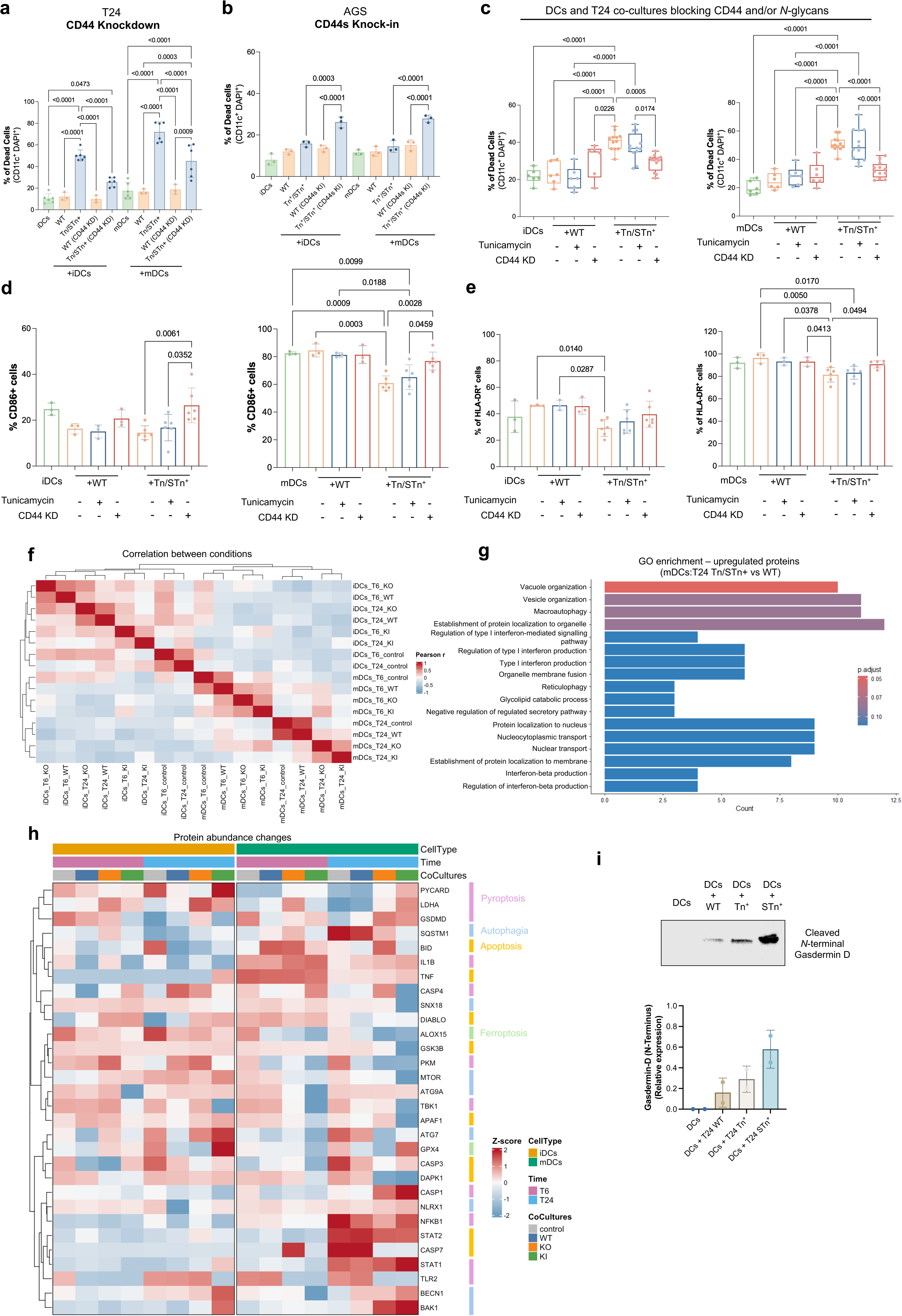
Tumour-cell CD44s mediates immature *O*-glycan-induced DC death and pyroptotic signalling. **a** CD44 is required for Tn/sTn-induced DC death. CD44 knockdown in Tn/sTn-expressing T24 cells reduced DAPI^+^CD11c^+^ death among iDCs and mDCs. **b** CD44s expression confers DC-killing capacity on Tn/sTn-expressing cells. CD44s knock-in increased iDC and mDC death induced by glycoengineered AGS gastric cancer cells. **c-e** CD44-dependent DC death and impaired activation do not require tumour-cell *N*-glycans. Tunicamycin-mediated inhibition of *N*-glycosylation did not rescue DC viability or activation, whereas CD44 knockdown reduced DC death (**c**) and restored CD86 (**d**) and HLA-DR expression (**e**), most clearly in mDCs. **f** DC proteomes segregate by maturation state and are further shaped by tumour glycosylation and exposure time. Pearson correlation matrix of iDCs and mDCs cultured alone or with WT, Tn- or sTn-expressing T24 cells for 6 or 24 h. **g** Tn/sTn-expressing cells induce vesicular, autophagic and type I interferon-related programmes in mDCs. GO analysis of proteins upregulated after 24 h exposure highlighted vesicle and vacuole organization, macroautophagy and type I interferon pathways; bars show counts and adjusted *P* values. **h** Tn/sTn exposure preferentially increases pyroptosis-associated proteins in mDCs, alongside changes in autophagy, apoptosis and ferroptosis regulators. Heatmap shows Z-scored protein abundances across DC states, timepoints and co-culture conditions. **i** Tn/sTn-expressing cells increase GSDMD cleavage in DCs, consistent with pyroptotic signalling. Representative immunoblot and densitometric quantification of *N*-terminal GSDMD. Kruskal-Wallis test followed by Dunn’s multiple-comparisons test. Exact *P* values are shown.

To characterize the associated cell-death programme, we performed temporal proteomic profiling of immature and mature moDCs after co-culture with WT or glycoengineered T24 cells for 6 or 24 h (**Supplementary Tables S2 and S3**). Glycoengineered cells induced broader proteomic remodeling than WT cells, particularly in mature moDCs at 24 h (**Fig. 3f**). Proteins increased in mature moDCs exposed to Tn/sTn-expressing cells were enriched in vesicle organization, macroautophagy and type I interferon-related pathways (**Fig. 3g**). The regulated protein set also contained multiple inflammasome- and pyroptosis-associated components, including PYCARD, CASP1, IL1B and GSDMD (**Fig. 3h**). Consistent with this signature, moDCs exposed to glycoengineered cells showed increased *N*-terminal gasdermin D (GSDMD) cleavage (**Fig. 3i**), together with increased IL-1β secretion (**Fig. 2l**). Thus, immaturely glycosylated CD44 induces a moDC death response accompanied by inflammasome activation and a pyroptosis-associated molecular programme.

### CD206 functionally engages immaturely glycosylated CD44

To identify moDC receptors that engage tumour-associated immature *O*-glycans, we combined two orthogonal discovery strategies (**Fig. 4a**). First, using the GlycoAvatar platform^20^, in which magnetic particles are coated with VVA-enriched glycoproteins from glycoengineered T24 cells, we identified 266 interacting proteins shared between immature and mature moDCs (**Fig. 4b-e, Supplementary Tables S4 and S5**). This set was enriched in membrane and receptor-associated functions, including receptor-mediated endocytosis, phagocytosis and vesicle organization (**Fig. 4d**). Second, affinity purification with the death-inducing CD44s-Tn glycopeptide (**Supplementary Table S6**) recovered 19 plasma-membrane proteins (**Fig. 4f, Supplementary Table S7**). Integration of both datasets identified the mannose receptor CD206/MRC1 as the highest-confidence shared candidate (**Fig. 4e,f**). In a reciprocal assay, biotinylated CD206 recovered CD44 from glycoengineered T24 membrane extracts, including after PNGase F treatment, independent of *N*-glycan status (**Fig. 4g,h, Supplementary Table S8**). Soluble CD206 (MRC1) bound Tn- and Tn/sTn-expressing T24 cells but not to WT cells, and binding was reduced by CD44 knockdown, suggesting interaction between both receptors (**Fig. 4i**). CD206 internalization was detected in moDCs co-cultured with glycoengineered, but not WT tumour cells, suggesting that this interaction is dependent on tumour-associated glycosylation (**Fig. 4j**).

**Fig. 4.**
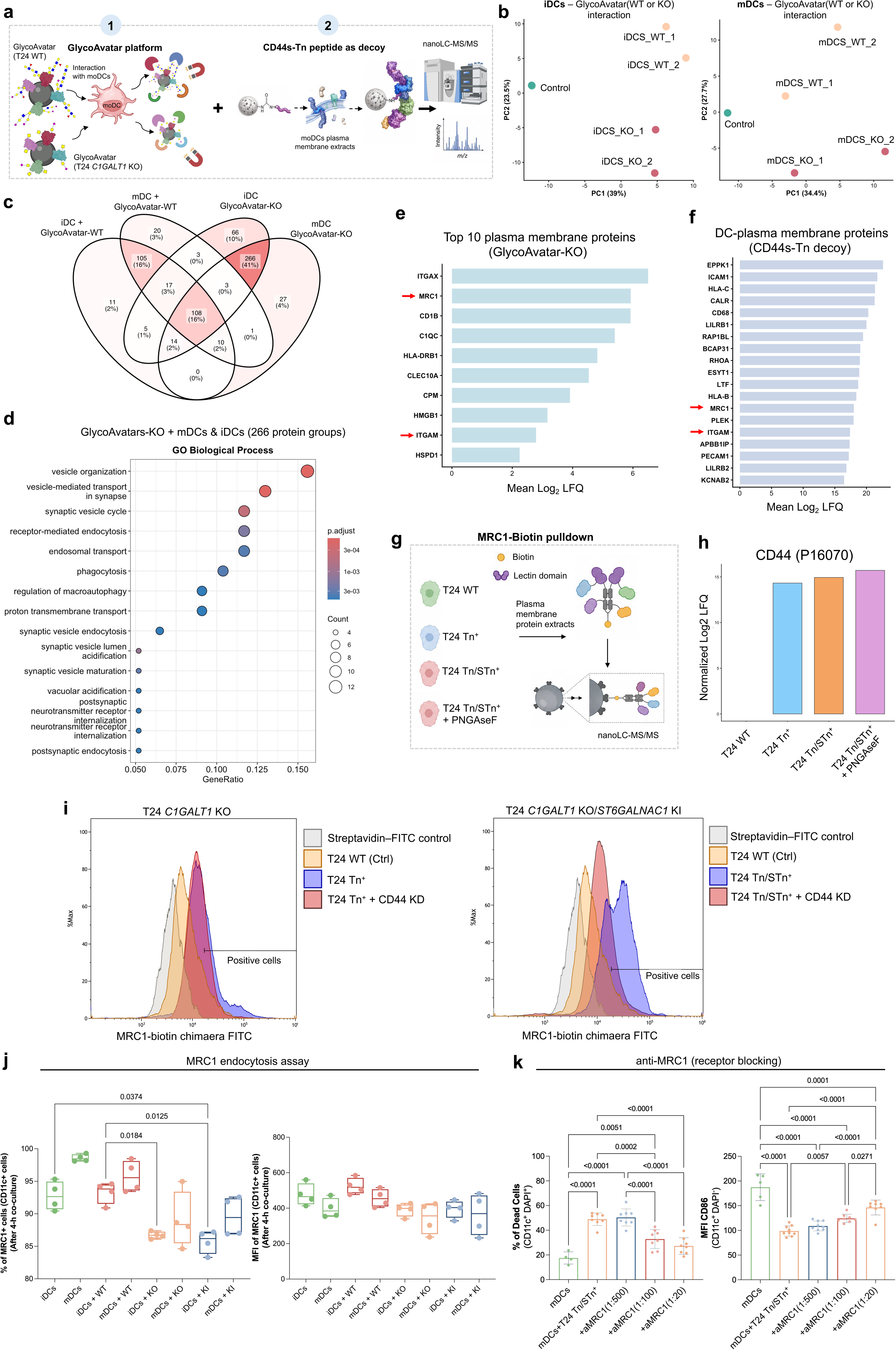
CD206 binds CD44 carrying immature *O*-glycans and mediates DC loss and impaired activation. **a** Experimental design of the GlycoAvatar and CD44s-Tn peptide-pulldown strategies for identifying DC receptors for tumour-derived immature *O*-glycans. **b** Immature *O*-glycans generate distinct DC-interaction profiles. Principal-component analysis of proteins recovered from iDCs and mDCs interacting with WT or *C1GALT1*-knockout GlycoAvatars. **c** GlycoAvatar screening identified 266 proteins shared by iDCs and mDCs interacting with *C1GALT1*-knockout glycoproteins. Venn diagram shows proteins recovered across DC states and GlycoAvatar backgrounds. **d** Shared interactors were enriched in vesicle organization, receptor-mediated endocytosis, phagocytosis and macroautophagy. Dot plot shows GO biological processes, gene ratios, protein counts and adjusted *P* values. **e,f** CD206 mannose receptor/MRC1 converges across both receptor-discovery approaches. Plasma-membrane candidates recovered with *C1GALT1*-knockout GlycoAvatars (**e**) and the CD44s-Tn peptide (**f**), ranked by mean log_2_ LFQ abundance. **g** Design of the reciprocal biotinylated-CD206 pulldown using membrane extracts from glycoengineered T24 cells. **h** CD206 recovers CD44 from Tn/sTn-expressing cells independently of *N*-glycans. Normalized CD44 abundance following CD206 pulldown, including PNGase F-treated extracts. **i** CD206/MRC1 binds preferentially to Tn/sTn-expressing cells through CD44. Representative flow-cytometry histograms of MRC1-biotin binding with or without CD44 knockdown. **j** Tn/sTn-expressing cells induce CD206 internalization in iDCs and mDCs, after 4 h of co-culture. Surface CD206 positivity and fluorescence intensity after 4 h co-culture. **k** CD206 blockade rescues DC viability and partially restores activation. Dose-dependent effects of anti-CD206 on mDC death and CD86 expression after co-culture with Tn/sTn-expressing cells. Ordinary one-way ANOVA followed by Dunnett’s multiple-comparisons test. Exact *P* values are shown.

Because MGL and Siglec-15 are frequently described receptors for Tn^17^ and sTn^21^, respectively, we asked whether they might provide alternative routes of glycan recognition. CD206 was expressed by both immature and mature moDCs, whereas MGL was less abundant in mature moDCs and Siglec-15 was not detected in this system (**Supplementary Fig. S17**). We therefore tested whether CD206 mediated the response to Tn/sTn-expressing T24 cells. Antibody-mediated CD206 blockade dose-dependently rescued mDC viability and partially restored CD86 expression in surviving cells (**Fig. 4k**). Together, receptor discovery, reciprocal affinity purification, cell-surface binding, internalization and functional blockade identify CD206 as a receptor for CD44 carrying immature *O*-glycans that contributes to moDC death and impaired activation. Persistence of CD206-CD44 recovery after PNGase F treatment further indicates that this interaction can occur independently of tumour-cell *N*-glycans.

### CD44s-associated immature-glycosylation state marks immune exclusion in BLCA

To assess the clinical relevance of tumour CD44 in immature glycosylation, we stratified TCGA-BLCA tumours by combined *CD44s*/*CD44st* abundance within cohorts with low *C1GALT1* or *C1GALT1C1* expression (**Fig. 5a**). Transcript-level analyses included *CD44st* because both isoforms share the same ectodomain and thus the same potential *O*-glycosites, but CD44st has a truncated cytoplasmic tail, failing downstream signaling activation^14^. Because reduced *C1GALT1* or *C1GALT1C1* expression is expected to favour accumulation of Tn, we next asked whether the association between CD44 abundance and outcome was also evident in tumours with increased sTn-associated glycosylation, using *ST6GALNAC1* expression as a complementary stratification. High CD44s/CD44st abundance was associated with shorter survival, including in the *ST6GALNAC1*-high subset (**Fig. 5b**). Within this sTn-prone subset, *CD44s*/*CD44st*-high tumours showed reduced type 1/DC-associated and increased immunoregulatory/TAM-associated transcription (**Fig. 5c**). *MRC1* (CD206) and *SIGLEC15* were higher, whereas *CLEC10A* (MGL) was unchanged (**Fig. 5d**). This is consistent with the CD206-dependent DC dysfunction observed experimentally.

**Fig. 5.**
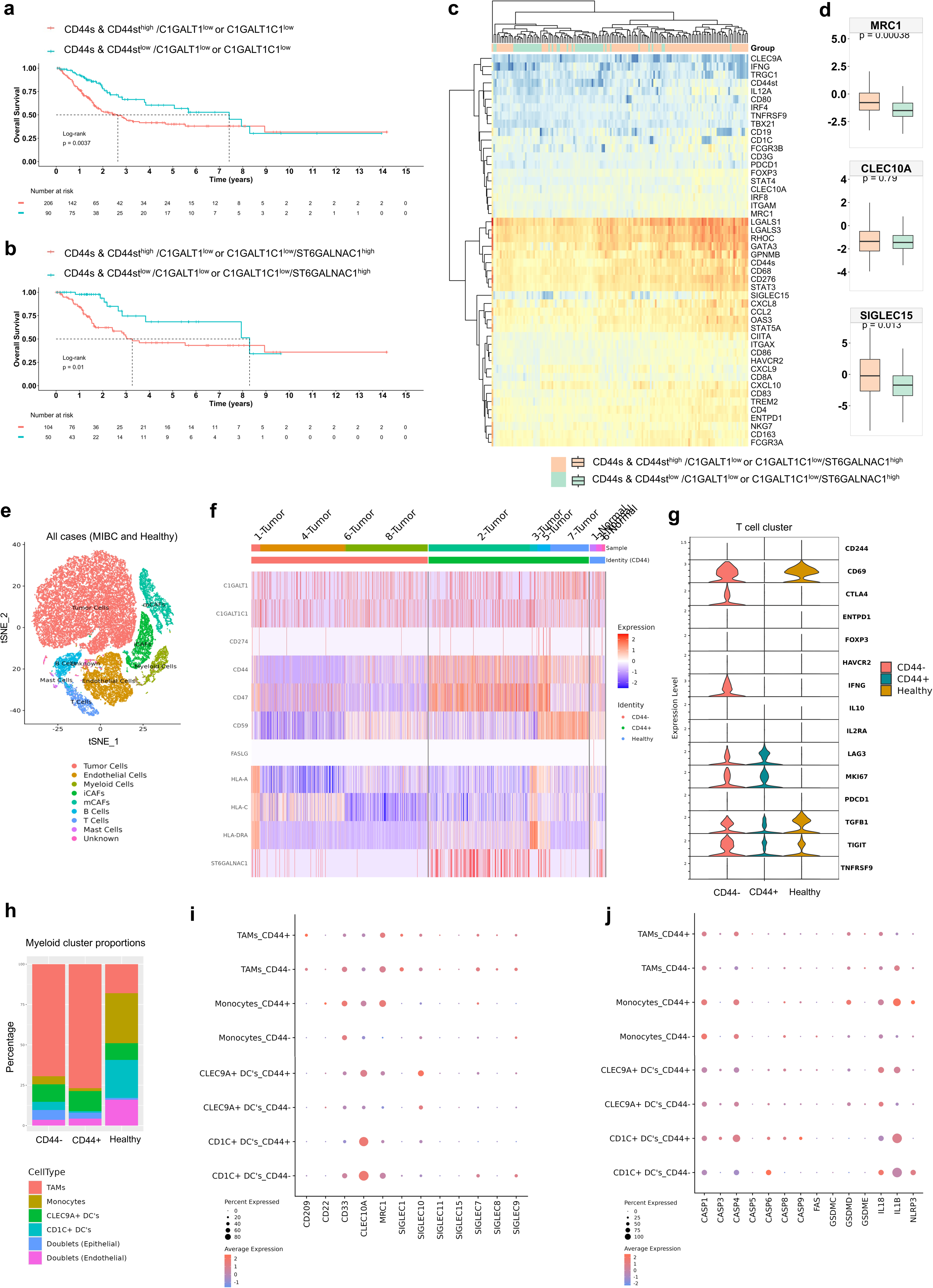
High abundance of *CD44s* and *CD44st* transcripts, which encode isoforms sharing a common ectodomain, associates with poor-prognosis and immunosuppression in BLCA within a transcriptional profile consistent with immature *O*-glycosylation. **a,b** High combined *CD44s* and *CD44st* transcript abundance is associated with shorter overall survival in TCGA-BLCA tumours with impaired *O*-glycan elongation (low *C1GALT1* or *C1GALT1C1* expression), either alone or together with enhanced Tn sialylation (high *ST6GALNAC1*). The encoded isoforms share the same extracellular glycosylation scaffold, but CD44st contains a truncated cytoplasmic tail. Kaplan-Meier curves are compared using two-sided log-rank tests. **c** Tumours with high *CD44s/CD44st* transcript abundance and a profile favoring sTn formation show reduced type 1 and DC-associated gene expression but increased immunoregulatory and macrophage-associated gene expression. d *CD44s*/*CD44st*-high, sTn-prone tumours show higher *MRC1* (CD206) and *SIGLEC15* (sTn receptor), whereas *CLEC10A* (MGL; Tn receptor) is unchanged. **e** t-SNE map of tumour, immune and stromal cells from MIBC and adjacent healthy bladder mucosa. **f** Both *CD44s* groups show low *C1GALT1/C1GALT1C1* and comparable immunogenicity, but *CD44s*-positive tumours have higher *ST6GALNAC1* and anti-phagocytic *CD47*. **g** T cells in CD44s-positive tumours showed reduced activation (*CD69*) and reduced type 1 effector function (*IFNG*) relative to CD44s-negative tumours. **h** MIBC groups are TAM-dominated relative to healthy bladder; *CD44s*-positive tumours have the highest TAM and lowest monocyte and CD1C⁺ DC representation. **i** TAMs in both groups expressed *MRC1*, with broader expression in monocytes and CLEC9A⁺ DCs from *CD44s*-positive tumours. **j** CD44s-positive tumours showed DC inflammasome/pyroptosis transcripts (*CASP1, CASP4, IL18, IL1B*); monocytes showed the strongest GSDMD-IL1B-NLRP3 signature.

We next reanalysed single-cell transcriptomes^22^ from MIBC and healthy bladder mucosa (**Fig. 5e and Supplementary Fig. S18a**). As immune cells also express *CD44*, tumours were classified by *CD44* expression within the tumour-cell cluster. Both *CD44*-negative and *CD44*-positive tumours showed low tumour-cell *C1GALT1*/*C1GALT1C1* expression (**Supplementary Fig. S18b**), consistent with immature glycosylation. *CD44*-positive tumours, however, expressed more *ST6GALNAC1* and *CD47*, consistent with enhanced Tn sialylation and anti-phagocytic protection, respectively, despite otherwise comparable immunogenicity profiles (**Fig. 5f**). Despite preserved proliferative capacity, as indicated by comparable *MKI67* expression, T cells showed increased *LAG3* expression and reduced *IFNG*, consistent with impaired effector function (**Fig. 5g**). Both groups were tumour-associated macrophages (TAM)-dominated relative to healthy mucosa; however,

*CD44* positive tumours displayed the highest TAM and lowest monocyte and CD1c+ DC representation (**Fig. 5h and Supplementary Fig. S19**). Consistent with the bulk *CD44s*/*CD44st* analysis, *MRC1* was enriched in TAMs from both groups and more broadly expressed in monocytes and CLEC9A+ DCs from *CD44*-positive tumours. Other glycan receptors remained lineage-restricted and largely *CD44*-independent (**Fig. 5i and Supplementary Fig. S20a**). *CD44*-positive tumours also showed stronger inflammasome/pyroptosis transcription in monocytes and DCs, with higher *CASP1*, *CASP4*, *IL18* and *IL1B* in DCs and a stronger *GSDMD*-*IL1B*-*NLRP3* signature in monocytes (**Fig. 5j and Supplementary Fig. S20b**). Together, these analyses define a *CD44*-associated, immature-glycosylation state linked to poor survival, TAM dominance, reduced DC representation and impaired T-cell activation.

### Immature tumour *O*-glycosylation promotes immune exclusion *in vivo*

To test whether immature *O*-glycosylation alters antitumour immunity *in vivo*, we generated a syngeneic model using glycoengineered MB49 bladder cancer cells (**Supplementary Fig. S21**). Because stable *C1GALT1* disruption could not be achieved in MB49 cells, we targeted *C1GALT1C1*/Cosmc, the essential chaperone required for C1GalT1 activity. *C1GALT1C1* disruption induced homogeneous Tn and sTn expression together with loss of extended *O*-glycans (**Supplementary Fig. S21a, b**). MB49 cell line also expresses CD44 (**Supplementary Fig. S21c**), as observed in the Human bladder cancer cell line T24. Consistent with the findings in human moDCs, *in vitro* co-culture with MB49 *C1GALT1C1* KO cells also increased death of murine bone marrow-derived dendritic cells (BMDCs) compared with WT co-cultures (**Supplementary Fig. S22**). This was accompanied by reduced MHC-II (I-A/I-E) expression, as previously reported for human DCs.

Following subcutaneous implantation in immunocompetent C57BL/6J mice (**Fig. 6a**), glycoengineered MB49 cells displayed accelerated tumour growth under both inoculum conditions (**Fig. 6b,c**) and produced larger endpoint tumour masses (**Fig. 6d**). Mice bearing glycoengineered tumours also presented enlarged spleens (**Fig. 6e**), consistent with systemic consequences of rapid tumour progression. This accelerated growth provides *in vivo* support for our previous association of Tn/sTn with aggressive and invasive bladder tumours^11,14^.

**Fig. 6.**
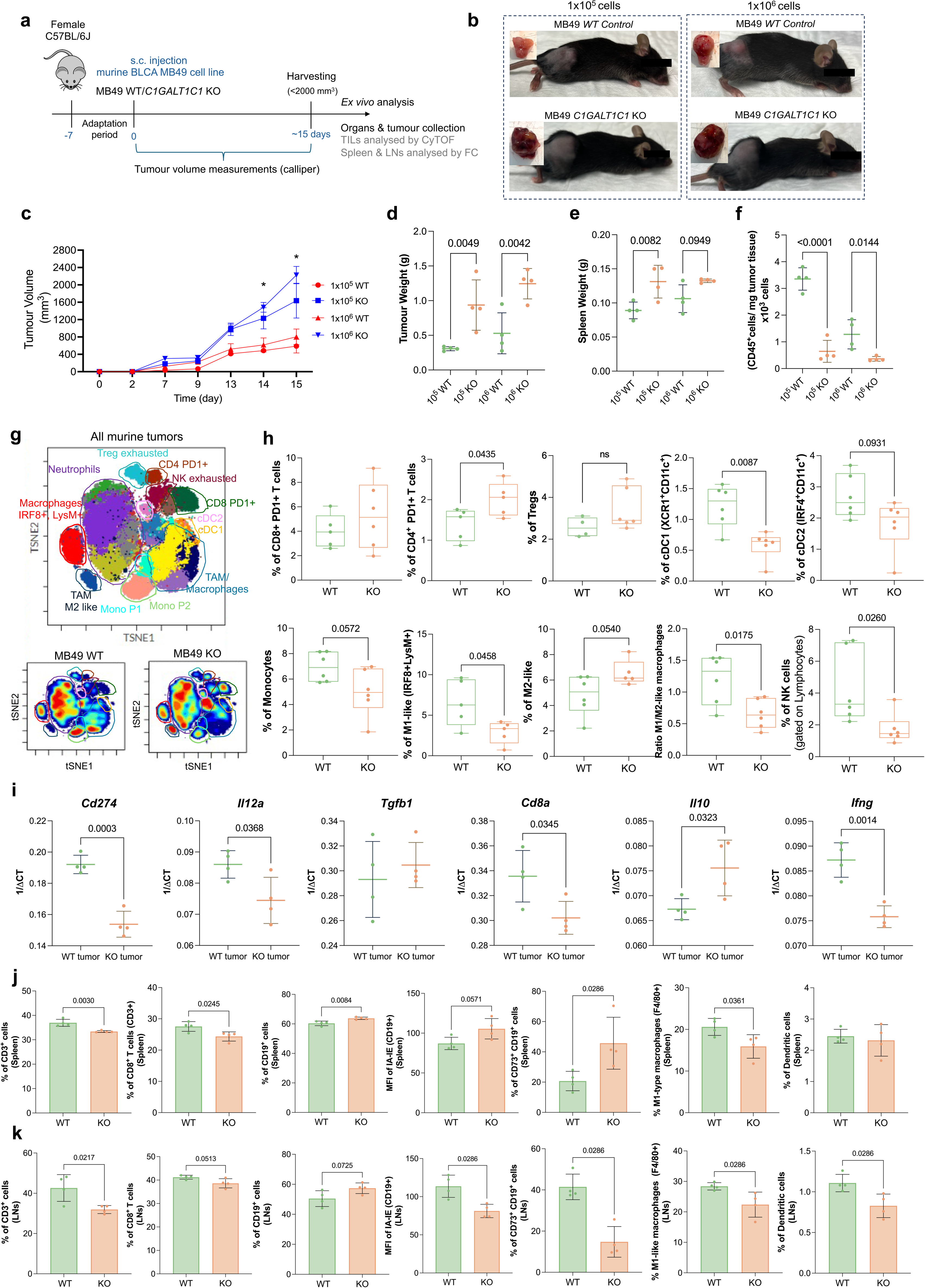
*C1GALT1C1* knockout in MB49 bladder cancer cells accelerates tumour growth and suppresses myeloid and cytotoxic lymphoid immunity in syngeneic mice. **a** Experimental design. C57BL/6J mice were inoculated subcutaneously with 10^5^ or 10^6^ WT or *C1GALT1C1*-knockout MB49 cells; tissues were collected 15 days later for immune phenotyping. Tumour growth kinetics were measured over time. b-d *C1GALT1C1*-knockout tumours grew faster and reached greater endpoint weights. Representative images (**b**), growth curves (**c**) and tumour weights (**d**). **e, f** Mice bearing *C1GALT1C1*-knockout tumours showed increased spleen weight (**e**) and their tumours contained fewer CD45⁺ leukocytes per milligram (**f**). **g, h** *C1GALT1* knockout tumours are depleted of antigen-presenting, inflammatory myeloid and cytotoxic lymphoid populations. t-SNE maps (**g**) and CyTOF quantification showing increased CD4^+^PD-1^+^ T cells and reduced cDC1, IRF8^+^LysM^+^ M1-like macrophages, M1/M2-like macrophage ratio and NK cells (**h**). **i** *C1GALT1C1*-knockout tumours showed reduced *Cd274*, *Il12a*, C*d8a* and *Ifng*, and increased *Il10* expression. **j** Spleens showed fewer CD3^+^ and CD8^+^ T cells and M1-like macrophages, but more CD19^+^ and activated CD73^+^ B cells. **k** Tumour-draining lymph nodes showed fewer CD3^+^ T cells, CD73^+^ B cells, M1-like macrophages and DCs and reduced I-A/I-E expression on B cells. n=4 mice per group (**b-f, i-k**) and n=6 mice per group (**g, h**). Data are mean ± s.d. (**c-f, i-k**); Two-way ANOVA with Šídák’s multiple-comparisons test (**c**); two-sided unpaired t-tests or Mann-Whitney U-tests according to distribution (**d-k**). Exact P values are shown; \**P* < 0.05.

Interestingly, glycoengineered tumours contained fewer CD45⁺ immune cells per milligram of tumour under both inoculum conditions (**Fig. 6f**). High-dimensional analysis of the residual infiltrate showed depletion of cDC1, IRF8⁺LysM⁺ M1-like macrophages and NK cells, together with a reduced M1/M2-like macrophage ratio (**Fig. 6g,h and Supplementary Figs. S23 and S24**). CD4⁺PD-1⁺ T cells were increased, whereas cDC2 (CD1+), regulatory T cells and CD8⁺PD-1⁺ T cells showed non-significant trends. Whole-tumour transcriptional analysis showed reduced *Ifng*, *Il12a*, *Cd8a* and *Cd274* and increased *Il10*, whereas *Tgfb1* remained unchanged (**Fig. 6i**). Secondary lymphoid organs (**Supplementary Fig. S25**) were also affected: spleens from mice bearing glycoengineered tumours contained fewer CD3⁺ and CD8⁺ T cells, more CD19⁺ cells proportion and fewer M1-like macrophages (**Fig. 6j**), while tumour-draining lymph nodes showed also reduced DC abundance (**Fig. 6k**). Together, these findings indicate that immature tumour *O*-glycosylation compromises both myeloid and cytotoxic lymphoid components of anti-tumour immunity, locally and in secondary lymphoid organs. Overall, this model recapitulates the DC depletion, myeloid skewing and impaired type 1 and cytotoxic immunity observed in human BLCA and establishes immature tumour *O*-glycosylation as a driver of immunosuppression.

### CD206 blockade restrains tumours with immature *O*-glycosylation *in vivo*

Having shown that CD206 blockade rescued moDC viability *in vitro*, we next asked whether targeting this receptor could restrain the growth of tumours with immature *O*-glycosylation *in vivo*. We first confirmed that *C1GALT1C1*-knockout MB49 tumours recapitulated the surface co-distribution of CD44 and VVA-reactive glycans observed in human tumours (**Fig. 7a**). We then treated mice bearing these tumours with murine anti-CD206/MRC1 in 3 interspersed administrations (**Fig. 7b**). Treatment slowed tumour growth, with the highest dose producing the largest reductions in endpoint tumour volume (**Fig. 7c,d**) and tumour weight (**Fig. 7e**). Treatment increased the number of CD45⁺ cells relative to tumour weight and remodelled the intratumoural myeloid compartment, reducing CD206⁺ leukocytes and TAMs and increasing both the M1/M2-like macrophage ratio and cDC2 representation (**Fig. 7e**). The higher antibody dose also increased effector-memory CD4⁺ and CD8⁺ T cells and B cells, whereas total T-cell abundance changed only modestly. This shift towards a more effector-rich infiltrate was accompanied by reduced *Tgfb1* and increased *Cd8a* transcripts in whole tumours (**Fig. 7f**). In tumour-draining lymph nodes, CD206 blockade increased M1-like macrophages, XCR1⁺ cDC1 and CD8⁺ T cells, increased the CD8/CD4 T-cell ratio and reduced the frequency of naïve CD4⁺ and CD8⁺ T cells (**Fig. 7g**). Activated B cells were also increased. Together, these data show that CD206 blockade restrains the growth of tumours with immature glycosylation while restoring myeloid and lymphoid antitumour immunity within tumours and tumour-draining lymph nodes.

**Fig. 7.**
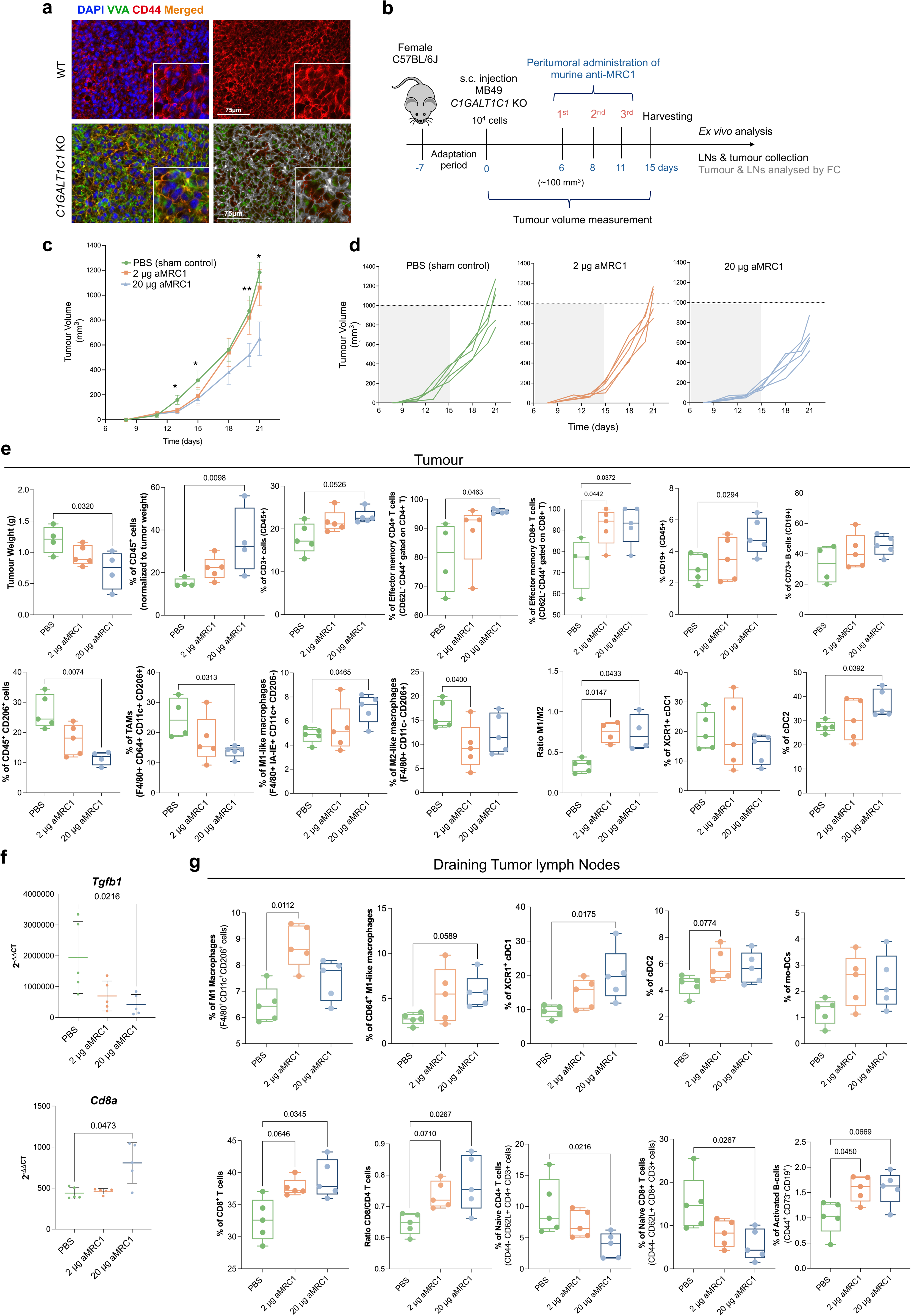
CD206 blockade restrains *C1GALT1C1*-knockout bladder tumour growth and restores antitumour immunity locally and in tumour-draining lymph nodes. **a** *C1GALT1C1*-knockout MB49 cells showed surface Tn expression, detected by VVA reactivity, that spatially overlapped with CD44. Representative immunofluorescence; scale bars, 75 μm. b Experimental design for assessing the effects of CD206 blockade on tumour growth and antitumour immunity. C57BL/6J mice bearing C1GALT1C1-knockout MB49 tumours received three peritumoural doses of PBS or anti-CD206 (2 or 20 μg), followed by tumour-growth monitoring and immune profiling. c,d Anti-CD206 restrained tumour growth, most strongly at 20 μg. e CD206 blockade reduced endpoint tumour weight and increased CD45+ cells relative to tumour weight. Effector-memory CD4+ and CD8+ T cells, B cells, M1-like macrophages, the M1/M2-like macrophage ratio, and cDC2 were increased, whereas CD206+ leukocytes and TAMs were decreased. **f** CD206 blockade reduced tumour *Tgfb1* and increased *Cd8a* expression, supporting reduced immunoregulation and enhanced CD8+ T-cell immunity. **g** In tumour-draining lymph nodes, CD206 blockade increased M1-like macrophages, XCR1^+^ cDC1, CD8^+^ T cells, the CD8/CD4 ratio and activated B cells, while reducing naive CD4^+^ and CD8^+^ T cells. Two-way ANOVA with Šídák’s multiple-comparisons test (**c**); Kruskal-Wallis test followed by Dunn’s multiple-comparisons test (**e-g**). Exact *P* values are shown; \**P* < 0.05, \*\**P*<0.01

## Discussion

Immature protein *O*-glycosylation is a recurrent feature of aggressive malignancies, yet its contribution to immune escape remains incompletely understood^23,24^. Here, we identify immature glycans in BLCA as active determinants of immunosuppression and appointed as targets for therapeutic modulation. In tumours, immature glycosylation was associated with reduced antigen-presentation and cytotoxic immune activity and with enrichment of immunoregulatory myeloid features. Functionally, it promoted DC loss, the accumulation and polarization of suppressive macrophages, defective T-cell priming and immune exclusion. Because Tn and sTn arise from the same biosynthetic pathway and coexist in tumour models and human lesions, their individual contributions cannot yet be fully resolved^25^. Nevertheless, the stronger immunosuppressive effects observed in sTn-enriched settings suggest that sialylation can further exacerbate this dysfunction. Together, these findings support a model in which the immature tumour glycocalyx coordinates myeloid dysfunction with defective lymphoid priming and function, leading to the establishment of an immunosuppressive state.

The central advance of this study is the identification of a glycosylation-dependent interaction between tumour-cell CD44 and the immune cell mannose receptor CD206 that connects glycocalyx remodelling to APC loss. Immature *O*-glycans have long been implicated in myeloid regulation through C-type lectin receptors, such as MGL, and members of the Siglec family, which detect aberrant glycoforms and imprint immunosuppressive programmes^18,26,27^. Consistent with this framework, tumour profiling revealed enrichment of selected lectin-receptor and immunoregulatory features in lesions displaying immature glycosylation, including CD206. In glycoengineered bladder carcinoma models, receptor discovery, affinity purification, cell-surface binding, internalization and functional blockade converged on CD206 as a mediator of DC dysfunction by pyroptosis. To our knowledge, recognition of CD44 by CD206 has not previously been reported. Our data support, through distinct molecular approaches, a functional receptor-ligand relationship through which immature glycosylation of tumour-cell CD44 contributes to DC dysfunction.

CD206 is classically described as a scavenger-type lectin that recognizes mannose-containing *N*-glycans, but its ligand repertoire also includes fucose-, GlcNAc- and sulphated glycan structures^28^. Its modular architecture, comprising an *N*-terminal cysteine-rich region, a fibronectin type II domain and eight C-type lectin-like domains, enables recognition of diverse glycoconjugates and extracellular matrix components, including collagen^29^. Notably, CD206 has previously been shown to recognize chondroitin sulphate-modified CD44 during lymphocyte trafficking^30^. Our findings extend this interaction to immaturely *O*-glycosylated tumour CD44, although whether CD206 directly recognizes truncated *O*-glycans or glycosylation-dependent changes in CD44 presentation remains unclear. Interestingly, *O*-glycan truncation can enhance CD44-mediated hyaluronan binding^31^, raising the possibility that extracellular matrix interactions further influence CD44 organization and accessibility to CD206.

In our models, DC loss persisted after enzymatic removal of tumour-cell *N*-glycans, indicating that classical *N*-glycan-dependent mannose recognition is dispensable for this effect. We therefore propose that immature *O*-glycans on tumour-cell CD44s alter its surface presentation or molecular organization in a manner that enables CD206 engagement and initiates a dysfunctional myeloid response culminating in moDC loss. Whether CD206 directly recognizes truncated *O*-glycans or instead senses glycosylation-dependent changes in CD44s conformation or surface organization will require structural and mutational validation. More broadly, our findings indicate that immunosuppression is determined by both glycan state and protein context. Together, these features define which lectin receptors can be productively engaged, consistent with our previous observations using GlycoAvatars^20^. These findings provide a rationale for therapeutically disrupting the glycoprotein-lectin interaction that sustains DC suppression.

A further major finding is that the CD44s-CD206 interaction is functionally linked to pyroptotic moDC loss. Pyroptosis is commonly regarded as an inflammatory and potentially immunogenic form of programmed cell death, yet in our models it restrained antitumour immunity^32,33^. Inflammasome-associated moDC death depleted a critical antigen-presenting population, shifted surviving moDCs towards a dysfunctional phenotype and disrupted the cellular environment required for efficient T-cell priming. The induction of IL-1β, caspase-1, and gasdermin D was consistent with activation of inflammatory death pathways that undermined immune competence rather than promoting productive immunity. These findings extend previous evidence that tumour-associated glycans reprogramme DCs towards impaired co-stimulatory capacity, altered cytokine output and defective T-cell activation^34^.

In human bladder tumours, transcriptional programmes consistent with immature CD44s glycosylation were associated with poor survival, broad repression of antigen-presentation and immune-effector pathways, and enrichment of suppressive myeloid features. This was also associated with inflammasome- and pyroptosis-related features, reinforcing the connection between aberrant tumour glycosylation, immunosuppressive inflammation and loss of effective antigen-presenting activity.

In summary, our study identifies immature protein glycosylation as a driver of immune exclusion whose effects depend on both glycan composition and protein context. These findings reveal a targetable vulnerability in aggressive, glycosylation-driven cancers that remain poorly served by current precision approaches. Disrupting the CD44s-CD206 interaction provides a mechanistically grounded strategy for targeting tumours that are simultaneously invasive and immunosuppressive. Beyond its therapeutic implications, this work provides a framework for interpreting glycoimmune interactions and developing strategies to restore lymphoid priming and reduce immune escape mechanisms. Future studies should define the structural and functional basis of CD44s recognition by CD206 and determine whether targeting this interaction can cooperate with immunotherapy across diverse tumour types.

## Online Methods

This study combined transcriptomic, histological, cellular, proteomic and *in vivo* approaches to determine how immature tumour *O*-glycosylation shapes antitumour immunity. Bulk and single-cell BLCA transcriptomic datasets were analysed to relate glycosylation-associated states to immune-cell representation, antigen-presentation pathways and clinical outcome. Human bladder cancer specimens were examined by immunohistochemistry, multiplex immunofluorescence and cytokine-gene profiling. Glycoengineered tumour-cell models were used in co-cultures with human monocyte-derived macrophages, dendritic cells and T cells to assess myeloid polarization, dendritic-cell viability and maturation, cytokine production and downstream T-cell responses. Mechanistic experiments included lectin blocking, CD44 perturbation, glycopeptide assays, temporal proteomics, GlycoAvatar-based receptor discovery, reciprocal affinity purification, soluble-CD206 binding and CD206 blockade. Syngeneic MB49 mouse models were used to assess tumour growth, immune infiltration, secondary-lymphoid-organ remodeling and response to anti-CD206 treatment. Human and animal studies were conducted under the appropriate institutional ethics approvals. The complete procedures, antibodies, reagents, bioinformatic workflows and statistical analyses are provided in the accompanying extended Methods document.

## Author Contributions

AM, YvK and JAF designed research; AM, RF, DF, EF, AP and CL performed research; AM, AB and MdK performed bioinformatic analysis; MG, CP, FC, LLS, MJO, HL, YvK, and JAF contributed new reagents/analytic tools; AM, DF, MRS and LPA analyzed data; AM and JAF wrote the paper. All authors revised the paper.

