## Supplemental Figures S1-S25 for "CD44-CD206 axis drives dendritic cell loss and immunosuppression in immaturely glycosylated bladder tumours"

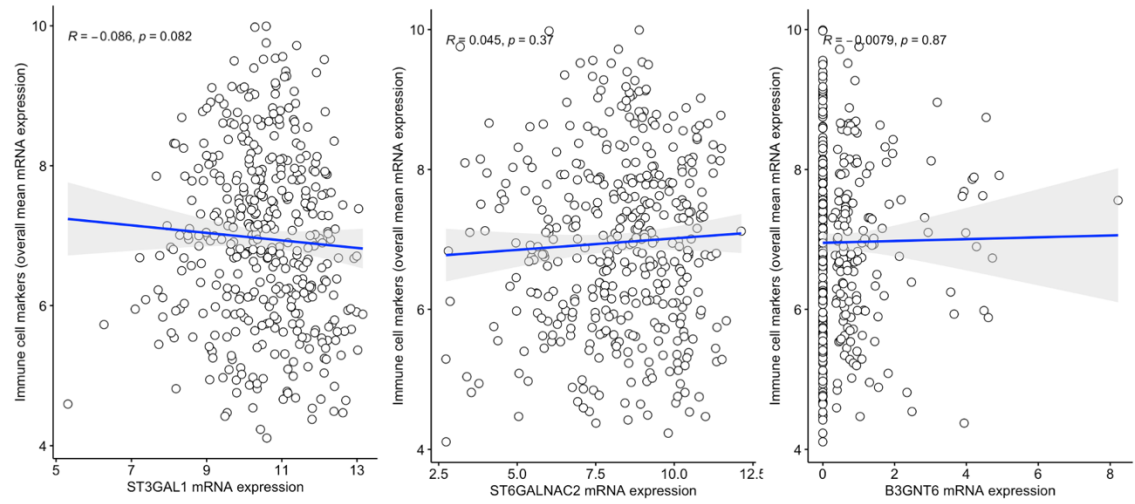

**Supplementary Fig. S1.** *ST3GAL1*, *ST6GALNAC2*, and *B3GNT6* expression levels are not correlated with immune cell markers in MIBC (n=407) from TCGA-BLCA.

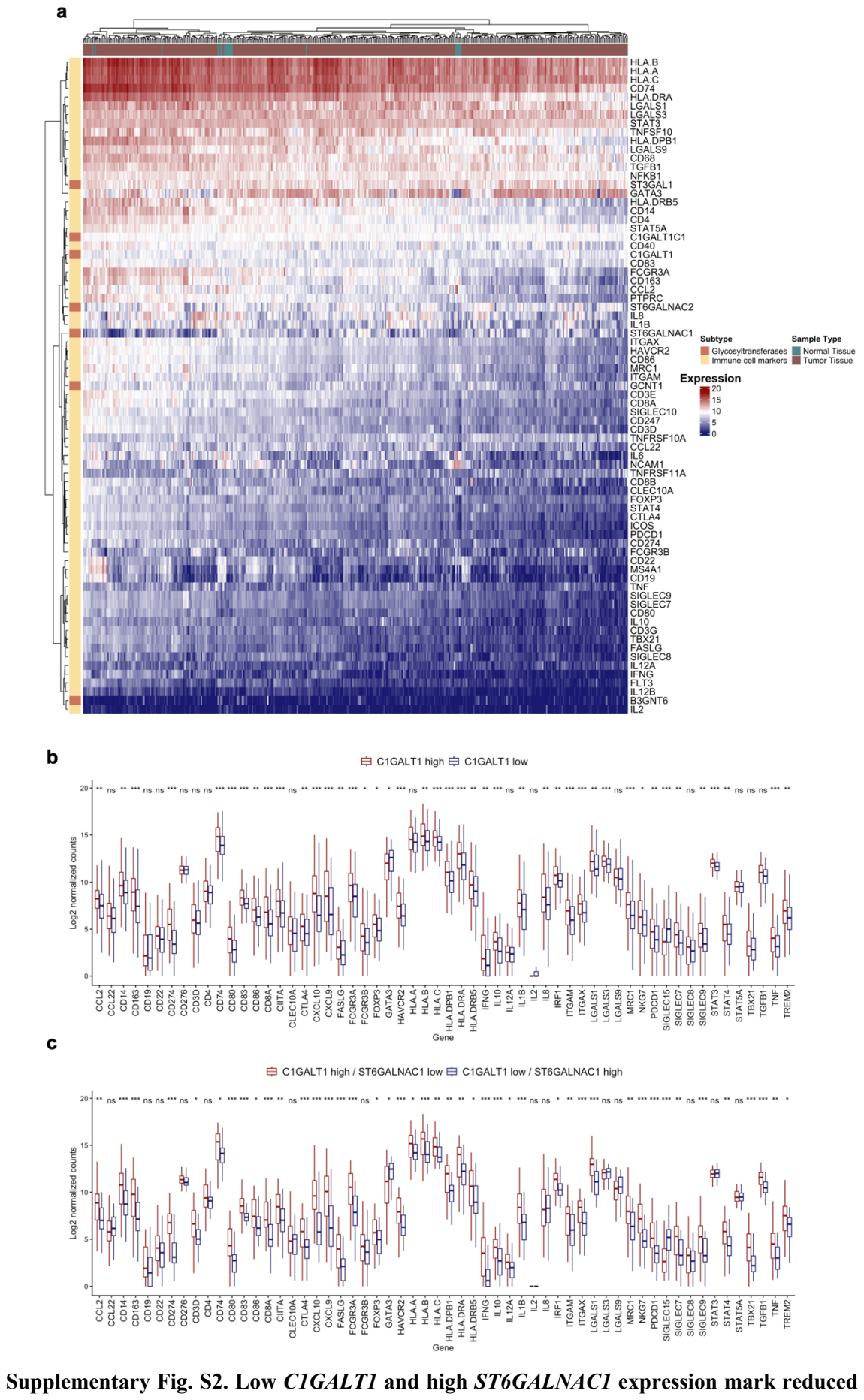

**immune-marker expression in TCGA-BLCA.** **a** Hierarchically clustered heat map showing the expression of *O*-glycosylation enzymes and immune-cell markers across normal bladder and primary TCGA-BLCA samples. Colours indicate $\log_2(\text{TPM} + 0.001)$ expression. **b** *CIGALT1*-low tumours, consistent with impaired *O*-glycan elongation and Tn accumulation, showed lower expression of *PTPRC* and a broad range of immune-cell markers than *CIGALT1*-high tumours. **c** *CIGALT1*-low/*ST6GALNAC1*-high tumours, with a transcriptional profile favouring sTn accumulation, similarly showed lower immune-marker expression than *CIGALT1*-high/*ST6GALNAC1*-low tumours. Groups represent the upper and lower expression tertiles among 407 primary tumours. Box plots show medians, interquartile ranges and ranges; two-sided Wilcoxon rank-sum tests. ns, not significant; \*$P < 0.05$, \*\*$P < 0.01$, \*\*\*$P < 0.001$ and \*\*\*\*$P < 0.0001$. The broad reduction in immune markers, including *PTPRC*, supports lower leukocyte representation in tumours with a transcriptional profile favouring short *O*-glycans.

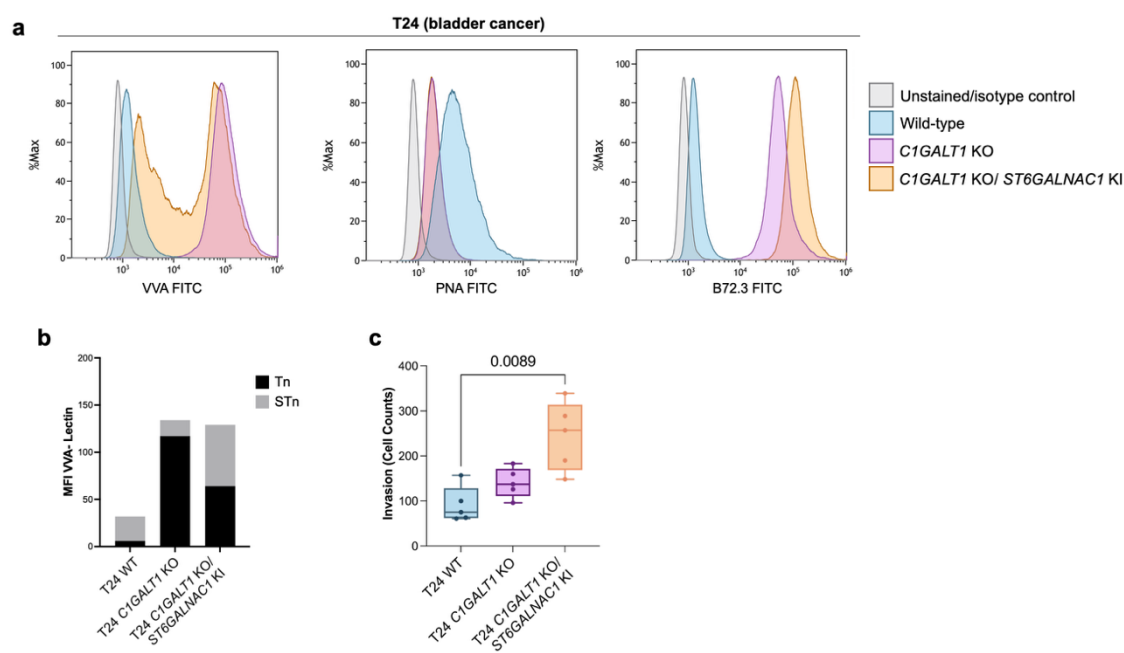

**Supplementary Fig. S3. *ST6GALNAC1*-mediated Tn sialylation enhances T24-cell invasion and reduces proliferation.** **a** Flow-cytometry analysis showed PNA-reactive core 1 *O*-glycans in wild-type T24 cells and increased VVA-reactive Tn following *C1GALT1* KO. *ST6GALNAC1* KI in *C1GALT1* KO cells increased B72.3-reactive sTn while retaining VVA reactivity. Unstained or isotype controls are shown. **b** Quantification confirmed Tn accumulation after *C1GALT1* loss and its partial conversion to sTn following *ST6GALNAC1* KI. To estimate the Tn: sTn ratio, Tn and sTn levels were assessed using VVA staining before and after neuraminidase (NeuAse) treatment. sTn-associated MFI was estimated as the difference between VVA MFI after NeuAse treatment and untreated: sTn MFI = VVA MFI (NeuAse-treated) – VVA MFI (untreated). **c** *C1GALT1* KO/*ST6GALNAC1* KI cells showed greater invasion than WT cells, whereas *C1GALT1* KO cells displayed an intermediate phenotype. Invasion was quantified by counting cells crossing the membrane. Each point represents an independent experiment. These findings indicate that Tn sialylation further promotes the invasive phenotype associated with truncated *O*-glycosylation. Statistical analyses were performed using the Kruskal–Wallis test followed by Dunn's multiple-comparisons test. Exact *P*-values are shown.

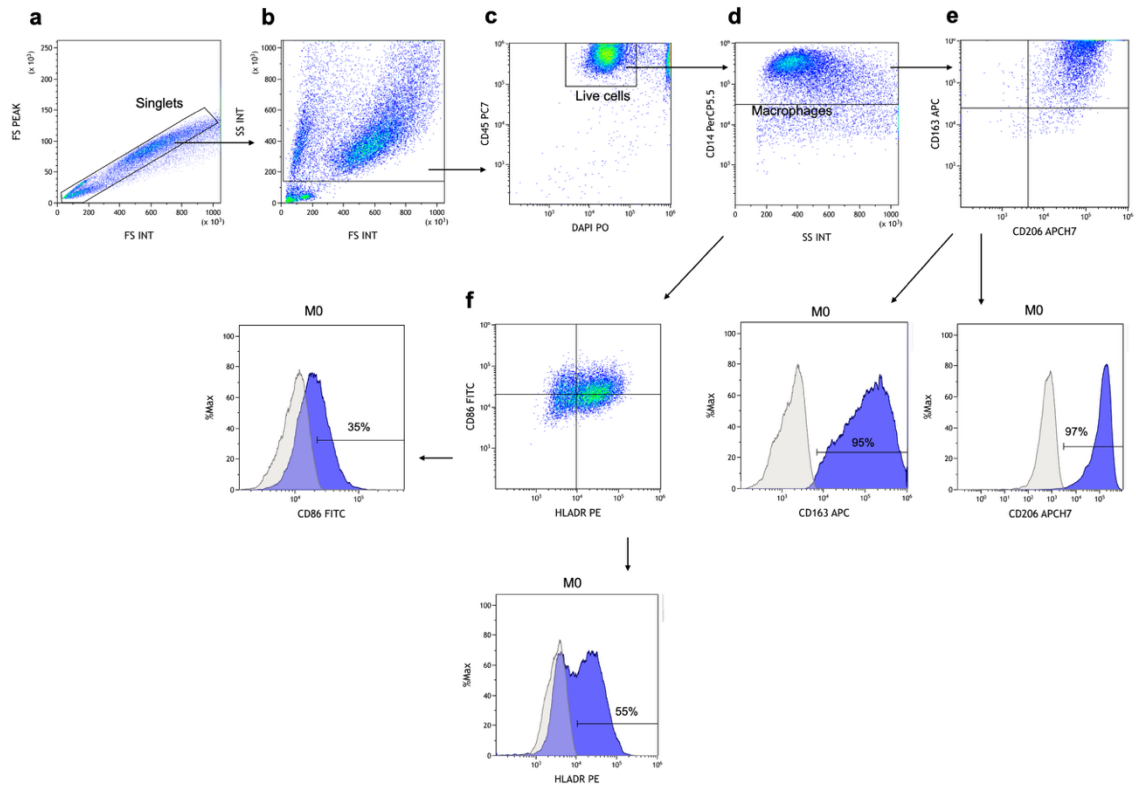

**Supplementary Fig. S4. Flow cytometric gating strategy for macrophage phenotype identification and evaluation of MFI and % for CD86, HLA-DR, CD206 and CD163 expression.** Cellular aggregates (a) and cellular debris (b) were excluded based on forward scatter (FS) and side scatter (SS) characteristics. (c) Viable leukocytes were identified by applying a CD45<sup>+</sup>DAPI<sup>-</sup> gate, effectively excluding both dead DAPI<sup>+</sup> cells and CD45<sup>+</sup> tumour cells. (d) A CD14<sup>+</sup> cell gate was selected to identify only the macrophage population. Within the CD14<sup>+</sup> cells gate, the MFI and % of cells positive for CD206, CD163 (e), CD86, and HLA-DR (f) were determined to characterise a pro-inflammatory (M1-like) or anti-inflammatory (M2-like) phenotype of macrophages. Isotype controls were used as negative controls.

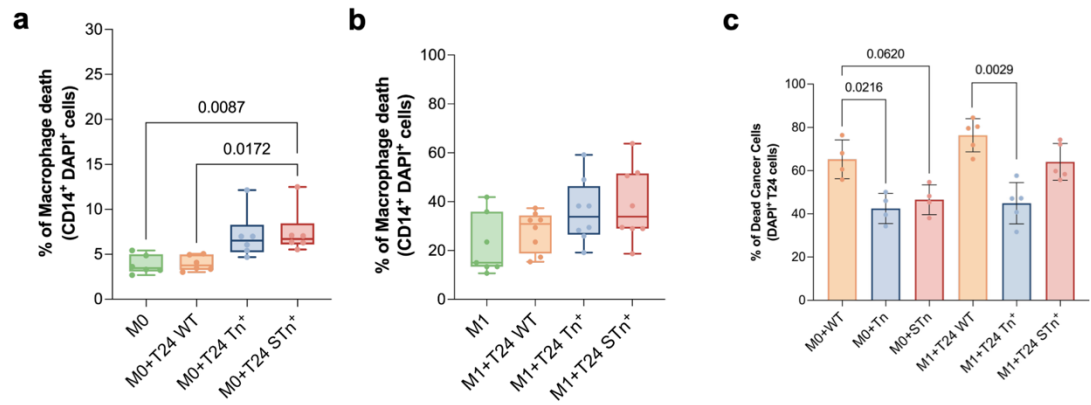

**Supplementary Fig. S5. Tn/sTn expression compromises macrophage survival and protects T24 cells from macrophage-mediated killing.** **a** *C1GALT1*-knockout (Tn<sup>+</sup>) and *C1GALT1*-knockout/*ST6GALNAC1*-knock-in (sTn<sup>+</sup>) T24 cells increased the death of unpolarized M0 macrophages, measured as DAPI positivity among CD14<sup>+</sup> cells (n=6). **b** Neither glycoengineered model significantly altered the death of IFN-γ/LPS-polarized M1 macrophages (n=8). **c** Tn/sTn-positive T24 cells showed lower death than WT cells following co-culture with either M0 or M1 macrophages (n=5). These findings indicate that truncated tumour O-glycans selectively impair M0 macrophage survival while conferring resistance to macrophage-mediated tumour-cell killing. Kruskal–Wallis followed by Dunn's multiple-comparisons test. Exact *P*-values are shown.

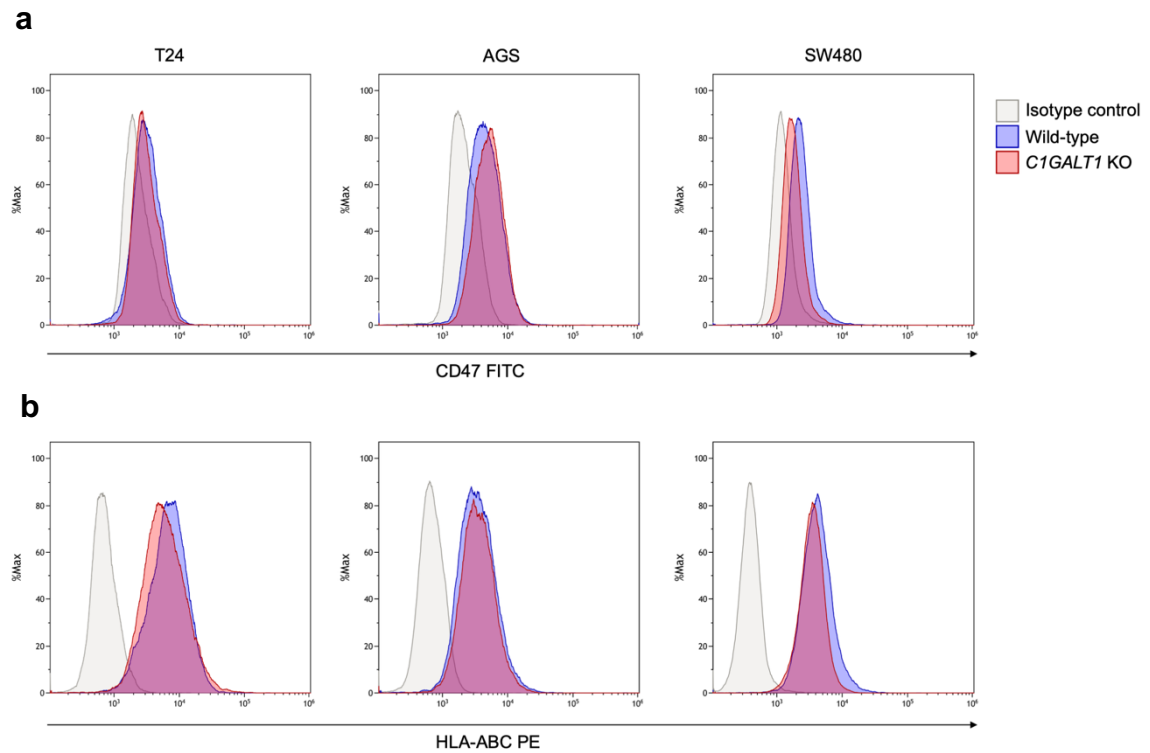

**Supplementary Fig. S6. Surface expression of CD47 and HLA class I is preserved following *C1GALT1* KO.** Representative flow cytometry histograms showing surface expression of CD47 (a) and HLA-ABC (b) on wild-type and *C1GALT1* KO T24, AGS and SW480 cells. Isotype controls are shown in grey.

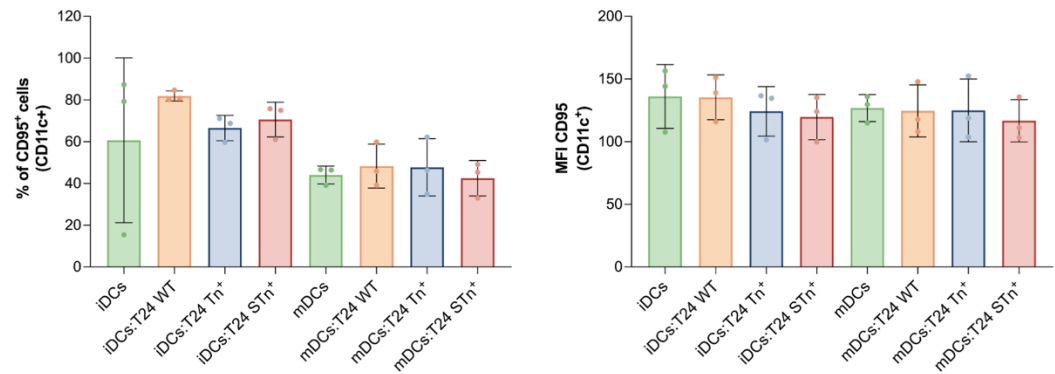

**Supplementary Fig. S7. Increased CD95 expression does not account for dendritic-cell death induced by Tn/sTn-positive T24 cells.** Co-culture with wild-type, *CIGALT1*-knockout (Tn<sup>+</sup>) or *CIGALT1*-knockout/*ST6GALNAC1*-knock-in (sTn<sup>+</sup>) T24 cells did not alter either the percentage of CD95<sup>+</sup> cells or CD95 mean fluorescence intensity within CD11c<sup>+</sup> immature or mature DCs. These results indicate that the increased DC death induced by Tn/sTn-positive tumour cells is not explained by increased CD95 abundance. Bars show means ± SD; points represent independent donors (n=3). Kruskal–Wallis followed by Dunn's multiple-comparisons test.

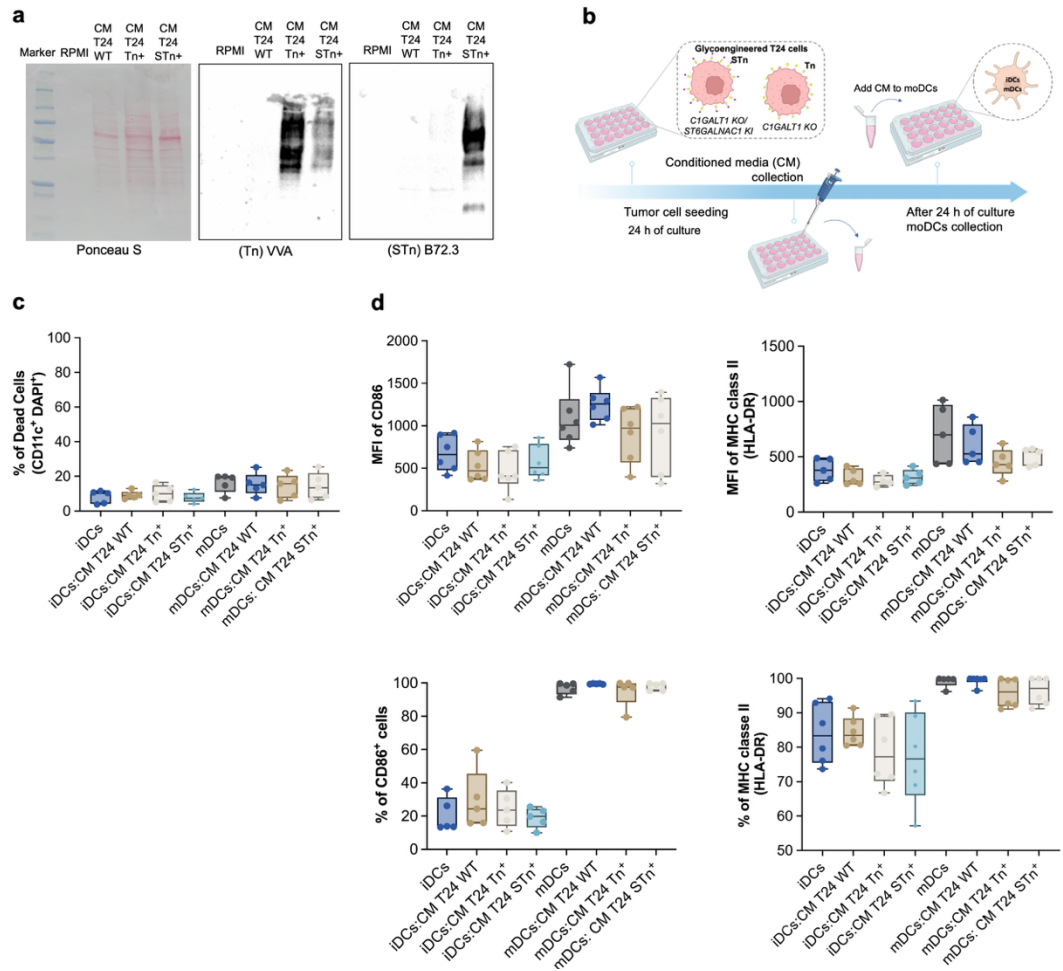

**Supplementary Fig. S8. Conditioned medium from Tn/sTn-positive T24 cells does not induce dendritic-cell death or dysfunction.** **a** Conditioned medium (CM) from both *C1GALT1* KO (Tn<sup>+</sup>) and *C1GALT1* KO/*ST6GALNAC1* KI (sTn<sup>+</sup>) T24 cells showed VVA reactivity. B72.3 reactivity was restricted to conditioned medium from the *ST6GALNAC1* KI model. Ponceau S shows total protein. **b** Experimental design for testing whether soluble factors released by glycoengineered T24 cells were sufficient to induce moDC death or alter maturation. Cell-free CM was collected after 24 h of tumour-cell culture and applied to immature or mature DCs for 24 h. **c** CM from Tn- or sTn-positive cells did not increase moDC death, measured as DAPI positivity among CD11c<sup>+</sup> cells. **d** CM also did not alter CD86 or HLA-DR expression; points represent independent donors (n=6). These results indicate that soluble tumour glycoproteins do not reproduce the effects of direct co-culture, supporting a contact-dependent mechanism.

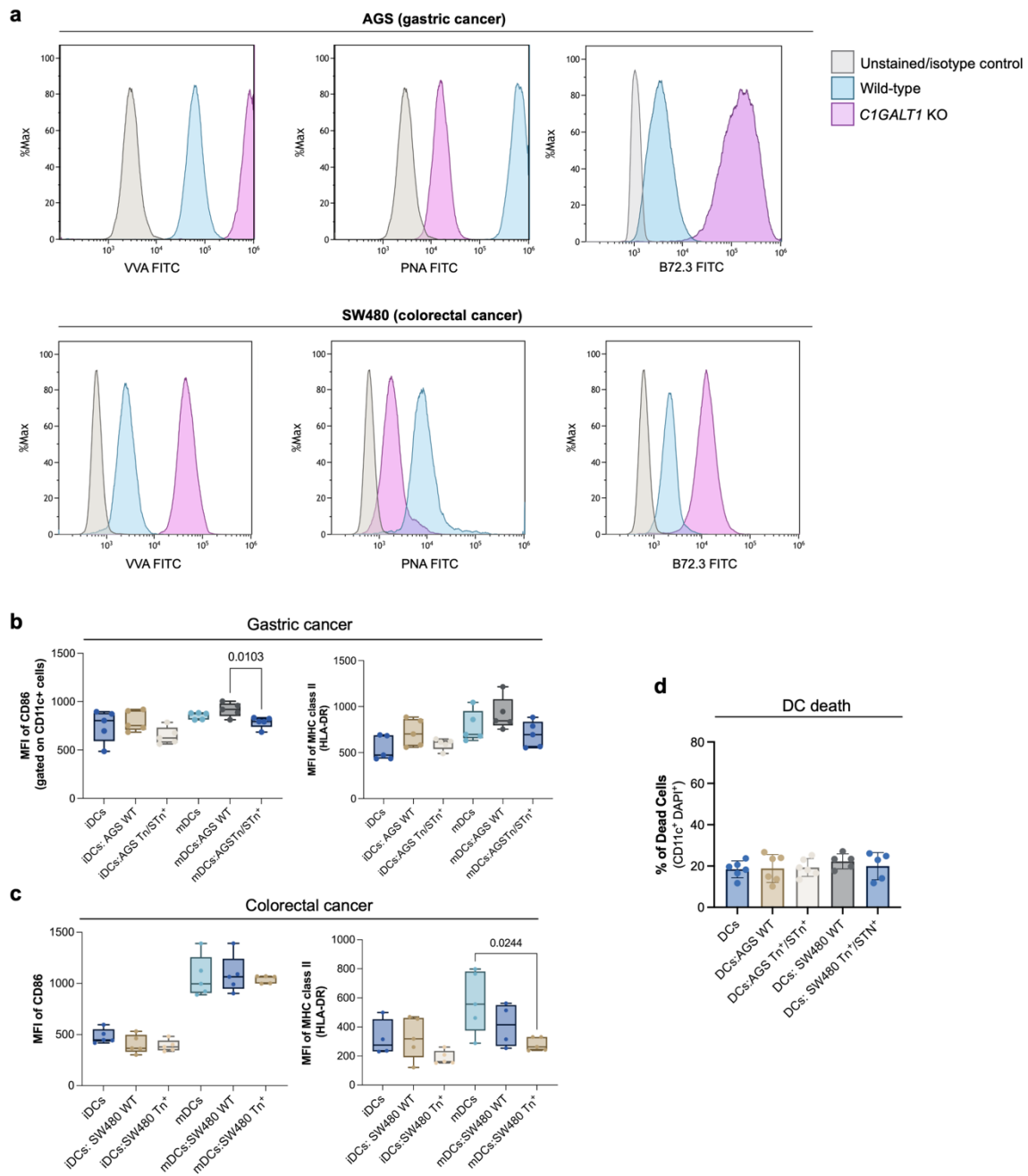

**Supplementary Fig. S9. C1GALT1 knockout in AGS and SW480 cells induces a truncated *O*-glycan phenotype and impairs dendritic cell maturation without affecting DC viability.**

**a** Representative flow cytometry histograms showing increased VVA and B72.3 binding and reduced PNA binding in *C1GALT1* KO AGS (gastric cancer) and SW480 (colorectal cancer) cells relative to wild-type controls, consistent with the accumulation of truncated *O*-glycans. Unstained/isotype controls are shown in grey. **b, c** Surface expression of CD86 and HLA-DR on immature (iDCs) and mature dendritic cells (mDCs) after co-culture with AGS (**b**) or SW480 (**c**) wild-type or *C1GALT1* KO cells. **d** DC viability after co-culture with AGS or SW480 wild-type or *C1GALT1* KO cells. Individual points represent independent biological replicates ($n=5$). Statistical analyses were performed using the Kruskal–Wallis test followed by Dunn's multiple-comparisons test. Exact *P* values are shown.

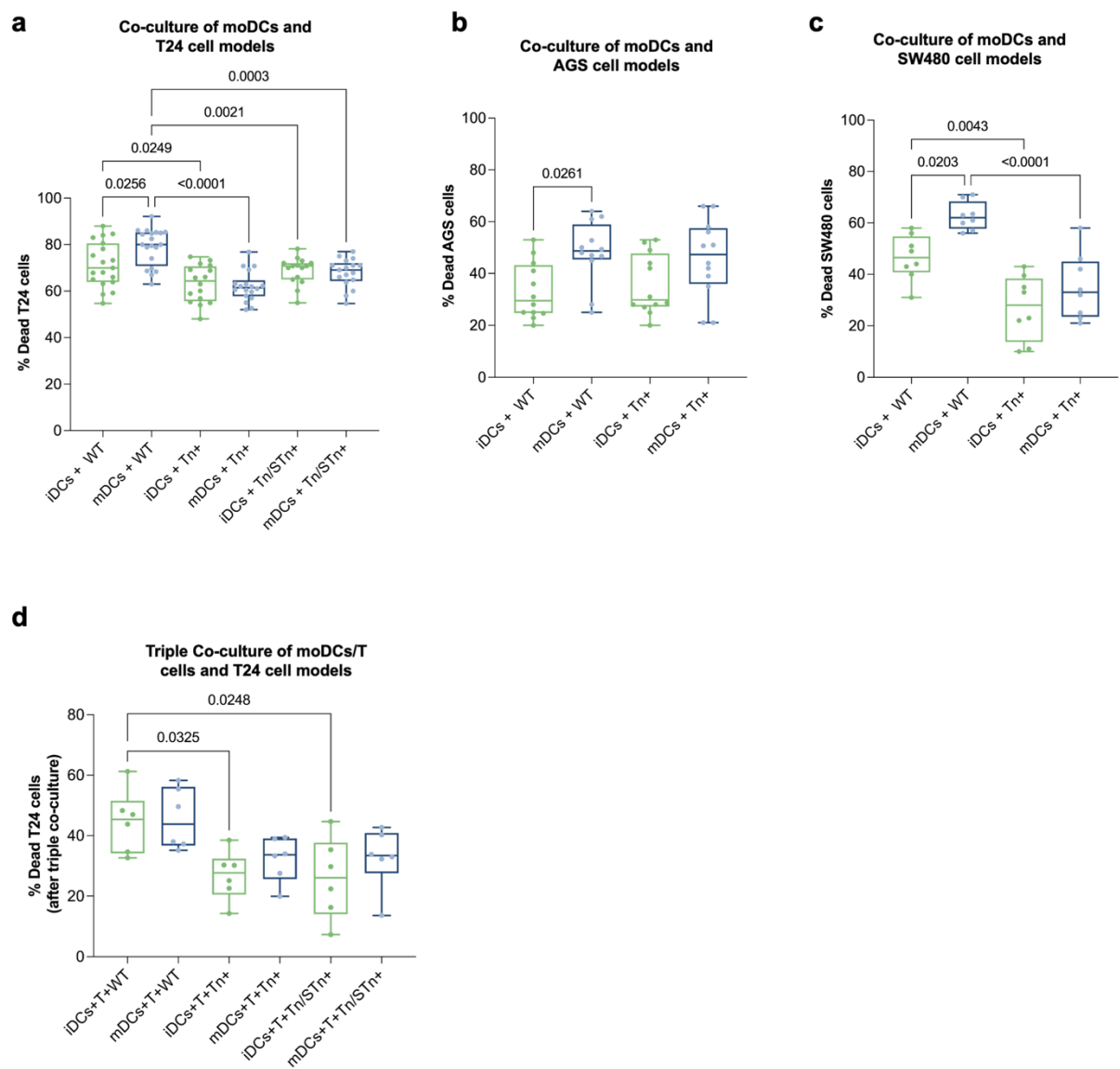

**Supplementary Fig. S10. Truncated *O*-glycans reduce dendritic cell-mediated tumour cell killing.** **a** Percentage of dead T24 wild-type, Tn+ and Tn/sTn+ cells following co-culture with iDCs or mDCs (n=12). **b,c** Percentage of dead AGS (n=12) (**b**) and SW480 (n=8) (**c**) wild-type and Tn+ cells following co-culture with immature or mature moDCs. **d** Percentage of dead T24 wild-type, Tn+ and Tn/sTn+ cells following triple co-culture with moDCs and T cells (n=6). Ordinary one-way ANOVA followed by Dunnett's multiple-comparisons test. Exact *P* values are shown.

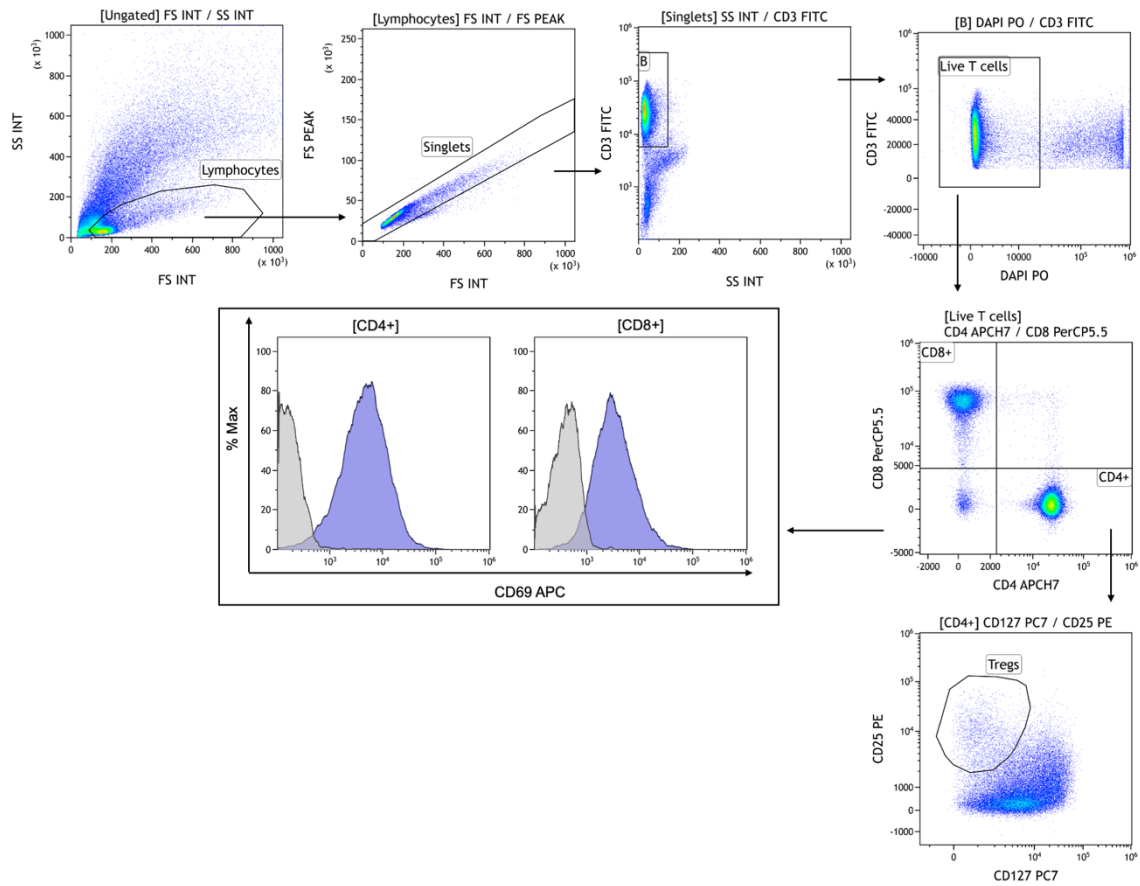

**Supplementary Fig. S11. Flow cytometry gating strategy used for the analysis of T-cells following triple co-culture with tumour cells and monocyte-derived dendritic cells (moDCs).** Lymphocytes were identified by forward and side scatter, followed by singlet discrimination and gating on live (DAPI<sup>-</sup>) CD3<sup>+</sup> T-cells. CD4<sup>+</sup> and CD8<sup>+</sup> T-cell subsets were identified for CD69 analysis, and Tregs were defined as CD25<sup>high</sup>CD127<sup>low/neg</sup> cells within the CD4<sup>+</sup> population. Representative plots are shown.

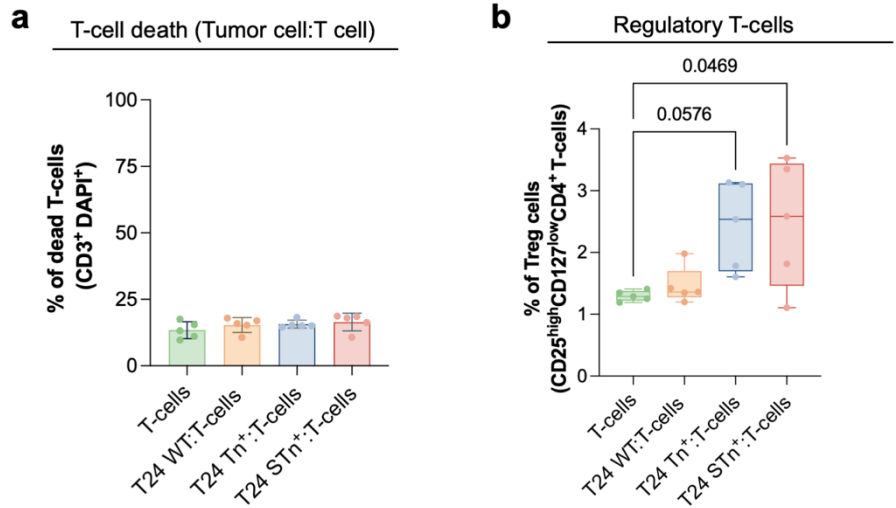

**Supplementary Fig. S12. T-cell death and regulatory T-cell frequency following co-culture with T24 glycoengineered cell models.** **a** Percentage of dead CD3<sup>+</sup> T cells following co-culture with wild-type, Tn<sup>+</sup> or Tn/sTn<sup>+</sup> T24 cells. **b** Frequency of regulatory T cells (Tregs; CD25<sup>high</sup>CD127<sup>low/neg</sup> of CD4<sup>+</sup> T cells) following co-culture with wild-type, Tn<sup>+</sup> or Tn/sTn<sup>+</sup> T24 cells. Individual points represent independent biological replicates (n=5). Statistical analyses were performed using ordinary one-way ANOVA followed by Dunnett's multiple-comparisons test. Exact *P* values are shown.

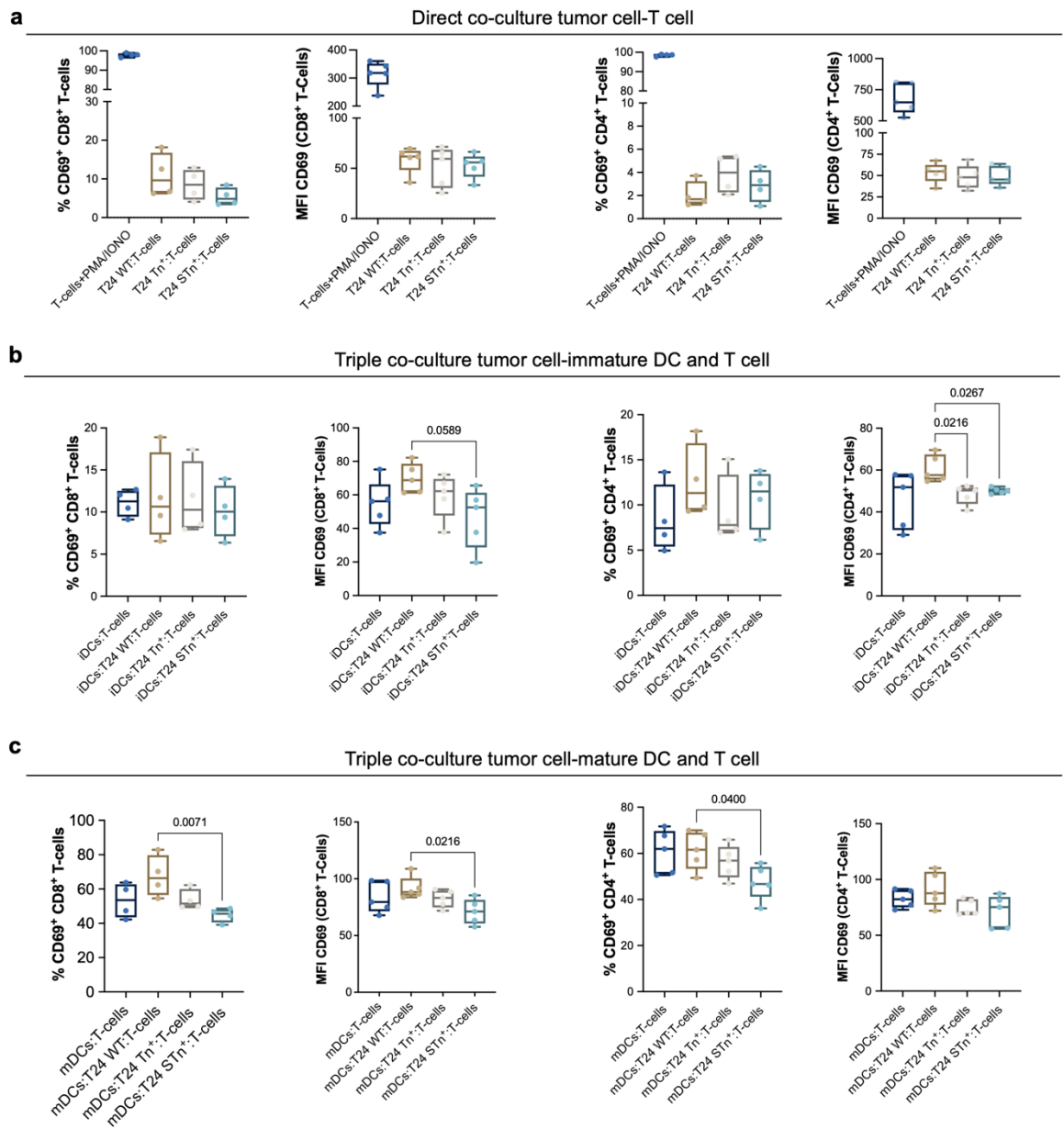

**Supplementary Fig. S13. T-cell activation following direct and triple co-culture with T24 glycoengineered cell models and moDCs.** **a** Frequency of CD69<sup>+</sup> CD8<sup>+</sup> and CD4<sup>+</sup> T cells and CD69 mean fluorescence intensity (MFI) following direct co-culture with wild-type, Tn<sup>+</sup> or Tn/sTn<sup>+</sup> T24 cells. PMA/ionomycin-stimulated T cells were used as a positive control. **b,c** Frequency of CD69<sup>+</sup> CD8<sup>+</sup> and CD4<sup>+</sup> T cells and CD69 MFI following triple co-culture with pre-primed iDCs (**b**) or mDCs (**c**) with T24 wild-type, Tn<sup>+</sup> or Tn/sTn<sup>+</sup> cells. Individual points represent independent biological replicates (n=5). Statistical analyses were performed using ordinary one-way ANOVA followed by Dunnett's multiple-comparisons test. Exact *P* values are shown.

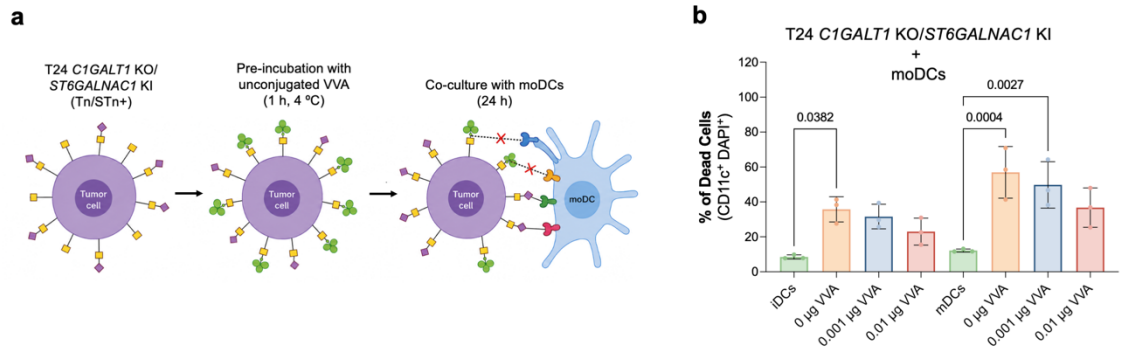

**Supplementary Fig. S14. VVA binding partially restores moDC-mediated killing of Tn/sTn+ tumour cells.** **a** Experimental workflow. T24 *C1GALT1* KO/*ST6GALNAC1* KI (Tn/sTn+) cells were pre-incubated with unconjugated VVA (1 h, 4 °C), washed and co-cultured with iDCs or mDCs for 24 h. **b** Percentage of dead tumour cells following co-culture with iDCs or mDCs after pre-incubation with VVA (0.001 µg and 0.01 µg per well). Individual points represent independent biological replicates (n=3). Statistical analyses were performed using the Friedman test followed by Dunn's multiple-comparisons test. Exact *P* values are shown.

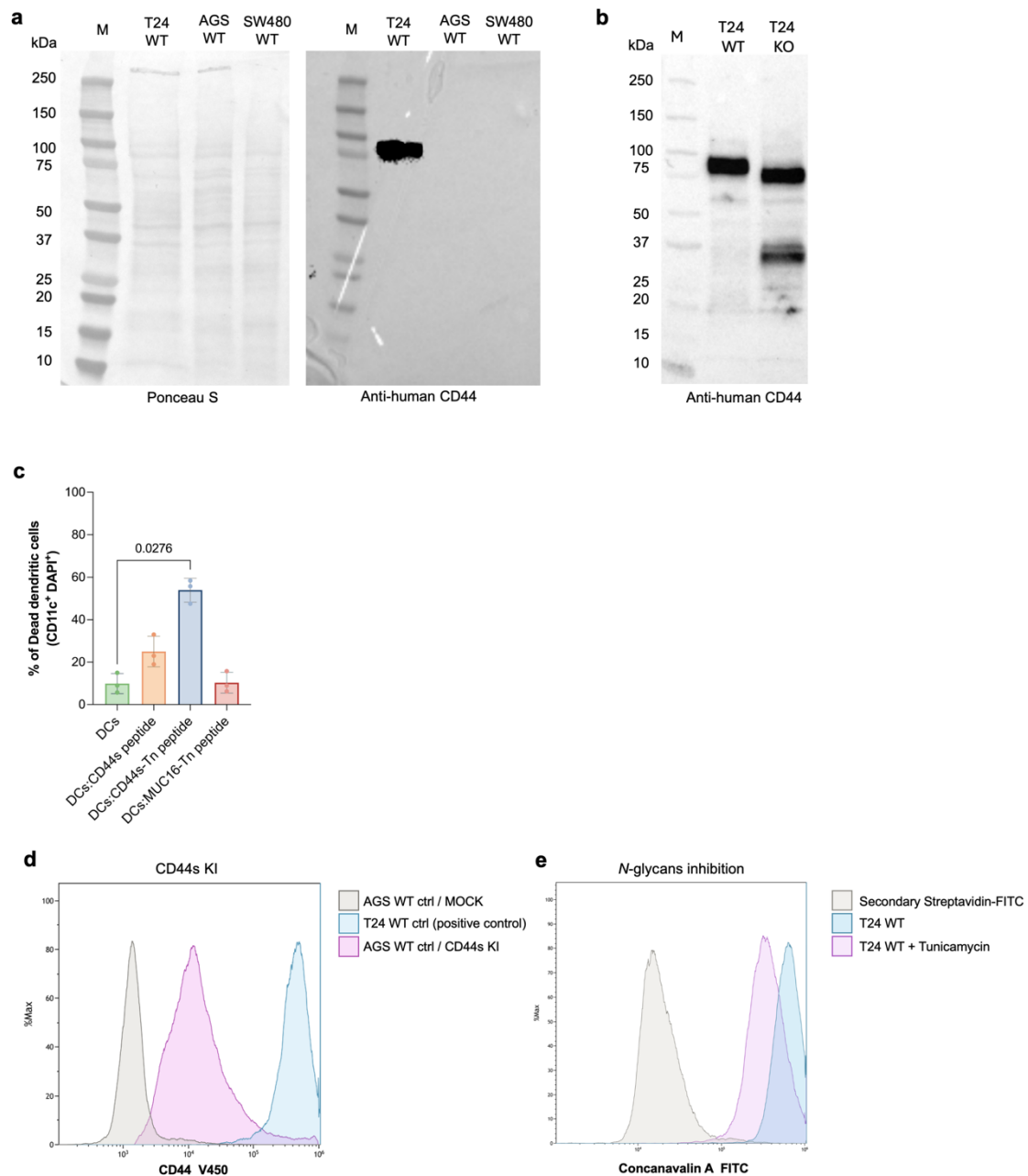

**Supplementary Fig. S15. Assessment of CD44 expression.** **a** Ponceau S staining and immunoblot analysis of CD44 expression in wild-type T24, AGS and SW480 cells. **b** Immunoblot validation of CD44 expression in T24 WT and *C1GALT1* KO cells. **c** Viability of immature dendritic cells (iDCs) following stimulation with recombinant 30-mer non-glycosylated CD44s peptide, Tn-glycosylated CD44 peptide or a MUC16 Tn-glycopeptide. Individual points represent independent biological replicates (n=3). Statistical analyses were performed using Friedman test followed by Dunn's multiple-comparisons test. Exact *P* values are shown. **d** Representative flow cytometry histograms showing surface CD44 expression in AGS control (mock) and CD44s KI AGS cells, with T24 wild-type cells included as a positive control. **e** Representative flow cytometry histograms showing Concanavalin A binding to untreated and tunicamycin-treated T24 cells, confirming inhibition of *N*-glycosylation.

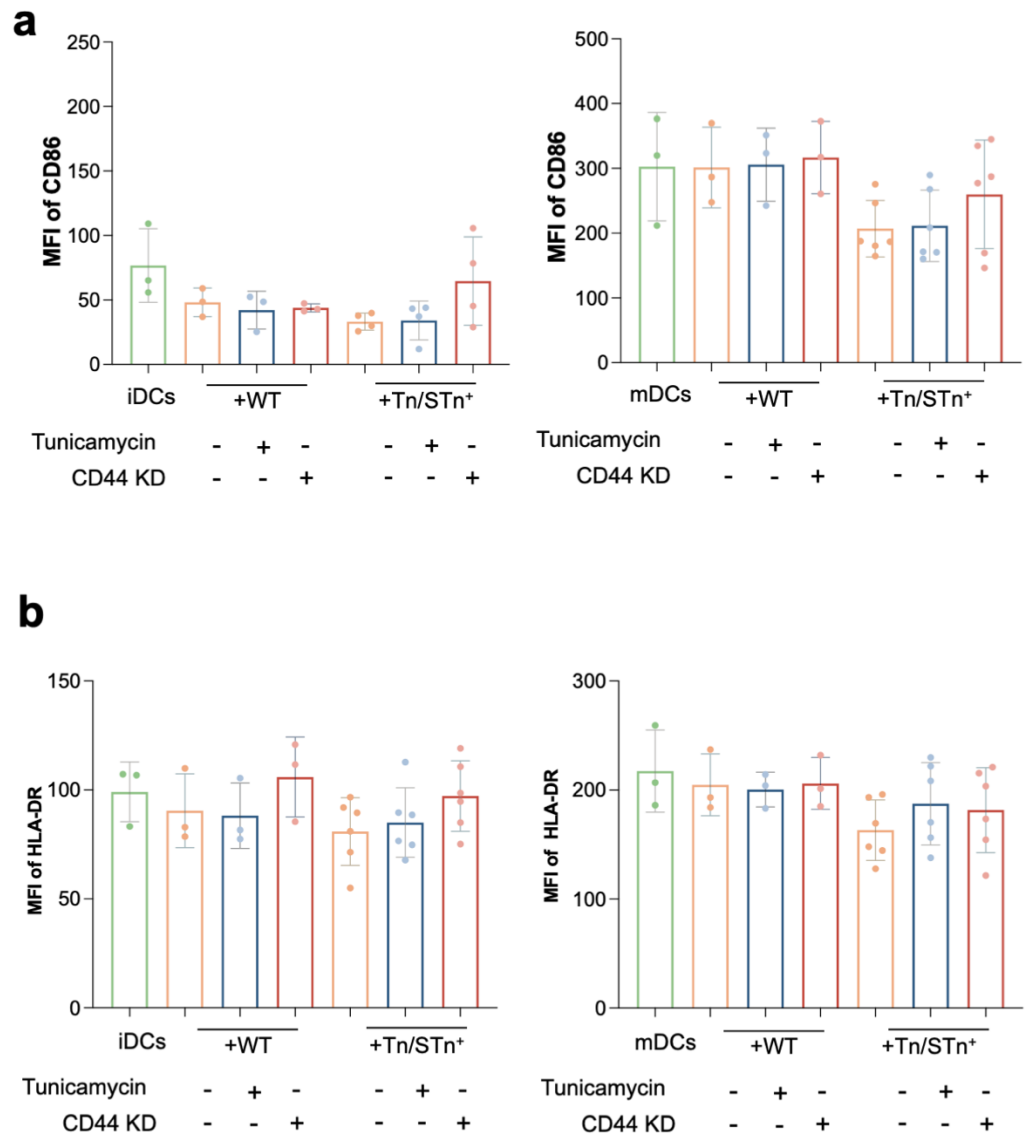

**Supplementary Fig. S16. Surface expression of CD86 and HLA-DR on CD86<sup>+</sup> and HLA-DR<sup>+</sup> dendritic cells following inhibition of *N*-glycosylation and/or CD44 knockdown. a,b** Mean fluorescence intensity (MFI) of CD86 (a) and HLA-DR (b) within the CD86<sup>+</sup> and HLA-DR<sup>+</sup> dendritic cell populations, respectively, following co-culture with wild-type or Tn/sTn<sup>+</sup> T24 cells, untreated or treated with tunicamycin, or following CD44 knockdown.

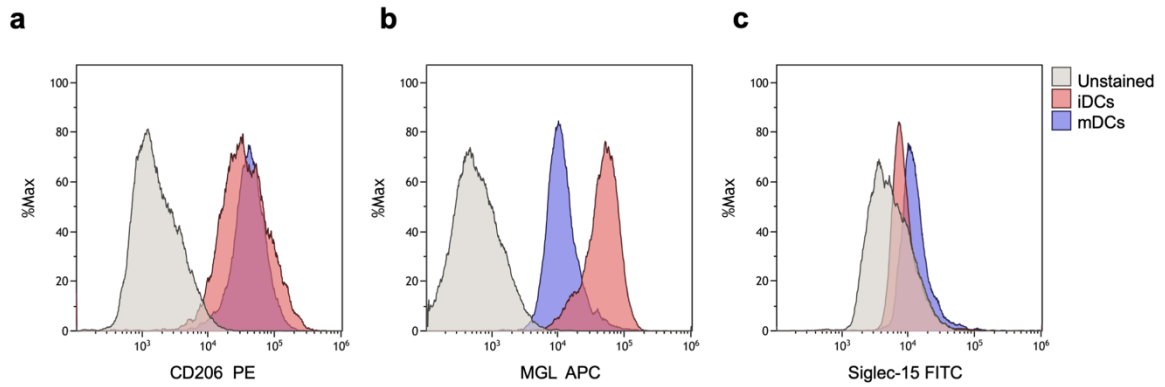

**Supplementary Fig. S17. Phenotypic characterization of immature and mature monocyte-derived dendritic cells.** **a–c** Representative flow cytometry histograms showing surface expression of CD206 (**a**), MGL (**b**) and Siglec-15 (**c**) on iDCs and mDCs. CD206 was highly expressed on both iDCs and mDCs, whereas MGL expression decreased following DC maturation. Little or no Siglec-15 surface expression was detected in either population. Unstained controls are shown in grey.

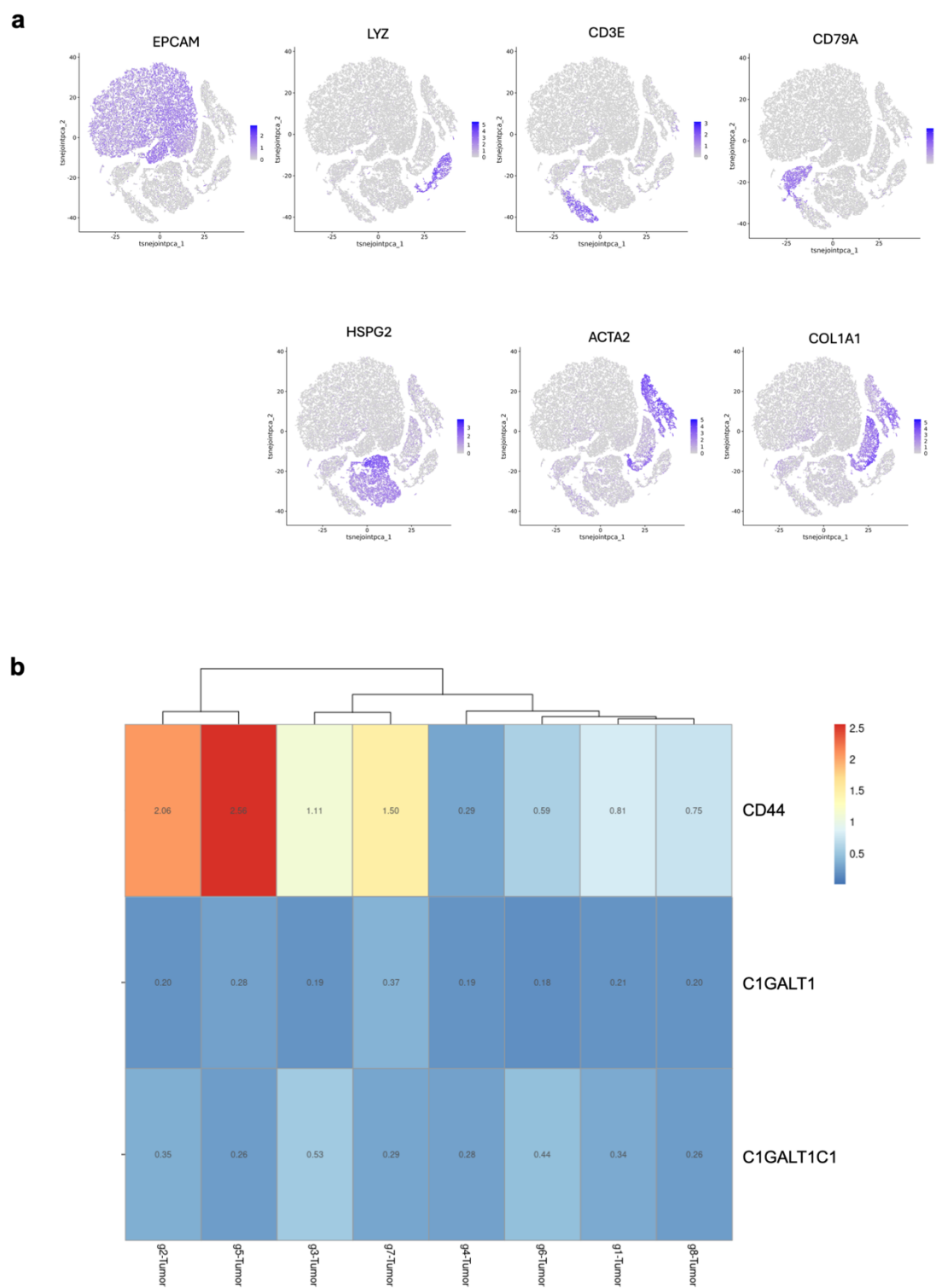

**Supplementary Fig. S18. Cell-type annotation and expression of CD44 and *O*-glycosylation-related genes in the scRNA-seq dataset (GSA-Human under the accession code HRA000212).** **a** UMAP feature plots showing the expression of canonical marker genes used for cell-type annotation, including epithelial (EPCAM), myeloid (LYZ), T-cell (CD3E), B-cell (CD79A) and endothelial (HSPG2), mCAFs (ACTA2) and iCAFs (COL1A1) markers. **b** Heatmap showing the average expression of CD44, C1GALT1 and C1GALT1C1 across tumour cell clusters.

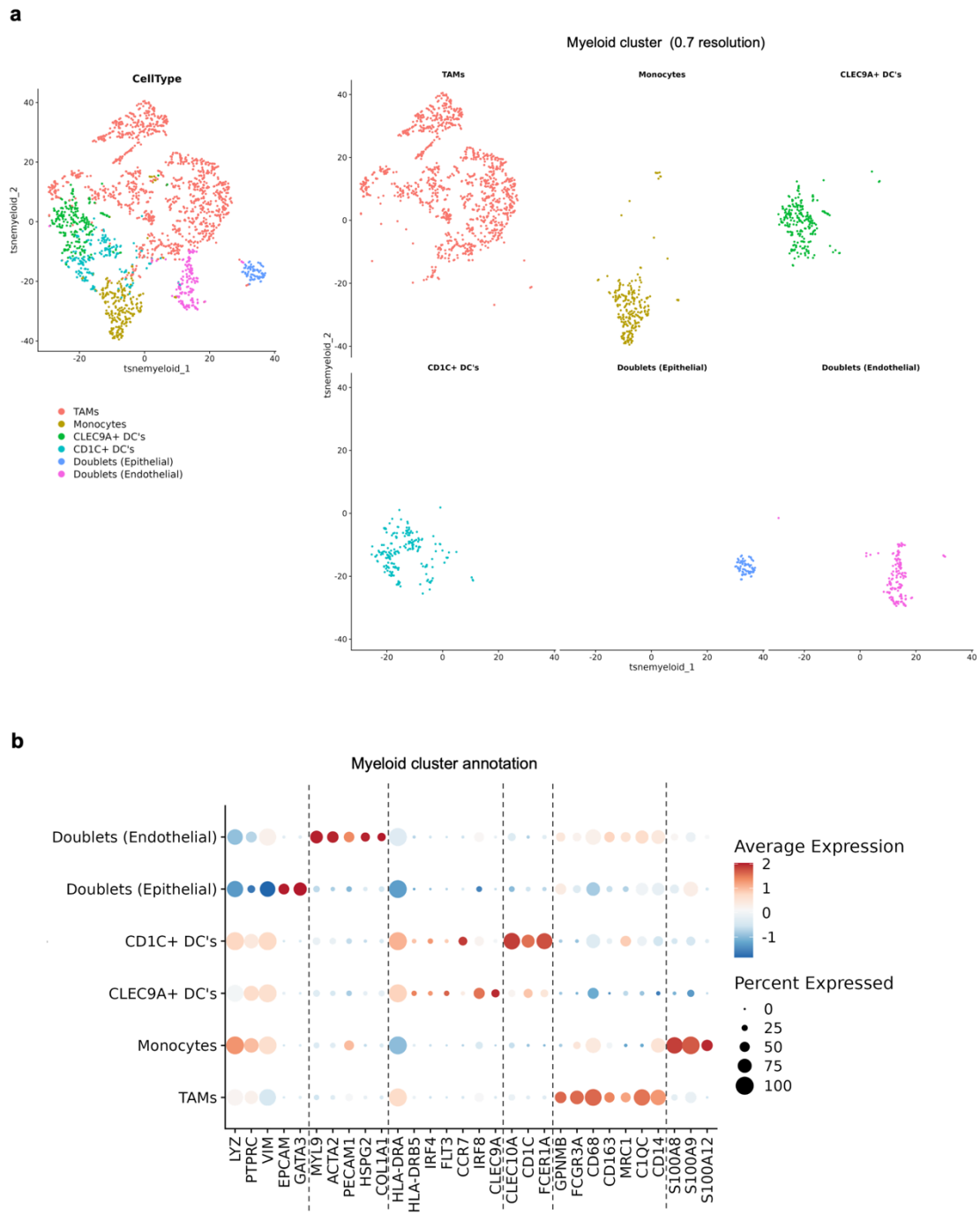

**Supplementary Fig. S19. Annotation of myeloid cell populations in the bladder cancer scRNA-seq dataset.** **a** UMAP representation of the myeloid compartment showing cell populations annotated based on canonical lineage markers. **b** Dot plot showing the expression of canonical marker genes used for cluster annotation. Dot size indicates the percentage of expressing cells and colour indicates the average scaled expression.

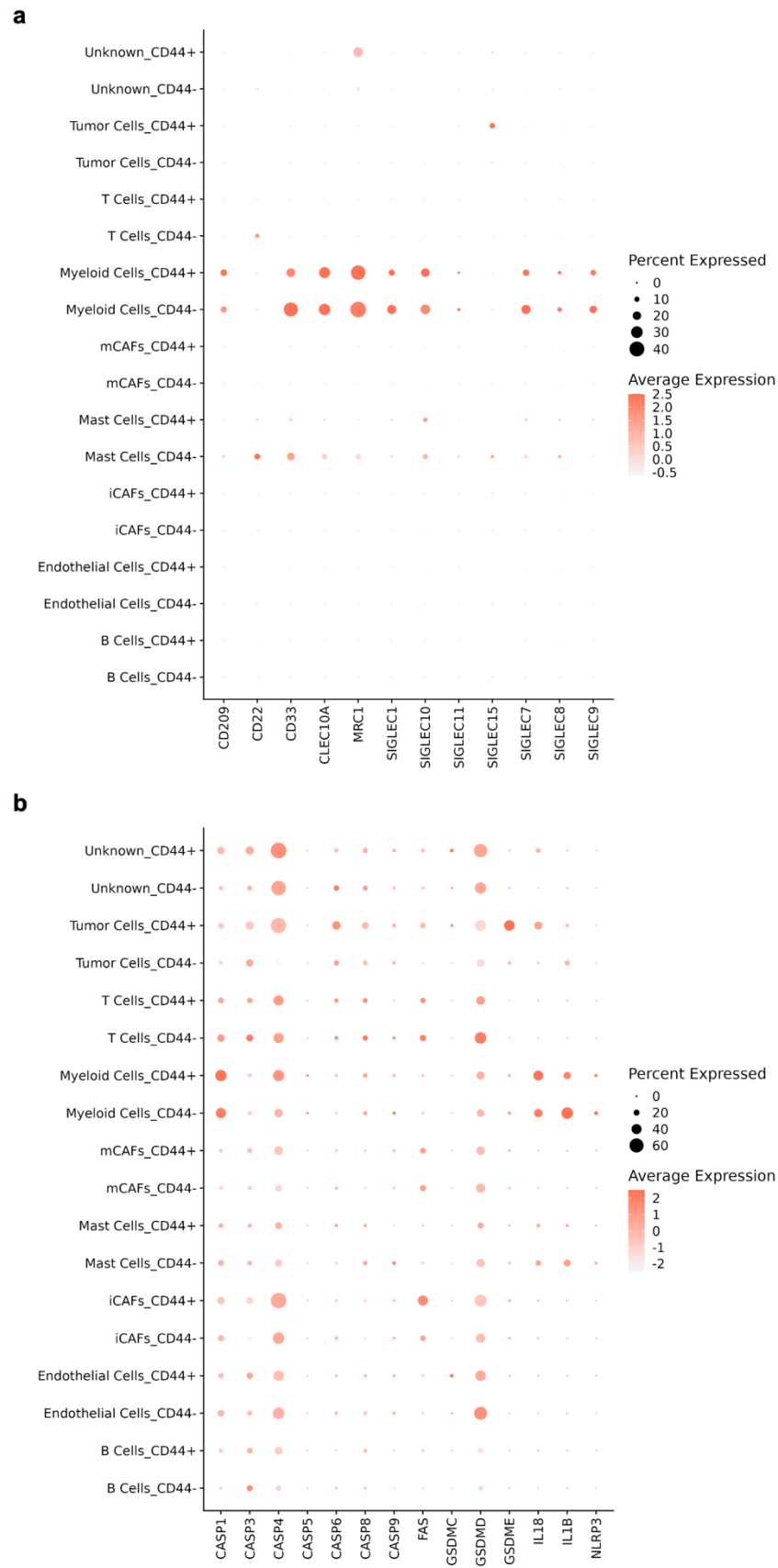

**Supplementary Fig. S20. Expression of glycan-binding receptors and pyroptosis-related genes across cell populations stratified by tumour cell CD44 expression. a,b** Dot plots showing the expression of glycan-binding receptors (a) and pyroptosis-related genes

**(b)** across annotated cell populations. Samples were stratified into CD44<sup>+</sup> and CD44<sup>−</sup> groups according to CD44 expression in the tumour cell cluster. Dot size indicates the percentage of expressing cells and colour indicates the average scaled expression.

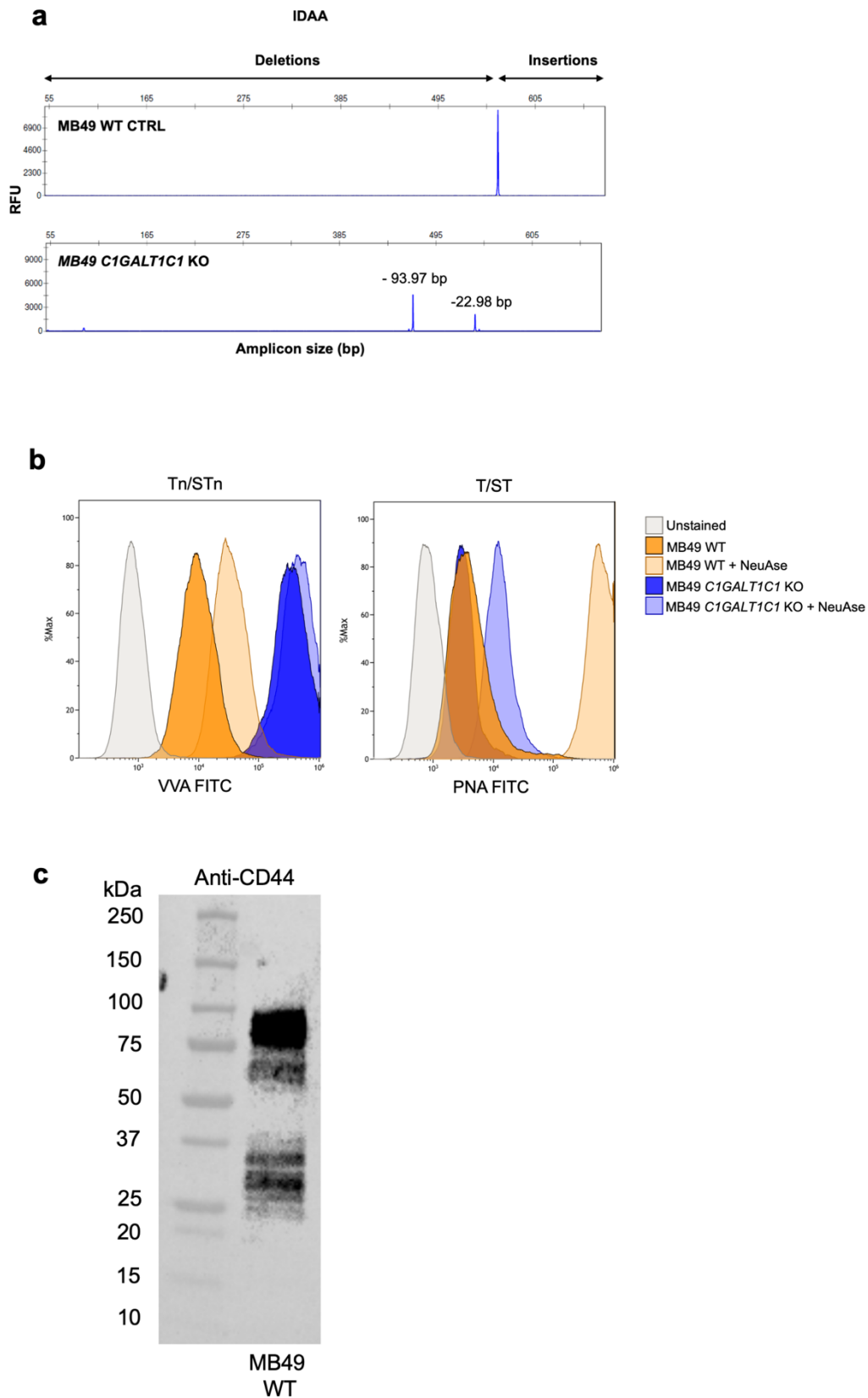

**Supplementary Fig. S21. Generation and validation of the MB49 *C1GALT1C1* knockout model.** **a** IDAA analysis confirming *C1GALT1C1* gene editing in MB49 cells. The wild-type control showed the expected amplicon size, whereas the *C1GALT1C1* knockout clone exhibited deletion alleles of the indicated sizes. **b** Representative flow cytometry histograms showing

increased VVA binding and reduced PNA binding in MB49 *C1GALT1C1* KO compared with wild-type cells. Neuraminidase treatment was used to evaluate the contribution of sialylation to the observed glycophenotype. Unstained controls are shown in grey. **c** Immunoblot showing CD44 expression in wild-type MB49 cells.

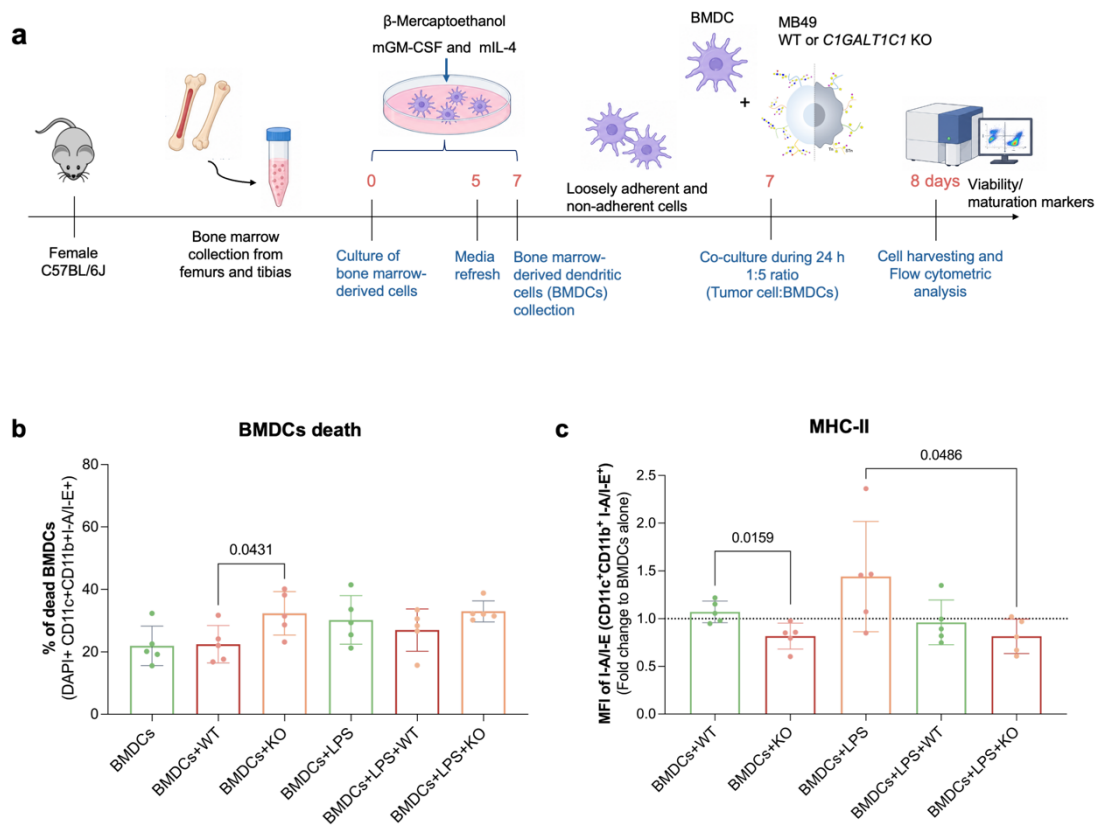

**Supplementary Fig. S22. Co-culture of MB49 bladder cancer cells with mouse bone marrow-derived dendritic cells (BMDCs).** **a** Bone marrow cells were isolated from C57BL/6 mice and differentiated into BMDCs for 7 days in the presence of murine IL-4 and GM-CSF. BMDCs were then co-cultured with MB49 WT or *C1GALT1C1* KO cells for 24 h in the presence or absence of LPS. **b** BMDC cell death was assessed by flow cytometry and quantified as the percentage of DAPI-positive, CD11c<sup>+</sup>CD11b<sup>+</sup>I-A/I-E<sup>+</sup> cells. **c** Surface MHC-II (I-A/I-E) expression on BMDCs was evaluated and expressed as the fold change in mean fluorescence intensity (MFI) relative to BMDCs cultured alone. BMDCs were generated independently from five mice (n=5). Statistical significance was determined using the Kruskal–Wallis test with Dunn's multiple-comparisons test; Exact *P* values are shown.

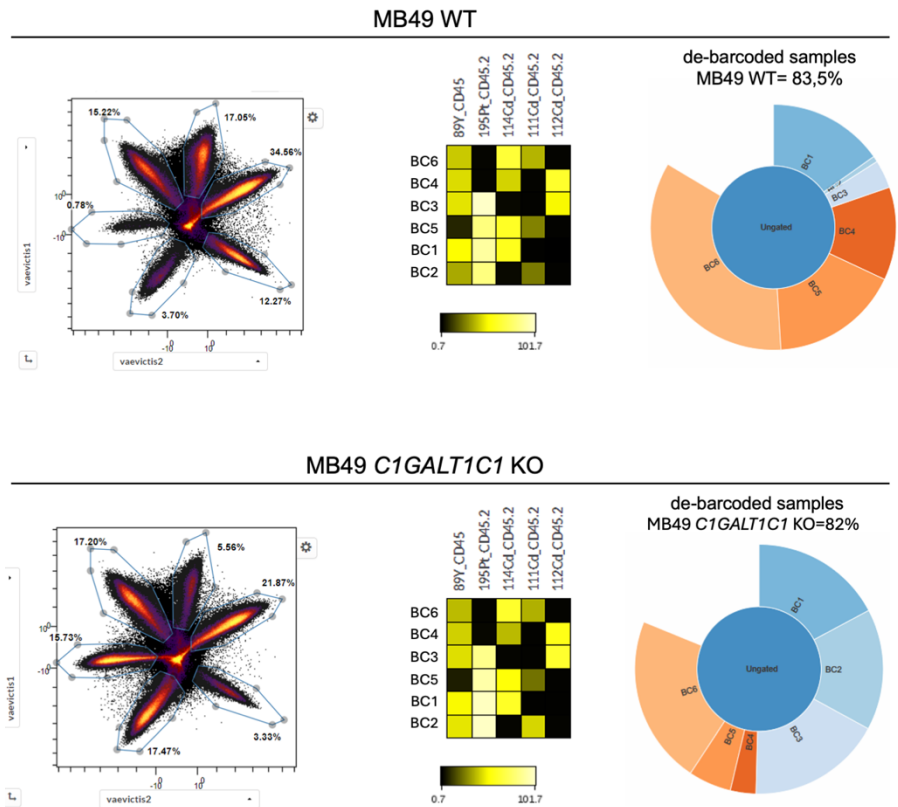

**Supplementary Fig. S23. CyTOF sample barcoding and debarcoding quality control.** Representative barcode separation, barcode signal intensity heatmaps and debarcoding results for tumours isolated from mice inoculated with wild-type (WT) or *C1GALT1C1* knockout (KO) MB49 cells and analysed by CyTOF. Pie charts indicate the proportion of successfully debarcoded events for each sample.

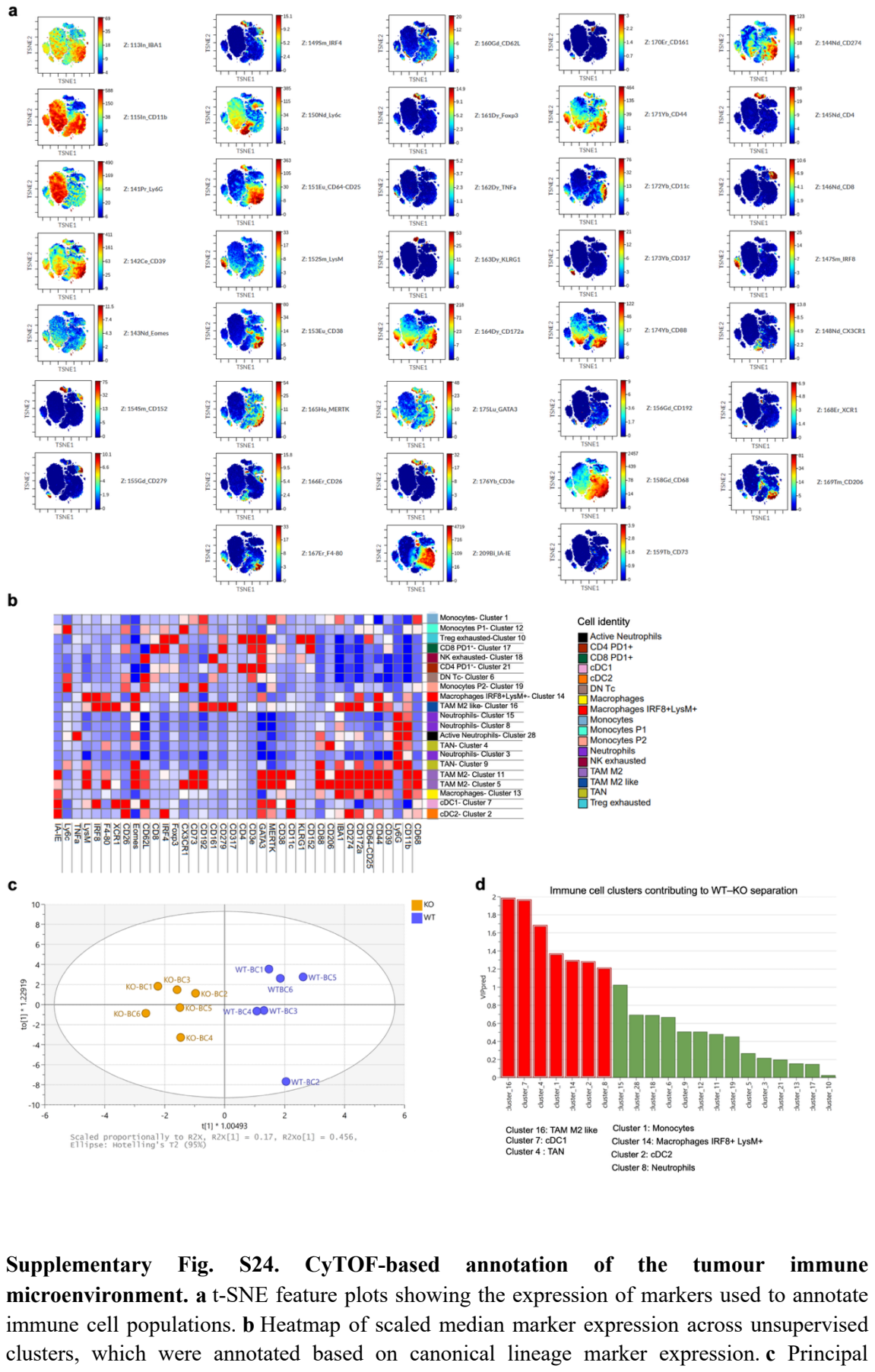

component analysis (PCA) of WT and *C1GALT1C1* KO MB49 tumours. Each point represents an individual tumour. **d** Variable importance in projection (VIP) scores ranking the immune cell clusters contributing to sample separation.

**a**

**Panel – Murine lymphoid gate strategy**

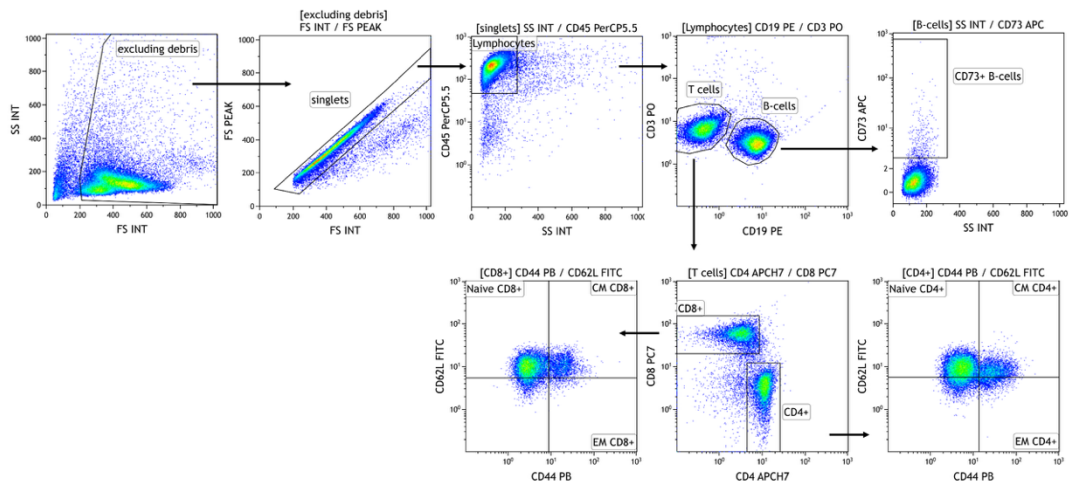

**b**

**Panel – Murine myeloid gate strategy**

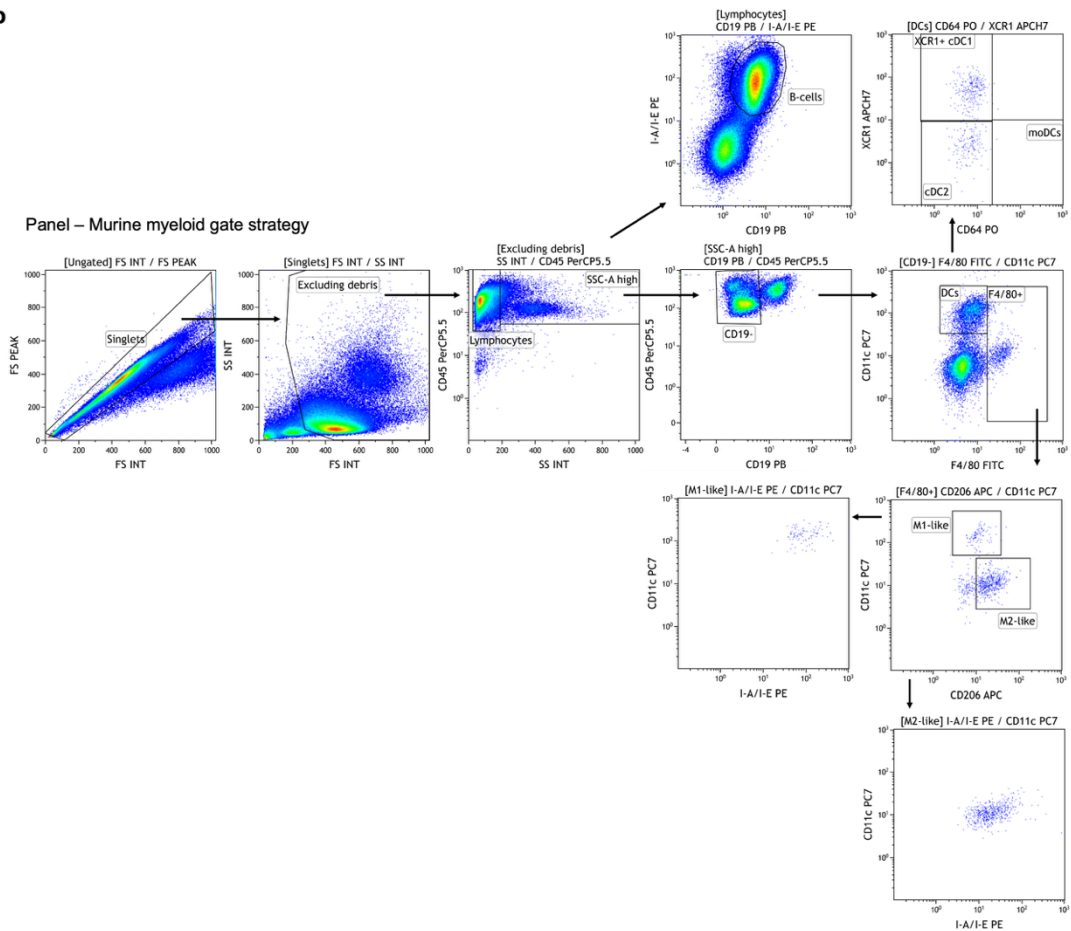

**Supplementary Fig. S25. Representative flow cytometry gating strategy for murine lymphoid and myeloid cell populations.** **a** Representative gating strategy used to identify lymphoid cell populations used for tumour-infiltrating immune cells, tumour-draining lymph nodes and spleen. After exclusion of debris and doublets, live CD45<sup>+</sup> leukocytes were gated to identify CD19<sup>+</sup> B cells and CD3<sup>+</sup> T cells. CD73 expression was assessed on B cells. CD4<sup>+</sup> and CD8<sup>+</sup> T-cell subsets were subsequently classified as naïve (CD44<sup>-</sup>CD62L<sup>+</sup>), central memory

(CM; CD44<sup>+</sup>CD62L<sup>+</sup>) or effector memory (EM; CD44<sup>+</sup>CD62L<sup>-</sup>). **b** Representative gating strategy used to identify myeloid cell populations. Following exclusion of debris, doublets and lymphocytes, CD45<sup>+</sup>SSC-A<sup>high</sup> cells were analysed to identify dendritic cells (CD11c<sup>+</sup>), including cDC1 (XCR1<sup>+</sup>) and cDC2 (XCR1<sup>-</sup>CD64<sup>-</sup>), and F4/80<sup>+</sup> macrophages, which were further classified as M1-like (CD11c<sup>+</sup> CD206<sup>-</sup> I-A/I-E<sup>bright</sup>) or M2-like (CD11c<sup>-</sup> CD206<sup>+</sup> I-A/I-E<sup>dim</sup>).
