## Supplemental Methods for "CD44-CD206 axis drives dendritic cell loss and immunosuppression in immaturely glycosylated bladder tumours"

### Materials and methods

#### Bulk RNA-sequencing analysis of TCGA data

Normalised gene expression data ($\log_2(\text{TPM} + 0.001)$) and corresponding clinical annotations, including overall survival data, for 412 patients with bladder cancer were obtained from The Cancer Genome Atlas (TCGA) through the UCSC Xena Browser (https://xenabrowser.net/). Only samples with complete clinical and expression data were included in downstream analyses.

The distribution of gene expression levels was assessed using the Shapiro–Wilk test. As most variables were not normally distributed, non-parametric methods were used. Comparisons between two independent groups were performed using the Wilcoxon rank-sum test. Pairwise associations between glycosyltransferases and immune cell-related markers were evaluated using Spearman’s rank correlation coefficient, as calculated using the `cor.test` function in R. For single pairwise comparisons, nominal *p*-values were reported. When multiple correlations were computed simultaneously (e.g., correlation matrices across gene panels), *p*-values were adjusted using the Benjamini–Hochberg false discovery rate (FDR) method.

Survival analyses were performed using the Kaplan–Meier method, and differences between groups were assessed with the log-rank test using the `survival` and `survminer` R packages. Patients were stratified into expression groups (high versus low) based on tertile cutoffs (upper versus lower tertiles). Hierarchical clustering was performed to explore gene expression patterns across defined signatures using Euclidean distance as the distance metric. Results were visualised using heatmaps generated with the `pheatmap` R package.

All statistical tests were two-sided, and a *p*-value $< 0.05$ was considered statistically significant unless otherwise stated. All statistical analyses were conducted using R software (version 4.2.1).

#### Human samples and ethics statement

Human samples were obtained in accordance with the principles of the Declaration of Helsinki. Monocytes were isolated from surplus buffy coats from healthy blood donors, provided by the Immunohemotherapy Department of Centro Hospitalar Universitário São João (CHUSJ), Porto, Portugal. Procedures were approved by the hospital ethics committee (protocol 90/19). Blood donors provided informed written consent that by-products of their blood donations could be used for research purposes. All procedures involving the collection and use of tumour tissues were performed with the patients’ written informed consent and after approval by the IPO Porto ethics committee (reference CES-IPOFG-EPE-86R1/017 for bladder cancer).

#### Immunohistochemistry

Formalin-fixed, paraffin-embedded (FFPE) muscle-invasive bladder cancer (MIBC) tissue samples obtained from the IPO Porto Biobank were first stained with B72.3+CC49 (0.5 $\mu\text{g/mL}$; Abcam) to classify the tissues as STn-positive or STn-negative. Subsequently, immunohistochemical analysis was performed using the CD68 (1:100; Dako) and CD163 (1:100; Dako) antibodies to detect macrophages, while CD45 (1:500; Cell Marque) staining was used to assess leukocyte infiltration within the tissue microenvironment. In brief, the FFPE tumour sections were deparaffinised in xylene, rehydrated and subjected to antigen retrieval for 20 min in boiling EDTA (pH 8) for CD163, CD68, and CD45 or in citrate (pH 6) for B72.3+CC49. Sections were then incubated with 3–4% hydrogen peroxide (Leica) for 5 min. After blocking with 5% bovine serum albumin (BSA) for 60 min and incubation with primary antibodies, staining was developed using the Novolink Max Polymer DS Kit (Leica), according to the manufacturer's instructions, followed by 3,3'-diaminobenzidine (DAB, Leica). Nuclei were counterstained with haematoxylin for 5 min. Sections were dehydrated, cleared in xylene and mounted with Entellan. The results were evaluated in a double-blind manner and reviewed by an experienced pathologist. In cases of disagreement, the slides were re-reviewed to reach a consensus. Images were acquired using a BA310 Trinocular microscope equipped with an integrated MOTICAM X5 Plus camera using Motic Images Plus 3.0 software (Motic).

### **Immunofluorescence**

Formalin-fixed paraffin-embedded MIBC tumours and the corresponding lymph node metastases were evaluated by immunofluorescence to detect activated dendritic cells using anti-human CD11c Alexa Fluor 647 (1:100; Abcam) and anti-human CD86 antibodies (1:50; Thermo Fisher Scientific). CD68 was detected using an Alexa Fluor 594-conjugated goat anti-rabbit secondary antibody (1:300; Thermo Fisher Scientific). Nuclei were counterstained with 4',6-diamidino-2-phenylindole dihydrochloride (DAPI; Thermo Fisher Scientific). Fluorescence images were acquired using a Leica DMI6000 FFW microscope and analysed using LAS X software (Leica). Additionally, FFPE sections of subcutaneous tumours obtained from the *in vivo* study using mice inoculated with MB49 WT or MB49 *C1GALT1C1* KO cells were analysed by immunofluorescence to evaluate CD44 and Tn antigen expression. Tn antigen expression was detected using FITC-conjugated *Vicia villosa* agglutinin (VVA-FITC; 40µg/mL; Vector), whereas CD44 was detected using an unconjugated rabbit anti-CD44 primary antibody (1:250; Abcam) followed by an Alexa Fluor 594-conjugated goat anti-rabbit secondary antibody. Nuclei were counterstained with DAPI. Fluorescence images were acquired using an EVOS M7000 microscope (Thermo Fisher Scientific) and analysed using Fiji. Colocalisation between CD44 and Tn signals was assessed using the Colocalization Threshold Test plugin in Fiji to generate pixel-intensity maps visualising the spatial overlap between the detected markers.

### **Cell lines and culture conditions**

Human T24, SW480 and AGS and murine MB49 cells were obtained from the American Type Culture Collection (ATCC, Manassas, VA, USA). Cells were cultured in RPMI 1640 with GlutaMAX™ (Gibco™, Thermo Fisher Scientific) supplemented with 10% fetal bovine serum (FBS) (Gibco™,

Thermo Fisher Scientific) and 1% penicillin/streptomycin (PenStrep) (10,000 units/mL penicillin; 10,000 µg/mL streptomycin; Gibco, Thermo Fisher Scientific) and incubated at 37 °C in a humidified atmosphere with 5% CO<sub>2</sub>. Cells were screened weekly and confirmed to be mycoplasma-free throughout the study.

#### **Glycoengineered cell models**

Two guide RNAs targeting human *CIGALT1* (gRNA1: GTAAAGCAGGGCTACATGAG, gRNA2: ACAACACTTTGTTACAACGC) were cloned into a CRISPR-Cas9-containing dual gRNA expression vector (pRP[2CRISPR]-Puro-hCas; VectorBuilder, Inc.). One microgram of mammalian CRISPR vector was transfected into wild-type human T24 (bladder cancer)<sup>1</sup>, AGS (gastric cancer) (ref) and SW480 (colorectal cancer)<sup>2</sup> cell lines using jetPRIME® transfection reagent according to the manufacturer's instructions.

For the murine MB49 bladder cancer cell line, two guide RNAs targeting murine *CIGALT1C1* (gRNA 1: GCGGTCTGCCTGAAATACGC, gRNA2: CGAAAGTAGTGGGGCGTGCA) were cloned into a CRISPR-Cas9-containing dual gRNA expression vector (pRP[2CRISPR]-Puro-hCas9-U6>mC1galt1c1[grna#1]-U6>mC1galt1c1[gRNA#375]; VectorBuilder, Inc.). One microgram of mammalian CRISPR vector was transfected into the wild-type MB49 cell line using jetPRIME® transfection reagent according to the manufacturer's instructions. Single clones were obtained by serial dilution in 96-well plates, and knockout (KO) clones were identified by Indel Detection by Amplicon Analysis (IDAA) using an ABI PRISM™ 3010 Genetic Analyzer (Thermo Fisher Scientific) and Sanger sequencing. One clone with distinct out-of-frame indels was selected. Clones carrying only silent mutations were used as phenotypic controls. IDAA results were analysed using Peak Scanner Software V1.0 (Thermo Fisher Scientific). Human *ST6GALNAC1* (hST6GALNAC1 [NM\_018414.5], VectorBuilder) expression was introduced into T24 *CIGALT1* KO cells by transfection with a conventional mammalian expression vector, using jetPRIME transfection reagent (PolyPlus Transfection, 114-01), according to the manufacturer's instructions. Successfully transfected cells were selected with puromycin (2 µg/mL; EMD Millipore, 540222). In parallel, a mock system containing a 300 bp stuffer ORF (VectorBuilder, VB191025-4255jjz) was established.

#### **CD44 knockdown in the glycoengineered T24 cell models**

CD44 knockdown in glycoengineered T24 cell models was performed using small interfering RNA (siRNA), optimised and described in Gaiteiro *et al*<sup>3</sup>. Gene silencing of CD44 was achieved using two distinct Silencer® Select siRNAs (Invitrogen), siRNA IDs s2681 (target exon 2) and s2682 (target exons 1 and 2). Silencer® Select Negative Control siRNA (4390843, Invitrogen) served as the negative control. Cells were detached and seeded into 24-well plates (1 × 10<sup>5</sup> cells/well) and transfected using Lipofectamine RNAiMAX (Invitrogen) according to the manufacturer's instructions. Cells were exposed to the transfection complexes for 72 h. All experiments were performed in duplicate. RT-PCR was conducted to confirm CD44 silencing.

#### CD44s knock-in in glycoengineered AGS cell models

The human CD44s isoform (pRP[Exp]-Puro-EF1A>hCD44[NM\_001001391.2]) was expressed in AGS cells by transfection with a conventional mammalian expression vector using jetPRIME® transfection reagent (Polyplus Transfection) according to the manufacturer's instructions. In parallel, a mock system containing a 300 bp stuffer ORF (VectorBuilder, VB191025-4255jjz) was established. Successfully transfected cells were maintained in medium containing puromycin (2 µg/mL; EMD Millipore).

#### RNA extraction from FFPE human and murine tissues

RNA was extracted from FFPE tumour tissues from MIBC patients, including tumours expressing the short-chain *O*-glycans Tn and STn ($n=14$) and tumours with negative or low expression of short-chain *O*-glycans ($n=12$). Tissues were classified based on staining positivity for Tn, detected with VVA lectin, and STn, detected with the B72.3+CC49 antibody. For murine samples, subcutaneous tumours were generated by inoculating female C57BL/6 J mice with $1 \times 10^5$ MB49 WT or *C1GALT1C1* KO cells ($n=4$ mice per group). Tumours were harvested at humane endpoints, formalin-fixed, and paraffin-embedded. RNA was extracted from three 10-µm FFPE tissue sections per sample using the High Pure FFPE RNA Isolation Kit (Roche) according to the manufacturer's instructions.

#### Reverse transcription and quantitative PCR (RT-qPCR)

Total RNA was reverse-transcribed into complementary DNA (cDNA) using the High-Capacity cDNA Reverse Transcription Kit (Applied Biosystems) according to the manufacturer's instructions. Reverse transcription was performed at 37 °C for 120 min, followed by enzyme inactivation at 85 °C for 5 min. Expression of human and mouse cytokine genes (**Table M1**) was quantified using a 7500 Real-Time PCR System (Applied Biosystems). For the reaction, the Xpert Fast Probe kit (GRISP) was used. Amplification consisted of an initial denaturation at 94 °C for 1 min, followed by 40 cycles of 94 °C for 30 s, 64 °C for 1 min, and 68 °C for 4 min, with a final extension at 72 °C for 10 min. All samples were analyzed in duplicate, and mean Ct values were used to quantify relative gene expression using the $2^{-\Delta\Delta Ct}$ method. Gene expression levels were normalized to GAPDH expression.

**Table M1.** TaqMan gene expression assay.

| Human | Assay ID |
| --- | --- |
| *GAPDH* | Hs02786624_g1 |
| *CD44* | Hs01075864_m1 |
| *IL12A* | Hs01073447_m1 |
| *TGFBI* | Hs00998133_m1 |
| *Il10* | Hs00961622_m1 |
| *CCL2* | Hs00234140_m1 |
| *CXCL8* | Hs00174103_m1 |
| *IFNG* | Hs00989291_m1 |
| Mouse | Assay ID |
| *Gapdh* | Mm99999915_g1 |
| *Tgfb1* | Mm01178820_m1 |
| *Il12a* | Mm00434169_m1 |
| *Cd8a* | Mm01182107_g1 |
| *Il10* | Mm01288386_m1 |
| *Cd274* | Mm03048248_m1 |
| *Ifng* | Mm01168134_m1 |
| *Foxp3* | Mm00475162_m1 |

### Human monocyte isolation and differentiation into moDCs and macrophages

Human monocytes were isolated from buffy coats obtained from healthy blood donors at Hospital São João (Porto, Portugal). Briefly, peripheral blood mononuclear cells (PBMCs) were collected from previously centrifuged buffy coats (30 min, $1,200 \times g$, RT, no brake) and incubated with the RosetteSep Human Monocyte Enrichment Cocktail (StemCell Technologies), according to the manufacturer's instructions. The mixture was diluted 1:1 in PBS supplemented with 2% FBS, layered over Histopaque-1077 (Sigma-Aldrich), and centrifuged under the same conditions. The enriched monocyte layer was collected and washed three times with PBS. For moDC differentiation, $1 \times 10^6$ monocytes were seeded on 6-well plates and differentiated for 5 days in complete RPMI 1640 with GlutaMAX™ (10% FBS and 1% penicillin–streptomycin) supplemented with 50 ng/mL each of IL-4 and GM-CSF (ImmunoTools). For macrophage differentiation, the isolated monocytes were resuspended in complete RPMI supplemented with 50 ng/mL M-CSF (ImmunoTools) and seeded onto glass coverslips placed in 6-well plates at a density of $1.5 \times 10^6$ cells per well and cultured for 7 days at 37 °C in a humidified atmosphere with 5% CO<sub>2</sub>. Following the differentiation period, macrophages were polarised towards an M1-like phenotype by stimulation for 2 days with IFN-γ (25 ng/mL) (ImmunoTools) and LPS (10 ng/mL) (Sigma) or left unstimulated as M0-like controls.

### Co-culture of macrophages with glycoengineered cancer cells

After differentiation, unstimulated immature macrophages were co-cultured with glycoengineered T24 cells. The cancer cell models were added to the 6-well plates containing macrophages at a 1:3 (cancer cell: macrophage) ratio for 24 h at 37 °C in a humidified atmosphere with 5% CO<sub>2</sub>. For the reprogramming assay, M1-polarised macrophages were co-cultured under the same conditions for 72 h. Following this co-culture period, the conditioned media were collected for subsequent cytokine analysis, and the cells were detached by adding Accutase (Thermo Fisher Scientific) to the wells, followed by incubation for 20 min on ice. After centrifugation (7 min, $350 \times g$, 4 °C), the cells were resuspended in FACS buffer (PBS, 3% FBS, 0.1% sodium azide) and stained for flow cytometry. Macrophage phenotype was assessed using anti-human CD86, anti-human HLA-DR, anti-human CD14, anti-human CD206, anti-human CD163, and anti-human CD45 in FACS buffer. Cell viability was assessed by DAPI exclusion (750 ng/mL). Fluorescence minus one (FMO) controls for HLA-DR and CD86 were used. Samples were acquired on a Beckman Coulter Navios flow cytometer, and data were analysed using Kaluza C analysis software (version 2.1, Beckman Coulter) to discriminate distinct cell populations, determine the percentage of positive and negative cells, and calculate the mean fluorescence intensity (MFI).

##### **IL-10 quantification by ELISA**

IL-10 concentrations were measured in the conditioned media of all co-culture conditions using an ELISA according to the manufacturer's instructions (#430604, BioLegend). In brief, 100 μL of each undiluted sample was added to a 96-well plate pre-coated with a capture antibody. After incubation and subsequent washing, an enzyme-linked polyclonal detection antibody was added and incubated for a further 2 h. Finally, unbound antibody–enzyme complexes were removed by washing, and a substrate solution was added to develop a colour signal proportional to the amount of bound IL-10. Absorbance was measured using an iMark™ microplate reader (Bio-Rad), and IL-10 concentrations (pg/mL) were calculated from the standard curve. All conditions and standards were run in duplicate.

##### **Co-culture of moDCs with glycoengineered cancer cells**

T24 WT and glycoengineered cell models were seeded at $5 \times 10^4$ cells/well in 24-well plates and cultured at 37 °C. After 24 h, moDCs were added at a ratio of 1:5 (cancer cell:moDC) in adhesion buffer (20 mM Tris-HCl, 150 mM NaCl, 1 mM CaCl<sub>2</sub>, 1 mM MgCl<sub>2</sub> and 0.5% BSA; pH 8.0) to the tumour cell wells. LPS (50 ng mL<sup>-1</sup>; Sigma) was used as a positive control for DC activation. After 24 h of co-culture, conditioned media were collected for subsequent multiplex cytokine analysis, and moDCs were detached in ice-cold 1× PBS for 30 min. After centrifugation (5 min, 350× g, 4 °C), DCs were resuspended in FACS buffer and stained for flow cytometry. moDC differentiation and maturation were assessed by immunostaining with anti-human CD86 and anti-human CD11c, anti-human HLA-DR and anti-human CD45 in FACS buffer (**Table M2**). Cell viability was assessed by DAPI exclusion (750 ng mL<sup>-1</sup>). Samples were acquired on a Navios flow cytometer (Beckman Coulter). Flow cytometry data were analysed using Kaluza C analysis software (version 2.1, Beckman Coulter) to discriminate cell populations, determine the MFI and the percentage of positive cells.

##### **Multiplex cytokine and chemokine analysis**

Supernatants from moDCs co-cultured with glycoengineered T24 cells were collected after 24 h of co-culture and centrifuged (5 min, 3,000 × g, 4 °C) to remove cell debris. The Luminex Human Focused 15-plex-discovery assay (HDF-15) was then performed by Eve Technologies (Calgary, AB) to quantify the following cytokines/chemokines: GM-CSF, IFN-γ, IL-1β, IL-1Ra, IL-2, IL-4, IL-5, IL-6, IL-8, IL-10, IL-12p40, IL-12p70, MCP-1, TNF-α and IL-13.

#### **Tunicamycin-mediated inhibition of *N*-glycans**

T24 WT, *CIGALT1* KO and *CIGALT1* KO/*ST6GALNAC1* KI cells were transfected with CD44-targeting siRNA as described above. After 48 h of silencing, cells were treated with 0.5 μg/mL tunicamycin (T7765, Sigma-Aldrich) for 24 h to inhibit *N*-glycan biosynthesis, whereas control cells received an equivalent volume of DMSO. Experimental conditions comprised untreated cells, CD44-silenced cells, tunicamycin-treated cells, and cells subjected to both CD44 silencing and tunicamycin treatment. Cells were then detached using Versene solution (Thermo Fisher Scientific), counted, and co-cultured with monocyte-derived dendritic cells in adhesion buffer for 24 h. moDC viability and maturation were subsequently assessed by flow cytometry. Before the co-culture assay, tumour cells were stained with Concanavalin A (Con-A) and anti-human CD44 antibody and analysed by flow cytometry to confirm inhibition of *N*-glycan biosynthesis and CD44 expression.

#### **Stimulation of moDCs with tumour cell-conditioned medium**

To evaluate the effects of tumour-derived soluble factors on dendritic cell activation, conditioned medium (CM) from each glycoengineered T24 cell model was collected and used to stimulate moDCs. Briefly, T24 cell models were seeded at the same density used for the co-culture experiments ($1 \times 10^5$ cells per well). After 24 h, cell-free conditioned medium was collected and immediately used for moDC stimulation. moDCs ($5 \times 10^5$ cells per well) were seeded in duplicate in 24-well plates and cultured in a 1:1 mixture of adhesion buffer and tumour cell-conditioned medium. LPS (50 ng mL<sup>-1</sup>) was included as a positive control for dendritic cell activation. Following 24 h of incubation, moDCs were harvested and analysed by flow cytometry as described above. To determine whether the glycosylation profile of the tumour secretome reflected the glycoengineered phenotype of each T24 cell model, conditioned medium generated under reduced-serum conditions was analysed by immunoblotting. Reduced-serum medium minimised interference from abundant serum proteins, particularly albumin, thereby facilitating the detection of tumour-derived glycoproteins.

#### **Triple co-culture of moDCs, tumour cells, and T cells**

Monocyte-derived dendritic cells were first co-cultured with glycoengineered tumour cells as described above. After 24 h of co-culture, T cells were added directly to the existing cultures without removing the tumour cells or moDCs, and the triple co-cultures were incubated for an additional 24 h under the same culture conditions. Controls comprised T cells cultured with moDCs alone or tumour cells alone, together with the corresponding monocultures. At the end of the incubation period, cells were detached on ice, collected, and processed for flow cytometric analysis. Cells were stained with fluorochrome-conjugated antibodies against CD3, CD4, CD8, CD25, CD127 and CD69 (**Table M2**). DAPI was used as a viability dye to exclude dead cells. Samples were acquired using a Navios flow cytometer (Beckman Coulter), and data were analysed using Kaluza C analysis software (version 2.1, Beckman Coulter).

##### **Temporal proteomic analysis of moDCs**

Glycoengineered T24 tumour cells were co-cultured with moDCs for 6 or 24 h to characterise temporal proteomic changes associated with tumour cell-induced effects on moDC viability. moDCs collected immediately before co-culture (0 h) served as the baseline control. At each time point (0, 6 and 24 h), cells were harvested with ice-cold PBS. After co-culture, cells were detached on ice and moDCs were isolated by depleting tumour cells using CD326 (EpCAM) magnetic beads, washed with ice-cold PBS 1x, pelleted and stored at $-80^{\circ}\text{C}$. Whole protein extracts were extracted using RIPA lysis buffer (50 mM Tris-HCl (pH 8.0), 150 mM NaCl, 1% NP-40, 0.5% sodium deoxycholate, and 0.1% SDS in water) supplemented with protease inhibitors (Halt Protease inhibitor cocktail (100x)) and quantified using the Detergent Compatible (DC) Protein Assay (Bio-Rad). Equal amounts of protein (10 $\mu\text{g}$) were denatured in Laemmli sample buffer supplemented with 5% (v/v) β-mercaptoethanol, heated at $95^{\circ}\text{C}$ for 5 min, separated by SDS-PAGE and subjected to in-gel digestion before LC-MS/MS analysis.

##### **Proteomics receptor discovery using the GlycoAvatars platform**

Proteomics-based receptor discovery was performed using the GlycoAvatars platform. Briefly, GlycoAvatars were generated from plasma membrane extracts of T24 WT and T24 *C1GALT1* KO cell models as described by Miranda *et al*<sup>4</sup>. The generated GlycoAvatars were incubated with immature and mature monocyte-derived dendritic cells under the conditions described in the original GlycoAvatars methodology. After the interaction assay, bound membrane interactors were recovered and identified by LC-MS/MS as described previously.

##### **Affinity purification of CD44s-Tn-interacting proteins**

NHS-activated agarose beads (NHS Act Sepharose<sup>TM</sup> 4 fast flow, Cytiva) were equilibrated by washing once with 1 mM HCl to remove the storage solution, followed by two washes with coupling buffer. CD44s-Tn 30-mer peptide (DSPWITDSTDRI PATRDQDTFHPSGGSHTT)<sup>5</sup>, was diluted in coupling buffer and covalently immobilised onto the beads by incubation for 2 h at room temperature under end-over-end rotation. Non-glycosylated CD44 30-mer peptide was used as a negative binding control. Remaining active NHS ester groups were quenched using quenching buffer for 30 min at room temperature. Beads were then washed alternately with the manufacturer's recommended wash buffers to remove non-covalently bound peptide and equilibrated in lysis buffer. Plasma membrane extracts pooled from moDCs from three donors (150 $\mu\text{g}$) were incubated overnight at $4^{\circ}\text{C}$ with peptide-coupled beads with continuous rotation. After incubation, unbound material was removed, and the beads were washed with stringency buffer to reduce nonspecific interactions, then washed again with a low-salt,

detergent-free buffer. Bound proteins were eluted by three sequential incubations with acidic elution buffer, and the combined eluate was precipitated with six volumes of cold acetone. Protein pellets were resuspended in Laemmli sample buffer supplemented with β-mercaptoethanol, separated by SDS-PAGE, and subjected to in-gel tryptic digestion for LC-MS/MS analysis.

#### **MRC1 affinity purification and proteomics**

Plasma membrane extracts were prepared from T24 WT, *CIGALT1* KO, and *CIGALT1* KO/*ST6GALNAC1* KI cells. An aliquot of the *CIGALT1* KO/*ST6GALNAC1* KI plasma membrane extract was treated with PNGase F to remove *N*-glycans, generating paired untreated and *N*-glycan-depleted protein samples. For *N*-glycan removal, the plasma membrane extract was supplemented with NP-40 (2 μL of 10% NP-40 per 20 μL of sample) to neutralise SDS before incubation with PNGase F for 4 h at 37 °C. Following digestion, free *N*-glycans were removed by centrifugal filtration using 10-kDa Amicon Ultra filters. Untreated samples were processed through the same filtration procedure to ensure identical sample handling. For affinity purification, 300 μg of plasma membrane extract was incubated with 4 μg of biotinylated MRC1 in TBS containing 0.1% SDS to maintain membrane protein solubilisation, supplemented with 2 mM $\text{CaCl}_2$ and 2 mM $\text{MgCl}_2$ because C-type lectin carbohydrate recognition is calcium-dependent. Samples were incubated for 2 h at 4 °C with gentle end-over-end mixing. Streptavidin-coated magnetic beads were then added and incubated for a further 60 min under the same conditions to capture MRC1-bound protein complexes. Beads were collected using a magnetic rack and washed three times with 500 μL of TBS containing 0.1% SDS, 2 mM $\text{CaCl}_2$ and 2 mM $\text{MgCl}_2$ to remove nonspecifically bound proteins. Bound proteins were eluted by incubation in Laemmli sample buffer supplemented with β-mercaptoethanol for 30 min at 70 °C. Eluted proteins were resolved by SDS-PAGE and visualised by reversible zinc staining. Each lane was excised and subjected to in-gel tryptic digestion. The resulting peptides were analysed by LC-MS/MS.

#### **In-gel digestion**

Protein samples were separated by SDS-PAGE and stained with PageBlue Protein Staining Solution or reversible zinc stain. Gel lanes were manually excised, cut into approximately 3 mm<sup>3</sup> pieces and subjected to in-gel tryptic digestion. Briefly, gel pieces were destained using 50 mM ammonium bicarbonate in 50% acetonitrile (for PageBlue protein-stained gels) or glycine-HCl (pH 2) (for zinc-stained gels), washed three times with Milli-Q water, dehydrated with acetonitrile, reduced with 10 mM dithiothreitol (DTT) for 30 min at 56 °C and alkylated with 55 mM iodoacetamide (IAA) for 20 min at room temperature in the dark, and dehydrated again with acetonitrile. Proteins were digested overnight at 37 °C using sequencing-grade modified trypsin (for temporal proteomic analysis of co-cultured moDCs, GlycoAvatars assay, and MRC1-Biotin pulldown; Promega) or sequencing-grade chymotrypsin (for CD44s-Tn pull-down; Promega). Peptides were extracted with 50% acetonitrile containing 2% formic acid, dried by vacuum centrifugation and resuspended in 0.1% formic acid before LC-MS/MS analysis.

#### **LC-MS/MS acquisition**

For protein identification, peptides were loaded into a Vanquish Neo UHPLC system coupled with a Q Exactive Plus Hybrid Quadrupole-Orbitrap mass spectrometer (Thermo Fisher Scientific), fitted with a nano-electrospray ion source (EASY-Spray source; Thermo Fisher Scientific). The mobile phases were 0.1% FA in ultrapure water (eluent A) and 0.1% FA in 80% ACN (eluent B). A 10 μL sample was injected into a trapping column (C18 PepMap Neo, 5 μm particle size; 300 μm × 5 mm) and separated on an analytical column (EASY-Spray C18 PepMap, 100 $\text{\AA}$, 75 μm ∼ 150 mm, 3 μm particle size) at 0.25 μL min<sup>-1</sup> and 35 $^{\circ}$C. Peptides were separated using a gradient of 2.5%–12% B over 7 min, 12%–46% B over 50 min, and 46%–99% B over 5 min, holding at 99% B for 10 min. The mass spectrometer was operated in positive-ion mode (m/z 300–2000) with a spray voltage of 1.9 kV and a capillary temperature of 275 $^{\circ}$C. Full-scan MS settings comprised a resolution of 140,000, AGC target of $3 \times 10^6$, and 200 ms maximum injection time, with the 15 most intense ions selected for higher-energy collisional dissociation (HCD) at a normalised collision energy of 30%. Data were recorded with Xcalibur (v4.5).

#### **Protein identification and quantification**

Temporal proteomic analysis of co-cultured moDCs, GlycoAvatars assay, CD44s-Tn pull-down, and MRC1-Biotin pull-down raw files were processed in Proteome Discoverer 3.1 (Thermo Fisher Scientific) using the SequestHT search engine and Percolator against the Swiss-Prot human proteome database (UP000005640 fasta downloaded on 21/11/2024). Search parameters included trypsin or chymotrypsin specificity with up to two missed cleavages, a 10-ppm precursor-ion tolerance and a 0.02-Da fragment-ion tolerance, carbamidomethylation of cysteine as a fixed modification (+57.021 Da) and methionine oxidation as a variable modification (+15.995 Da). Protein groups supported by fewer than two unique peptides or that did not meet high-confidence FDR thresholds were excluded from downstream analyses. Gene Ontology (GO) annotation for cellular component, molecular function and biological process was performed in Proteome Discoverer 3.1. Protein abundance was quantified using label-free MS1 intensity-derived precursor areas. Differential protein abundance was expressed as log<sub>2</sub> fold change between the relevant experimental conditions for each comparison. Statistical significance was assessed using moderated t-tests with FDR correction.

#### **Proteomic data analysis**

Proteins were considered significantly enriched when log<sub>2</sub> fold change > 1.5, adjusted P < 0.05 and detected in two biological replicates for GlycoAvatars and CD44s-Tn pull-down assays (**Fig. 4**) and three biological replicates for temporal proteomics analysis of co-cultured moDCs (**Fig. 3**).

Downstream statistical and bioinformatic proteomic analyses were performed in R (RStudio version 4.6.0). Protein abundance data were normalised prior to statistical analysis, and differential protein abundance between experimental conditions was calculated as log<sub>2</sub> fold change. Principal component analysis (PCA), z-score normalisation, data visualisation, and statistical analyses were performed using the tidyverse, limma, FactoMineR, factoextra, and ggplot2 packages. Venn diagrams were generated using ggVennDiagram (or VennDiagram), and Gene Ontology enrichment analyses were performed using clusterProfiler. Protein abundance plots were generated using ggplot2.

#### **Western blot**

Total protein extracts from MB49, T24, AGS and SW480 were separated on a 4–20% precast polyacrylamide gel (Bio-Rad) and transferred to nitrocellulose membranes (GE Healthcare Life Sciences). CD44 was detected using an anti-CD44 antibody (1:5,000; ab157107, Abcam) for 1 h at room temperature. Secondary detection was performed using goat anti-rabbit IgG (H+L) horseradish peroxidase (HRP)-conjugated antibody (1:60,000; Thermo Fisher Scientific) for 30 min at room temperature. The Amersham ECL Prime Western Blotting Detection Reagent (GE Healthcare Life Sciences) was used for chemiluminescent detection. Tn and sTn expression in conditioned medium from T24 WT, T24 KO and T24 KO/KI cells was also evaluated by immunoblotting using VVA lectin (1:5,000; VectorLabs) and B72.3 antibody (1 µg/mL, ab199002, Abcam), respectively. After 1 h of incubation at room temperature, VECTASTAIN Elite ABC-HRP (1:10; Vector Laboratories) and peroxidase affiniPure goat anti-mouse IgG (H+L) polyclonal antibody (1:70,000; ImmunoResearch) were used for VVA and B72.3 analysis, respectively.

To evaluate gasdermin D activation, dendritic cells were recovered after co-culture with tumour cells, lysed, and total protein extracts were subjected to immunoblot analysis as described above. Gasdermin D cleavage was assessed using an anti-cleaved *N*-terminal GSDMD antibody ([EPR20829-408] (ab215203)) followed by incubation with Goat Anti-Rabbit IgG H&L (HRP) antibody (ab6721). Immunoreactive bands were visualised using Amersham ECL Prime Western Blotting Detection Reagent (GE Healthcare Life Sciences). Densitometric analysis was performed using Image Lab software (Bio-Rad) on a ChemiDoc XRS imaging system (Bio-Rad).

#### **Co-culture, internalisation and blocking assays**

Glycoengineered T24 tumour cells were co-cultured with immature or mature moDCs at a tumour cell-to-moDC ratio of 1:5 for 24 h. Following 24 h co-culture, moDCs were recovered and analysed by flow cytometry. For VVA blocking assays, T24 cell models were detached using Versene solution (Thermo Fisher Scientific) and incubated with unconjugated VVA (0.01 and 0.001 µg) for 1 h at 4 °C. Cells were subsequently washed to remove unbound lectin and co-cultured with immature or mature moDCs for 24 h. For MRC1 blocking assays, T24 tumour cells were seeded 24 h before the co-culture experiments. Immature or mature moDCs were incubated with anti-MRC1 blocking antibody (Purified anti-human

CD206 (MMR), Clone 15-2, 0.5 mg mL<sup>-1</sup>, BioLegend) at 1:20, 1:100 and 1:500 dilutions and co-cultured with T24 cell models for 24 h, in the same co-culture conditions as previously described for moDCs. To assess CD206 internalisation, moDCs were co-cultured with glycoengineered T24 cell models for 4 h. Cells were then stained for surface CD206 (MRC1) without permeabilisation, and surface CD206 expression was quantified by flow cytometry. Following co-culture incubations, moDCs were detached using cold PBS 1x, incubated on ice for 15 min to allow cell detachment, followed by immunostaining and analysis by flow cytometry.

##### ***In vitro* flow cytometry assays**

Cells were detached using Accutase™ Cell Detachment Solution (BD™), washed with PBS and subsequently fixed with 2% paraformaldehyde (PFA; Sigma-Aldrich). For the T24, AGS and SW480 cell models, cells were incubated for 1 h at room temperature (RT) with VVA, PNA and the monoclonal antibody B72.3+CC49 (2 µg per 1 × 10<sup>6</sup> cells). For the MB49 WT and *CIGALTIC1* KO cell models, cells were incubated using VVA and PNA in the presence or absence of 70 mU neuraminidase (NeuAse) in sodium acetate buffer (pH 5.5) overnight. The T24 cell models were stained with biotinylated MRC1 for 1 h at 4 °C, in HBSS 1x medium supplemented with 0.5% BSA, followed by incubation with Alexa Fluor 488-streptavidin at a 1:300 dilution for 30 min at RT. Immature and mature dendritic cells were stained for the surface expression of MRC1 (CD206), CD301 (MGL) and Siglec-15 (CD33L3) using fluorochrome-conjugated antibodies (**Table M2**). Samples were acquired on a Navios flow cytometer (Beckman Coulter), and data were analysed using Kaluza C analysis software (version 2.1, Beckman Coulter).

**Table M2.** Conventional Human flow cytometry panels.

| Antibodies | Clone | Source | Dilution |
| --- | --- | --- | --- |
| Anti-Human |  |  |  |
| **Macrophage panel** |  |  |  |
| CD86 FITC | BU63 | Immunotools | 1:25 |
| HLA-DR PE | HI43 | Immunotools | 1:25 |
| CD14 PerCP5.5 | HCD14 | BioLegend | 1:50 |
| CD45 PE-Cy7 | 2D1 | BioLegend | 1:100 |
| CD163 APC | RM3/1 | BioLegend | 1:100 |
| CD206 APC-H7 | 15-2 | BioLegend | 1:100 |
| **Dendritic cell panel** |  |  |  |
| CD86 FITC | BU63 | Immunotools | 1:25 |
| HLA-DR PE | HI43 | Immunotools | 1:25 |
| CD45 PE-Cy7 | 2D1 | BioLegend | 1:100 |
| CD11c APC | BU15 | Immunotools | 1:50 |
| **T-cell panel** |  |  |  |
| CD3 APC-H7 | SK7 | BD Bioscience | 1:50 |
| CD4 PB | RPA-T4 | Beckman Coulter | 1:50 |
| CD8 PerCP5.5 | SK1 | BD Bioscience | 1:100 |
| CD25 PE | 2A3 | BD Bioscience | 1:50 |
| CD127 PE-Cy7 | HIL-7R-M21 | BD Bioscience | 1:100 |
| CD69 APC | FN50 | BD Bioscience | 1:50 |
| **Glycans/receptor detection** |  |  |  |
| VVA FITC |  | Vector Laboratories | 1:1000 |
| PNA FITC |  | Vector Laboratories | 1:1000 |
| TAG-72 | B72.3+CC49 | Thermo Fisher Scientific | 1:10 |
| Concanavalin A biotin |  | Vector Laboratories | 1:2000 |
| CD44 VF450 | IM7 | Thermo Fisher Scientific | 1:100 |
| CD206 (MRC1) biotin | CD6-H52H9 | Acro Biosystems | 1:200 |
| CD301 (CLEC10A) APC | H037G3 | BioLegend | 1:50 |
| CD206 PE | 15-2 | BioLegend | 1:100 |
| Siglec-15 AF488 |  | R&D systems | 1:50 |

### scRNA-seq analysis

Publicly available single-cell RNA-sequencing (scRNA-seq) data from eight muscle-invasive bladder cancer (MIBC) and three healthy bladder tissues generated by Chen *et al.*<sup>6</sup> were obtained from GSA-Human (accession HRA000212) and reanalysed using Seurat (v5.3.0) in R Studio (v4.0). Following quality control, normalisation and dimensionality reduction, cells were clustered using the Seurat workflow and manually annotated based on the expression of canonical lineage markers, including *EPCAM* (epithelial cells), *LYZ* (myeloid cells), *CD3E* (T cells), *CD79A* (B-cells), *ACTA2* (myofibroblasts), *COL1A1* (fibroblasts) and *HSPG2* (endothelial cells). The myeloid compartment was subsequently reclustered (resolution = 0.7) and manually annotated according to the expression of established lineage markers, identifying tumour-associated macrophages (TAMs), monocytes, CLEC9A<sup>+</sup> dendritic cells (DCs), CD1C<sup>+</sup> DCs and epithelial and endothelial doublets. MIBC samples were stratified by expression of the Ensembl transcript [ENSG00000026508], which encodes CD44. Based on median transcript expression, tumours were classified as CD44-positive or CD44-negative, while healthy bladder samples served as non-malignant controls. Cell composition and differential gene expression analyses were subsequently performed across the identified immune cell populations. Violin plots, dot plots and dimensionality reduction plots were generated using Seurat, and Heatmaps were generated using pheatmap (version 1.0.13).

### **Animal ethics statement**

Animal experiments were conducted in accordance with the European Union Directive 2010/63/EU and Portuguese national legislation (Decreto-Lei n.º 113/2013 de 7 de Agosto). Two independent *in vivo* experiments were conducted at two animal facilities bred under specific pathogen-free conditions at the CIPHE (Marseille, France) animal facilities (agreement number: B1301407), and C57BL/6J mice were purchased from Janvier Laboratories. The second experiment was conducted at the i3S-UP animal facility (Porto, Portugal) in accordance with institutional and European animal-care guidelines. The C57BL/6J mice were purchased from Charles River. The following humane endpoints were predefined: any signs of distress, suffering or pain, body-weight loss of >20%, dehydration, anorexia, anuria or a moribund state, whether or not related to the experimental procedure.

### **Generation of bone-marrow-derived dendritic cells**

Bone marrow cells were isolated from the femurs and tibias of healthy C57BL/6J mice. Bones were flushed with sterile supplemented RPMI 1640 medium using a 25-gauge needle, and the resulting cell suspension was passed through a 70-µm cell strainer. Red blood cells were lysed using ACK lysis buffer, and the remaining cells were washed and cultured in complete RPMI 1640 supplemented with 10% FBS, 2 mM L-glutamine, 1% penicillin/streptomycin (PenStrep) (10,000 units/mL penicillin; 10,000 µg/mL streptomycin), 50 µM β-mercaptoethanol, recombinant murine granulocyte-macrophage colony-stimulating factor (GM-CSF; 20 ng ml<sup>-1</sup>) and murine IL-4 (20 ng ml<sup>-1</sup>). Cells were maintained at 37 °C in a humidified atmosphere containing 5% CO<sub>2</sub>. On day 5, 500 µl of fresh complete medium containing GM-CSF and IL-4 was added. Loosely adherent and non-adherent cells were harvested on day 7 and used as bone marrow-derived dendritic cells (BMDCs) for subsequent experiments.

### **Co-culture of BMDCs with MB49 cells**

Wild-type or *C1GALT1C1* KO MB49 cells were seeded in tissue culture plates and allowed to adhere overnight. BMDCs were then added at a tumour cell: BMDC ratio of 1:5 and co-cultured for 24 h in complete RPMI 1640 medium at 37 °C in a humidified atmosphere containing 5% CO<sub>2</sub>. Following co-culture, cells were collected, washed with FACS buffer and analysed by flow cytometry for the expression of dendritic cell activation and maturation markers and cell viability using a DxFLEx flow cytometer (Beckman Coulter).

### **Syngeneic MB49 murine bladder cancer model**

Seven-week-old female C57BL/6J mice were subcutaneously inoculated in the right flank with either $1 \times 10^5$ or $1 \times 10^6$ MB49 WT or *C1GALT1C1* KO cells resuspended in a 1:1 mixture of Matrigel and serum-free RPMI. Tumour growth was monitored by calliper measurements every 2–3 days, and tumour volumes were calculated using the formula (length × width<sup>2</sup>)/2. Mice were euthanised when tumours reached a volume of approximately 1,500 mm<sup>3</sup> or when predefined humane endpoints were met. Tumours, tumour-draining lymph nodes and spleens were collected for downstream immune profiling. Tumour-infiltrating leukocytes (TILs) were isolated for mass cytometry (CyTOF) analysis, whereas immune cell populations from spleens and lymph nodes were analysed by conventional flow cytometry.

##### **Tumour dissociation and mass cytometry staining**

Tumours (n=6 MB49 WT, n=6 MB49 KO) were harvested, mechanically and enzymatically dissociated using a GentleMACS Octo Dissociator (Miltenyi Biotec) together with the Tumour Dissociation Enzymatic Kit (Miltenyi Biotec, #130-096-730), after which the cell suspension was filtered, and enzymatic activity was stopped with PBS/EDTA (0.1 M). Cells were then washed in HBSS/2% FCS, and Fc receptors were blocked with 24G2 hybridoma (10 min, 4 °C) prior to CD45<sup>+</sup> enrichment using CD45 MicroBeads (Miltenyi Biotec, #130-052-301) according to the manufacturer's protocol on a MultiMACS Cell24 separator (Miltenyi Biotec). Enrichment efficiency and viability were subsequently confirmed by flow cytometry (Attune NxT) using CD45.2-APC, Ter119-PE, and Sytox Green staining. For mass cytometry, cells were stained with 100 µL of CisPt 198 (Standard Biotoools, #201198) for dead cell exclusion and transferred to a Curiox Laminar Wash™ MINI 1000 to limit cell loss during the subsequent barcoding and washing steps. CD45 barcoding was then performed using a combination of three metal-tagged CD45.2/CD45 antibodies (30 min, 4 °C, maximum volume 80 µL), followed by a Curiox wash in CSB (Standard Biotoools #201068) (9 washes, 80 µL, flow 5) to remove unbound barcodes, after which samples were pooled by treatment group. Pooled cells were next stained with an anti-chemokine receptor mix (MERTK, CD192, CX3CR1; 30 min, RT), washed again, and then incubated with the remaining surface antibody panel (30 min, 4 °C). After transfer to a V-bottom plate, cells underwent fixation, permeabilization, and intracellular staining using the Foxp3/Transcription Factor Staining Buffer Set (Thermo Fisher Scientific, #00-5523-00), followed by overnight DNA staining at 4 °C with Cell-ID Intercalator Ir (201192B, Standard Biotoools), diluted 1:1000 in Cytotfix/Cytoperm (BD 51-2090KZ). The following day, cells were washed twice in CSB and stored at -80 °C in FCS containing 10% DMSO until acquisition. Prior to acquisition, cells were washed and diluted to 5 × 10<sup>5</sup> cells/mL in H<sub>2</sub>O containing 10% EQ Four Element Calibration Beads (Standard Biotoools, #201078). Flow cytometry standard (FCS) files were normalised to the bead signal using the Standard Biotoools algorithm; beads were excluded based on 165Ho versus 153Eu signal, and singlet events were defined using Gaussian parameters, event length, DNA channels (191Ir/193Ir), and 198Pt-negative staining in Cytobank software (Beckman Coulter).

##### **Debarcoding and multidimensional data analysis**

The data were debarcoded using the custom R Shiny interface, 'Vaevictis' (available at https://github.com/stuchly/vaevictis), to generate debarcoding dimensions (**Supplementary Fig. S23**). Manual debarcoding was performed in Cytobank using the Vaevictis dimensions. The resulting debarcoded files were further analysed using either opt-SNE (perplexity = 30, iterations = 1000) or Rphenograph through ‘CIPHEBank’, an in-house R Shiny interface designed for classifying and visualising tumour-infiltrating lymphocyte (TIL) subpopulations based on both cell surface and intracellular markers (**Table M3**). Rphenograph used the Euclidean distance to identify k-nearest neighbours (k = 30) and the Jaccard coefficient to calculate similarities. The Louvain algorithm was then applied to partition the network and optimise modularity, yielding 21 metaclusters (**Supplementary Fig. S24**). Expert-guided manual annotation was performed using the median expression of marker intensities for each cluster. Heatmaps and hierarchical clustering were created with Morpheus (https://software.broadinstitute.org/morpheus). The cluster data were visualised in Cytobank to produce density plots and cluster overlays.

**Table M3.** Mass cytometry (CyTOF) antibodies.

| Marker | Metal | Clone | Compartment | Dilution | Reference | Source |
| --- | --- | --- | --- | --- | --- | --- |
| CD45.2 | 111 Cd | 104 | barcoding |  | BLE109802 | Biolegend |
| CD45.2 | 112 Cd | 104 | barcoding |  | BLE109802 | Biolegend |
| CD45.2 | 114 Cd | 104 | barcoding |  | BLE109802 | Biolegend |
| CD45 | 195Pt | 30F11 | barcoding |  | 92J006195 | Standard Biotools |
| CD45 | 89Y | 30F11 | barcoding |  | 3089005C | Standard Biotools |
| CD11b | 115In | M1/70 | Surface | 200 | BLE101202 | Biolegend |
| CD11c | 172Yb | N418 | Surface | 200 | BLE117341 | Biolegend |
| Ly6G | 141Pr | 1A8 | Surface | 200 | 3141008B | Standard Biotools |
| CD39 | 142Nd | 24DMS1 | Surface | 100 | 3142005B | Standard Biotools |
| CD274 | 144Nd | 10F.9G2 | Surface | 25 | BLE124302 | Biolegend |
| CD4 | 145Nd | RM4-5 | Surface | 100 | 3145002B | Biolegend |
| CD8a | 146Nd | 53-6.7 | Surface | 600 | BLE100746 | Biolegend |
| IRF8 | 147Sm | V3GYWCH | Intracellular | 600 | 53-9852-82 | invitrogen |
| CX3CR1 | 148Nd | SA011F11 | Surface RT | 1600 | BLE149002 | Biolegend |
| IRF4 | 149Sm | IRF4.3E4 | Intracellular | 800 | BLE646402 | Biolegend |
| Ly6C | 150Nd | HK1.4 | Surface | 100 | BLE128002 | Biolegend |
| CD64 | 151Eu | X54-5/7.1 | Surface | 100 | 3151012B | Standard Biotools |
| CD25 | 151Eu | 3C7 | Surface | 100 | 3151007B | Standard Biotools |
| LysM | 152Sm | MA1-82873 | Intracellular | 25 | BGN-0696-5B1 | thermo |
| CD152 | 154Sm | UC10-4B9 | Intracellular | 100 | 3154008B | Standard Biotools |
| CD279 | 155Gd | 29F.1A12 | Surface | 200 | 135202 | Biolegend |
| CD192 | 156Gd | SA203G11 | Surface RT | 100 | 93501 | Biolegend |
| CD68 | 158Gd | FA-11 | Intracellular | 100 | 14-0681-82 | Biolegend |
| CD73 | 159Tb | TY11.8 | Surface | 400 | BLE127202 | Biolegend |
| FoxP3 | 161Dy | MF14 | Intracellular | 50 | BLE126402 | Biolegend |
| TNFα | 162Dy | MP6-XT22 | Intracellular | 100 | 3162002B | Standard Biotools |
| Klrg1 | 163Dy | 2F1 | Surface | 800 | 16-5893-85 | eBioscience |
| CD172a | 164Dy | P84 | Surface | 50 | 51-410-28671 | BD Bioscience |
| MER-TK | 165Ho | REA477 | Surface RT | 30 µL | 130-107-477 | Miltenyi |
| CD26 | 166Er | H194-112 | Surface | 200 | BLE137801 | Biolegend |
| F4/80 | 167Er | BM8 | Surface | 400 | 123102 | Biolegend |
| XCR1 | 168Er | ZET | Surface | 1600 | 148202 | Biolegend |
| CD206 | 169Tm | C068C2 | Surface | 100 | 3169021B | Standard Biotools |
| CD161 | 170Er | PK136 | Surface | 100 | 3170002B | Standard Biotools |
| CD44 | 171Yb | IM7 | Surface | 100 | 3171003B | Standard Biotools |
| CD317 | 173Yb | 129c1 | Surface | 25 | BLE127102 | Biolegend |
| Gata3 | 175Lu | 16E10A23 | Intracellular | 25 | BLE653802 | Biolegend |
| CD62L | 160Gd | MEL-14 | Surface | 100 | 3160008B | Standard Biotools |
| CD3e | 176 Yb | 145-2C11 | Surface | 100 | BLE100302 | Biolegend |
| IA IE | 209Bi | M5/114.15.2 | Surface | 200 | 3209006B | Standard Biotools |
| CD38 | 153Eu | 90 | Surface | 400 | BLE102702 | Biolegend |
| CD88 | 174Yb | 20.7 | Surface | 400 | BLE135802 | Biolegend |
| Eomes | 143Nd | 7C9B03 | Intracellular | 100 | 662002 | Biolegend |
| Iba1 | 113In | E4O4W | Intracellular | 400 | 17198B | Cell signaling |

##### Anti-MRC1 treatment in the MB49 *CIGALTIC1* KO syngeneic murine model

Seven-week-old female C57BL/6J mice were subcutaneously inoculated in the flank with $1 \times 10^4$ MB49 *CIGALTIC1* KO cells. Once tumours reached a volume of 75–100 mm<sup>3</sup>, mice were randomly allocated to receive 2 µg or 20 µg of murine anti-mouse MRC1 (CD206) monoclonal antibody by peritumoural injection. Control mice received an equivalent volume of PBS. Treatments were administered three times at 2–3-day intervals. Calliper measurements were used to monitor tumour growth throughout the study. At the experimental endpoint, mice were euthanised, and tumours and tumour-draining lymph nodes were collected for immune profiling by conventional flow cytometry.

##### Conventional flow cytometry *in vivo* studies

Single-cell suspensions ($1.5 \times 10^6$ cells) were washed in FACS buffer and incubated with Fc receptor-blocking reagent for 15 min at 4 °C to minimise non-specific antibody binding. Cells were then stained with the antibody panel listed in **Table M4** for 45 min at 4 °C in the dark. Following washing, cells were acquired on a DxFLEX flow cytometer (Beckman Coulter). Compensation was performed using single-stained controls, and FMO controls were used to define gates for CD44, CD62L, CD73 and I-A/I-E. Data were analysed using Kaluza software (version 1.7, Cytognos SL, Salamanca, Spain). Cell populations were identified according to the gating strategy shown in **Supplementary Fig. S25** and quantified as the percentage of positive cells or MFI, as indicated.

**Table M4.** Lymphoid and myeloid murine panels.

| Antibodies | Clone | Source | Dilution |
| --- | --- | --- | --- |
| Anti-Mouse |  |  |  |
| **Myeloid panel** |  |  |  |
| CD19 PB | 1D3 | BD Bioscience | 1:50 |
| CD64 V500 | X54-5/7.1 | BioLegend | 1:50 |
| F4/80 FITC | BM8 | Thermo Fisher Scientific | 1:20 |
| I-A/I-E PE | M5/114.15.2 | BD Bioscience | 1:100 |
| CD45 PerCP5.5 | 30-F11 | BD Bioscience | 1:100 |
| CD11c PE-Cy7 | N418 | BioLegend | 1:50 |
| CD206 APC | C068C2 | BioLegend | 1:100 |
| XCR1 APC-H7 | ZET | BioLegend | 1:50 |
| Ly-6G APC-H7 | 1A8 | BD Bioscience | 1:100 |
| **Lymphoid panel** |  |  |  |
| CD44 V450 | IM7 | BioLegend | 1:50 |
| CD3 eBioscience500 | 17A2 | Thermo Fisher Scientific | 1:50 |
| CD62L FITC | MEL-14 | BD Bioscience | 1:50 |
| CD19 PE | 1D3 | BD Bioscience | 1:50 |
| CD45 PerCP5.5 | 30-F11 | BD Bioscience | 1:100 |
| CD8 PE-Cy7 | 53-6.7 | BD Bioscience | 1:50 |
| CD73 APC | TY/11.8 | BD Bioscience | 1:50 |
| CD4 APC-H7 | GK1.5 | BD Bioscience | 1:50 |

**Randomisation and statistical analysis**

No statistical methods were used to predetermine sample size. Animals were randomly assigned to experimental groups. Statistical analyses were performed using GraphPad Prism (version 10.1.1, GraphPad Software). Data are presented as mean ± SD, unless otherwise indicated. Box plots show the median, interquartile range (IQR) and range. Biological replicates represent independent donors for human cell experiments, independent mice for *in vivo* experiments or independent experiments, as specified in the figure legends. Data distribution was assessed before selecting statistical tests. For experiments using cells derived from the same donor across multiple conditions, paired statistical analyses were performed to account for donor-to-donor variability. Exact *P* values are reported in the figures, and $P < 0.05$ was considered statistically significant.
